# A systematic evaluation of dynamic functional connectivity methods using simulation data

**DOI:** 10.1101/2024.07.09.600728

**Authors:** Binke Yuan, Junjie Yang, Xiaolin Guo, Xiaowei Gao, Zhe Hu, Junjing Li, Jiaxuan Liu, Yaling Wang, Zhiheng Qu, Wanchun Li, Zhongqi Li, Wanjing Li, Yien Huang, Jiali Chen, Hao Wen, Junle Li, Dongqiang Liu, Hui Xie

**Affiliations:** Philosophy and Social Science Laboratory of Reading and Development in Children and Adolescents (South China Normal University), Ministry of Education, China; Key Laboratory of Brain, Cognition and Education Sciences, Ministry of Education, China: Institute for Brain Research and Rehabilitation, South China Normal University, Guangzhou, China; Research Center of Brain and Cognitive Neuroscience, Liaoning Normal University, Dalian, China; Key Laboratory of Brain and Cognitive Neuroscience, Liaoning Province, Dalian, PR China

**Keywords:** Dynamic functional connectivity, simulation data, temporal reoccurring state, sojourn time distribution, mean squared error, spatial and temporal correlation

## Abstract

Numerous dynamic functional connectivity (dFC) methods have been proposed to study time-resolved network reorganization in rest and task fMRI. However, a comprehensive comparison of their performance is lacking. In this study, we compared the efficacy of seven dFC methods (and their enhanced versions) to track transient network reconfiguration using simulation data. The seven methods include flexible least squares (FLS), dynamic conditional correlation (DCC), general linear Kalman filter (GLKF), multiplication of temporal derivatives (MTD), sliding-window functional connectivity with L1-regularization (SWFC), hidden Markov models (HMM), and hidden semi-Markov models (HSMM). Multiple datasets of non-fMRI-BOLD and fMRI-BOLD signals with predefined covariance structures, signal-to-noise ratio levels, and sojourn time distributions were simulated. We adopted inter-subject analysis to eliminate the effects of signals of non-interest, resulting in enhanced methods: ISSWFC, ISMTD, ISDCC, ISFLS, ISKF, ISHMM, and ISHSMM. Efficacy was defined as the spatiotemporal association between simulated and estimated data. We found that all enhanced dFC methods outperformed their original versions. Efficacies depend on several factors, such as considering the neurovascular effect in simulated data, the covariance structure between two time series, state sojourn distribution, and signal-to-noise ratio levels. These results highlight the importance of selecting appropriate dFC methods in fMRI study.

## Introduction

Continuous recording of Blood Oxygen Level Dependent (BOLD) signals using functional MRI (fMRI) enables the investigation of brain network reorganization on a timescale of seconds and its relationship with cognition, behavior, and clinical status (Cohen, 2018; Hutchison et al., 2013). Network reorganization or dynamic functional connectivity (dFC) can be estimated frame by frame or within a short window (∼ 20– 30 time points). Numerous dFC methods have been proposed, such as the sliding-window functional connectivity with L1-regularization (SWFC) (Allen et al., 2014; Shakil et al., 2016), dynamic conditional correlation (DCC) (Choe et al., 2017; Lindquist et al., 2014; Yuan et al., 2023a; Yuan et al., 2023b), Multiplication of Temporal Derivatives (MTD) (Shine et al., 2015), Flexible Least Squares (FLS) (Kalaba and Tesfatsion, 1989; Liao et al., 2014), general linear Kalman filter (Kang et al., 2011; Milde et al., 2010), Hidden Markov models (HMM) (Chen et al., 2022; Eavani et al., 2013; Vidaurre et al., 2017), and Hidden semi-Markov models (HSMM) (Shappell et al., 2019). Although these methods have demonstrated their accuracy, reliability, and potential caveats, a comprehensive comparison regarding the efficacy of these methods is lacking.

The efficacy of dFC methods is generally regarded as its ability to unveil the underlying spatiotemporal covariance structures of signals of interest (Calhoun et al., 2014; Cohen, 2018; Xie et al., 2019). Two types of factors are related to the efficacy of dFC methods: the implicit and external factors. The implicit factors mainly refer to the underlying mathematical assumptions and formulations, such as the mathematical transform function applied to the raw data before the computation of covariance (e.g., GARCH(1,1) model of DCC), the general relation function used to calculate the covariance structure (e.g., Pearson correlation for SWFC), and the constraint method for weighting the signals of interest (e.g., Gaussian tapered window used for SWFC) (Thompson and Fransson, 2018). The free parameters in each step will affect the efficacy. For example, as the most widely used method, SWFC results inherently dependent (Shakil et al., 2016; Thompson et al., 2018) on the number of samples per window (i.e., the window length), the overlap between two consecutive windows, the way to deal with the data at the beginning and end of the window (rectangular window vs Gaussian tapered window), and L1 regularization (Allen et al., 2014; Xie et al., 2019). For HMM, the sojourn time distribution of a dFC state is implicitly geometrical (Shappell et al., 2019; Yu, 2010). Therefore, HMM is inappropriate in data with other sojourn time distributions, such as Poisson distribution (Shappell et al., 2019).

The external factors include the preprocessing steps of fMRI-BOLD data (Hutchison et al., 2013), the data types (e.g., simulated vs real fMRI-BOLD data, simulated non-fMRI-BOLD vs simulated fMRI-BOLD data), the benchmark of efficacy (e.g., association with spatiotemporal covariance structure, cognition, clinical state, test-retest reliability, etc.), additional steps for increasing signal-to-noise ratio (SNR) (e.g., adopting inter-subject analysis, ISA), etc. The effects of preprocessing steps on functional connectivity estimation have been well-documented in previous studies (Satterthwaite et al., 2013; Vergara et al., 2017; Weissenbacher et al., 2009). Among the numerous preprocessing steps, head motion correction and global signal regression are two leading factors affecting functional connectivity estimation. The effects of data types on dFC estimation are tremendous and often reach divergent conclusions. For example, in simulated non-fMRI-BOLD data, Lindquist et al. (2014) demonstrated DCC’s superiority over SWFC in tracking the time-varying covariance structure. However, in real fMRI-BOLD data, DCC performed worse in tracking wakefulness states (Damaraju et al., 2020) and behavioral changes (Xie et al., 2019). The most likely reason is that in the study by Lindquist et al. (2014), the authors simulated non-fMRI-BOLD signals without considering the neurovascular effect, i.e., without convolving the hemodynamic response function. The low temporal resolution and hemodynamic response between neural activity and fMRI response determine the lower limits of state sojourn time and transition in fMRI data (Lindquist et al., 2009). To increase SNR, many strategies have been proposed, such as adopting a simple moving average on dFC results to increase the sensitivity (Shine et al., 2015; Xie et al., 2019), and adopting ISA in detecting stimulus-induced inter-regional correlation (Di and Biswal, 2020; Simony et al., 2016). The crux of ISA lies in the assumption that brain regions engaged by a task exhibit synchronous fluctuations across subjects, whereas regions unresponsive to stimuli do not. For task-fMRI data, adopting ISA for dFC estimation (e.g., ISSWFC, ISDCC) has been demonstrated as an effective way to isolate stimulus-induced inter-regional correlations and suppress effects from between-subject-unrelated spontaneous and non-neuronal signals.

It should be noted that the influences of the two factors are not isolated. SWFC performed well when window size and state changes were comparable (Xie et al., 2019). When the duration of state changes was randomly or smaller than the window size, SWFC often gave poor estimation (Shakil et al., 2016; Thompson et al., 2018).

The works mentioned above represent the current methodological understanding of the efficacy of dFC methods. However, several limitations of existent studies need to be addressed. First, the underlying covariance structure (or dFC patterns), sojourn time distributions, and precise state transition are unknown in real fMRI-BOLD datasets. Thus, selecting an appropriate benchmark or metric for efficacy assessment is challenging. Even using multitask scans as a framework, behaviorally relevant dFC patterns can be affected by overlapping representations between tasks (Gonzalez-Castillo et al., 2015). Second, although simulation data with predefined covariance structures are often used as benchmarks in efficacy comparisons (Lindquist et al., 2014; Shappell et al., 2019; Thompson et al., 2018), few simulation studies consider the neurovascular effect. i.e., simulated fMRI-BOLD datasets. Third, several widely used dFC methods, such as FLS (Liao et al., 2014), GLKF, HMM (Song et al., 2021b), and HSMM (Shappell et al., 2019), have never been systemically compared with other dFC methods.

This work aimed to systemically compare the efficacy of the seven dFC methods and their enhanced versions using simulated signals. We simulated two types of time series, one that did not consider the neurovascular effect (i.e., simulated non-fMRI-BOLD signals) and another that did (i.e., simulated fMRI-BOLD signals). For the first type, we simulated two time series with three kinds of predefined covariance structures. Each type of simulation was repeated 1000 times, maintaining the same covariance structures of interest while incorporating three levels of random Gaussian noise. For the second type, we simulated ten fMRI-BOLD time series with four connectivity patterns and two kinds of sojourn time distributions. Each kind of simulation was repeated 100 times, maintaining the same covariance structures of interest while incorporating simulated spontaneous signals (i.e., intrinsic neural signals or resting state signals) and three levels of random Gaussian noise. To suppress the effect of noise and spontaneous signals and increase the SNR of data of interest, we developed enhanced versions of the seven dFC methods by combining ISA.

## Method

### Simulated non-fMRI-BOLD datasets with predefined covariance structure (Simulated datasets 1, SD1)

The simulation method of SD1 is similar to Linquist et al. (2015), with only a few parameter differences. We simulated two time series with three predefined covariance structures. The sampling rate is 1 second. For the first simulation (SD1-1), the two univariate time series had zero mean and zero covariance throughout the time. In this setting, the estimated dFC should be close to zero (uncorrelated) across the entire range of the time course. For the second simulation (SD1-2), the two univariate time series had zero mean and a covariance distribution of Gaussian curves of three varying widths (20, 40, 60). The Gaussian covariance distribution represents a rapid change in the correlation coefficient. For the third simulation (SD1-3), the two univariate time series had zero mean and a covariance distribution of slowly varying periodic functions (sin(t/64), where t = 1,…600). The periodic covariance distribution represents a slowly varying correlation coefficient. Figure 1A depicts the covariance structures of the three kinds of simulation.

**Figure 1.**
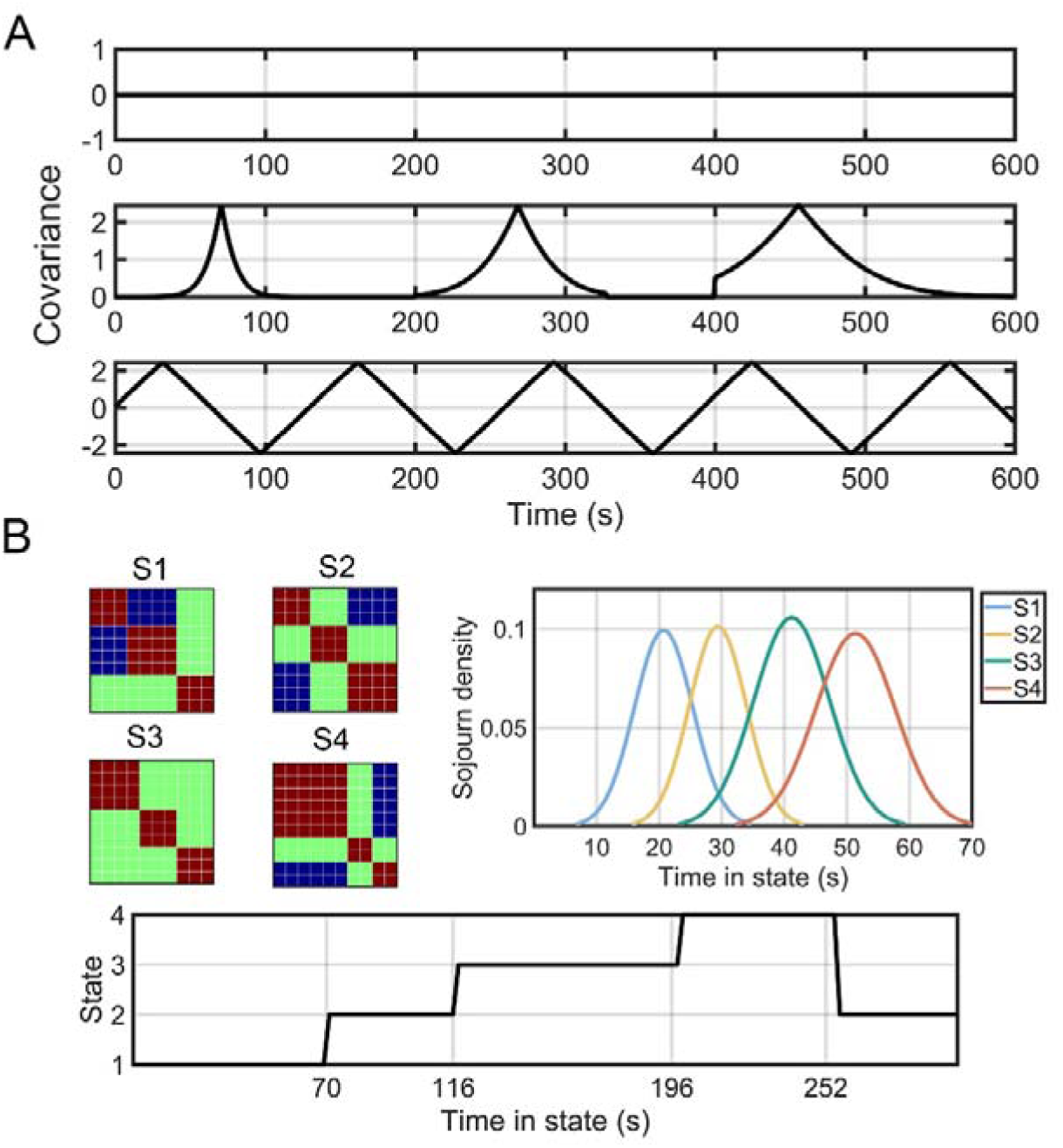
The two types of simulation methods. A: In the first type of simulation method, non-fMRI-BOLD signals were simulated, and the neurovascular effect was not considered. We simulated two time series with three predefined covariance structures: 1) Zero mean and zero covariance throughout the entire time course (SD1-1). 2) Zero mean with covariance distributed as Gaussian curves of varying widths (20, 40, 60) (SD1-2). 3) Zero mean with covariance distributed as slowly varying periodic functions (sin(t/64), where t = 1,…,600) (SD1-3). To evaluate noise sensitivity, Gaussian noise was added to the simulated time series with standard deviations of 0.1 (SD1-1-1, SD1-2-1, SD1-3-1), 0.3 (SD1-1-2, SD1-2-2, SD1-3-2), and 0.6 (SD1-1-3, SD1-2-3, SD1-3-3). Each noise condition was simulated 1000 times. All time series had a length of 600. B: In the second type of simulation method, we incorporated the neurovascular effect by convolving the time series with a hemodynamic response function to simulate fMRI-BOLD signals. Ten fMRI-BOLD time series with four predefined states, modular structures, and state sojourn distributions were simulated. Two state sojourn distributions were generated: 1) A state-transition vector of 1–2–3–4–2, with dwell times of 70, 46, 80, 56, and 48 seconds (SD2-1, totaling 296 seconds). 2) Poisson-distributed state sojourns with λ = 20, 30, 40, and 50 for states 1, 2, 3, and 4, respectively (SD2-2). To make the signals more similar to real fMRI-BOLD signals, we added between-data-unrelated spontaneous signals with a probability of 0.1 and Gaussian noise with standard deviations of 0.1 (SD2-1-1 and SD2-2-1), 0.3 (SD2-1-2 and SD2-2-2), and 0.6 (SD2-1-3 and SD2-2-3). Each spontaneous and noise condition was simulated 100 times and added to the data of interest.

To evaluate the sensitivity to noises, Gaussian noise was added to the above simulated time series with a standard deviation of 0.1 (SD1-11, SD1-21, SD1-31), 0.3 (SD1-12, SD1-22, SD1-32) and 0.6 (SD1-13, SD1-23, SD1-33), respectively. Each set of noise data was simulated 1000 times. The length of the time series was 600.

### Simulated fMRI-BOLD time series with predefined connectivity states and state sojourn distribution (SD2)

SD2 simulated two event-related fMRI-BOLD datasets. The first simulation (SD2-1) was identical to the one used by Allen et al. (2014). The fMRI-BOLD signals were generated with four discrete states and ten regions of interest. The four neural connectivity patterns (states) were with distinct modular structures (Fig.1B). We simulated 100 datasets with 148-time points and a repetition time (TR) of 2 seconds. The state-transition vector was 1–2–3–4–2, and the dwell times were: 70, 46, 80, 56, and 48 seconds. The event-related time course is created by convolving the neural events with the canonical hemodynamic response function in SPM12 (https://www.fil.ion.ucl.ac.uk/spm/software/spm12/). To evaluate the efficacy of dFC methods unveiling the task-related effects, we simulated between-data-unrelated intrinsic neural signals (i.e., spontaneous signals). For each TR, the intrinsic neural signals were randomly generated from a standard uniform distribution with a certain probability (0.1) and then added to the event-related time course. To make the signals more similar to the real BOLD signals, we added Gaussian noises with a standard deviation of 0.1 (SD2-1-1), 0.3 (SD2-1-2), and 0.6 (SD2-1-3), respectively. In real task fMRI data, the intrinsic neural signals were dominant, and the percent of signal change induced by task stimulation was ∼1%–5% (Fox and Raichle, 2007). Thus, in this study, the amplitude of intrinsic neural signals was three times of the event-related data.

In the second simulation (SD2-2), the same neural connectivity patterns were simulated except for the state sojourn distribution. The state sojourn distributions were Poisson distributed with λ = 20, 30, 40, and 50 for states 1, 2, 3, and 4, respectively (Fig.1B) (Shappell et al., 2019). For each TR, the intrinsic neural signals were randomly generated with a certain probability (0.1), multiplied by 3, and then added to the event-related time course. Gaussian noises with a standard deviation of 0.1 (SD2-2-1), 0.3 (SD2-2-2), and 0.6 (SD2-2-3) were then added to the simulated data, respectively.

Examples of signals in SD2-1-2 and SD2-2-2 were presented in Supplementary Figure 36.

### The seven dFC methods

Table 1 summarizes the seven dFC methods, with the mathematical functions detailed in the supplementary materials. Prior research has documented the effects of free parameters on each dFC method. Therefore, in this study, we used the optimal parameters recommended by these previous studies and focused on comparing the efficacy of different dFC methods.

**Table 1.**
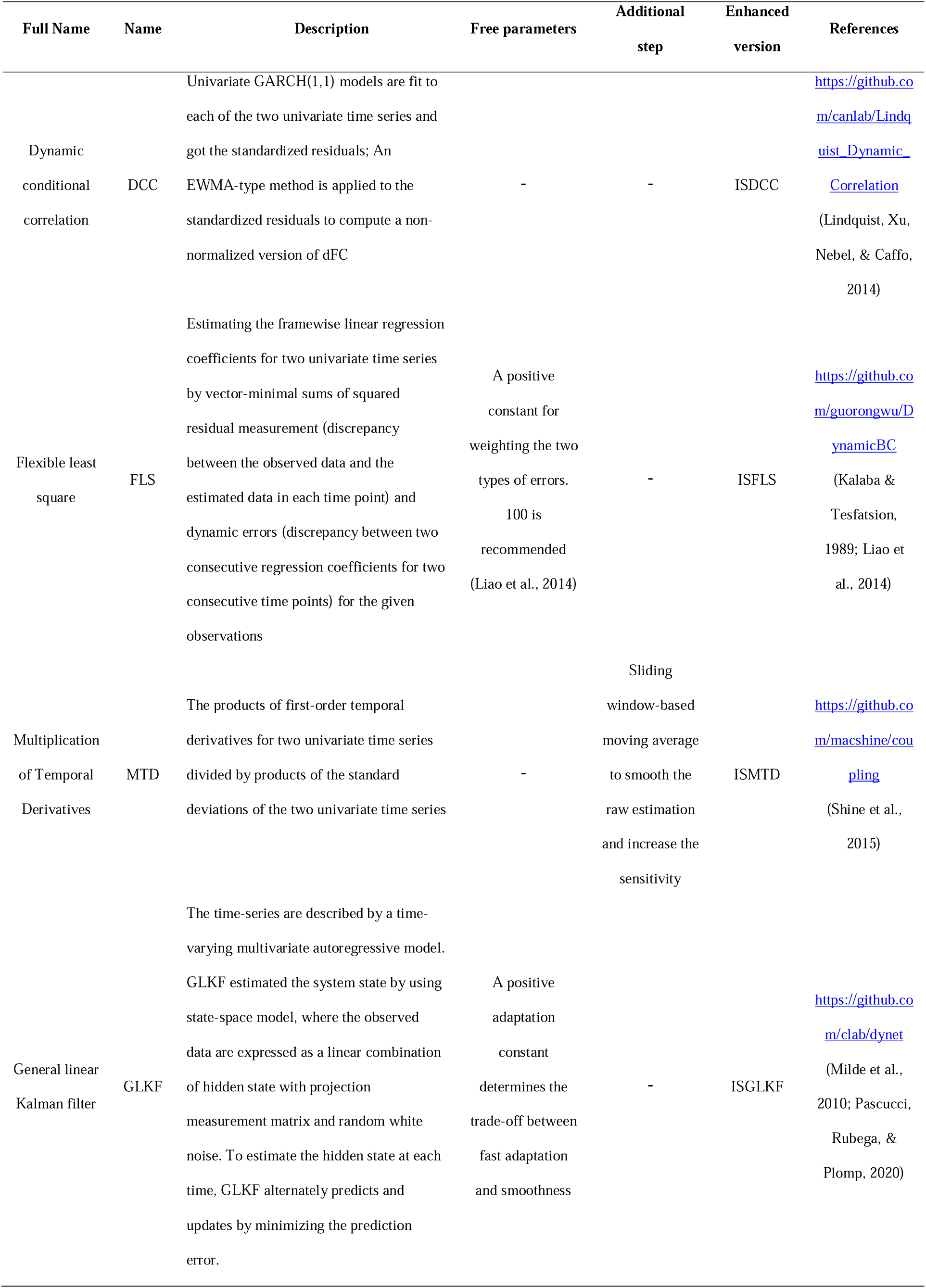
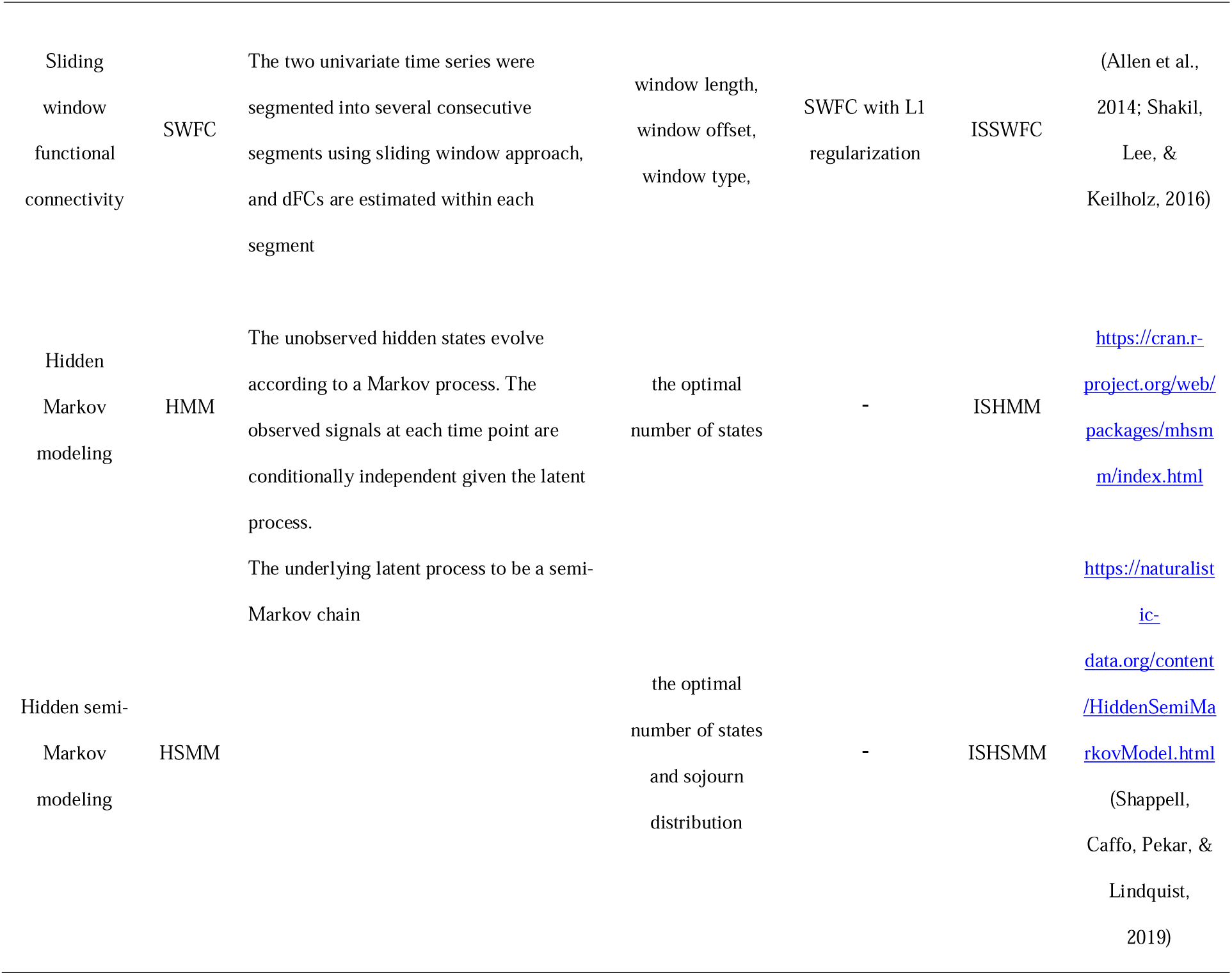
A summary of the seven dFC methods and their enhanced versions.

### Flexible Least Squares (FLS)

FLS treated dFC estimation as a dynamic regression coefficients estimation in a time-series linear regression model (Kalaba and Tesfatsion, 1989). It assumed that the regression coefficients evolved across time slowly and the model errors were related to two types of errors: 1) residual measurement error given by the discrepancy between the observed dependent variables and the estimated linear regression model at each time; 2) the residual dynamic error given by the discrepancy between two consecutive regression coefficients. The FLS solution is defined to yield both the lowest residual measurement error and dynamic error for the given time series.

### Dynamic conditional correlation (DCC)

DCC was first proposed by Engle (2002) and widely used in the finance literature. Lindquist et al. (2014) introduced DCC to characterize dFC in resting state fMRI. The DCC algorithm adopts generalized autoregressive conditional heteroscedastic (GARCH) to express the conditional variance of a single time series at time *t* as a linear combination of the conditional variance plus the squared values of the previous time point. To compute the time-varying conditional correlation between two univariate time series, an exponentially weighted moving average (EWMA) window is applied to the standardized residuals obtained by the GARCH model. In contrast to sliding-window methods, EWMA applies declining weights to the past observations in the time series, and the most recent time points are placed in the heaviest weights. DCC has shown its accuracy and reliability than SWFC (Choe et al., 2017) in simulation data. However, Damaraju et al. (2020) showed that SWFC outperformed DCC in classifying sleep states, which may be due to the higher sensitivity of DCC to noise than smoother sliding window estimates.

### General linear Kalman filter (GLKF)

GLKF is a technique for estimating system state and covariance for multi-trial multivariate time series (Milde et al., 2010). In GLKF, the current observed data are expressed as a linear combination of the hidden state with the projection measurement matrix. The filter alternates two main steps to estimate the hidden state: prediction and update. In the prediction step, the filter recursively estimates the hidden state at each time. In the update step, the filter compares the predicted state and covariance with the actual measurement.

### Multiplication of temporal derivatives (MTD)

MTD consists of two steps. For two time series *x*_*i*_(*t*) and *y*_*i*_(*t*), their temporal derivatives (*td*_*x*_ and *td*_*y*_), i.e., the temporal difference between time *t* and *t*-1, are firstly calculated. The two temporal derivatives are then multiplied at each time point and divided by the product of the standard deviation for *td*_*x*_ and *td*_*y*_. Positive MTD value at time *t* indicate that *x*_*i*_(*t*) and *y*_*i*_(*t*) cofluctuate in the same direction, whereas negative value indicates *x*_*i*_(*t*) and *y*_*i*_(*t*) fluctuate in the opposite direction. To reduce the susceptibility to very high-amplitude fluctuations or outliers, e.g., high volatility due to head motion, MTD values were smoothed by applying a sliding-window average. Like the SWFC, long window lengths are insensitive to rapid network reorganization, whereas short window lengths make MTD more susceptible to outliers (Shine et al., 2015; Esfahlani et al., 2019; Xie et al., 2018). This study chose a window length of 5 time points, an optimal value with high sensitivity and specificity (Shine et al., 2015).

### Sliding-window functional connectivity with L1-regularization (SWFC)

The SWFC was equal to the Pearson correlation coefficient of two windowed time series. This study used a Gaussian tapered window by convolving a rectangle with a Gaussian kernel (σ=3). The window length was 22 time points. To increase the estimation accuracy, L1 regularization was adopted. The free parameter λ, which enables a trade-off between the sparsity of the inverse covariance matrix and goodness of fit, was estimated within each data point.

### Hidden Markov models (HMM) and Hidden semi-Markov models (HSMM)

In HMM, it is assumed that the observed data are generated by a finite set of hidden states, and the Markov chain governs the relationship between states. The current state was conditionally predicted by the probability of the previous state and transition probability matrix. HMM and HSMM differ in whether the underlying latent process is a Markov or semi-Markov chain (Shappell et al., 2019; Yu, 2010). A Markov chain assumes that the conditional independence between two states is ensured every time. The sojourn time (i.e. number of consecutive time points in a state) distribution for a given state was implicitly assumed as geometrically distributed. In contrast, a semi-Markov chain assumed that conditional independence was only ensured when state transition occurred. The sojourn time in a state was variable and could be explicitly defined. Shappell et al. (2019) demonstrated that HSMM is more flexible than HMM in estimating state sojourn distribution in simulation and real fMRI data.

The parameters of the two models were estimated using the mhsmm package for R (O’Connell and Højsgaard, 2011), based on applying the Expectation-Maximization (EM) and forward-backward algorithms (Guédon, 2003; Shappell et al., 2019). The EM algorithm estimated all model parameters by alternatively performing an expectation (E) step and a maximization (M) step. In the E-step, the probability of a state at each time, the state-transition probability at each time, and the state sojourn time (only for HSMM) were estimated using the forward-backward algorithm. In the M-step, the expected log-likelihood was maximized. These E-and M-steps were repeated until convergence. For HSMM, the state duration density was also estimated.

### Enhanced versions by adopting ISA

We enhanced the abovementioned dFC methods by adopting ISA and only focused on the interest component. For a simulated data *i* without the neurovascular effects, denoted by *i* = 1, 2, …, 1000, the simulated signals, *x*_*i*_(*t*), have two elements: a shared component (*s*_*i*_(*t*)) and noises signals (*n*_*i*_(*t*)). While for the simulated BOLD signals *x*_*i*_(*t*), have three elements: a shared component (*s*_*i*_(*t*)), data-specific spontaneous fluctuations (*i*_*i*_(*t*)), and noise signals (*n*_*i*_(*t*)). By adopting ISA, we can firstly estimate the *s*_*i*_(*t*) by simply averaging the *x*_*i*_(*t*) across a subset of subjects and cancels out subject-specific components (i.e., *i*_*i*_(*t*) and *n*_*i*_(*t*)) through linear regression or directly calculated dFC between *x*_*i*_(*t*) and the estimated *s*_*i*_(*t*). To estimate shared components, we adopted the leave-one-out method. For n times simulations, the time series of the n-1 simulations were averaged. For the first five dFC methods, the data of the ith simulation and the averaged data were merged as one data matrix. For the new data matrix, dFCs were calculated, and the upper right corner of the correlation matrix is extracted as the inter-subject dFC matrix at this frame or window. For HMM and HSMM, we performed linear regression analysis with the averaged *s*_*i*_(*t*) to regress out spontaneous fluctuations and noise signals. The seven new dFC methods were renamed as ISSWFC (Di and Biswal., 2020), ISMTD, ISDCC (Chen et al., 2024), ISGLKF, ISFLS, ISHMM, ISHSMM. The enhanced versions were expected to increase their sensitivity to signals of interest and suppress the data-specific effects of spontaneous signals and random Gaussian noise

### K-means clustering

For simulated fMRI-BOLD data, to identify the temporally reoccurring states, the k-means clustering algorithm was adopted to decompose the functional connectivity matrices into several non-overlapping clusters (Yuan et al., 2023a; Yuan et al., 2023b). The matrices were assigned into different clusters (i.e., states) according to the similarity in their spatial patterns. The sum of the distances between the individual matrix and the centroid of the cluster they belonged to is at the minimum for each cluster. The centroid was regarded as the representative pattern for the given cluster. The optimal number of clusters k was estimated based on the elbow criterion, i.e., the ratio between the within-cluster distance to between-cluster distances. L1 distance function (‘Manhattan distance’) was implemented to assess the point-to-centroid distance. The matrices of each frame were finally assigned to one of these clusters.

### Maximum likelihood estimation for the HMM and HSMM

Estimating hidden Markov or semi-Markov chains was based on applying the Expectation-Maximization (EM) and forward-backward algorithms (Guédon, 2003; Shappell et al., 2019). The EM algorithm estimated all model parameters by alternatively performing an expectation (E) step and a maximization (M) step. In the E-step, the probability of a state at each time, the state-transition probability at each time, and the state sojourn time (only for HSMM) are estimated using the forward-backward algorithm. In the M-step, the expected log-likelihood is maximized. These E-and M-steps are repeated until convergence. For HSMM, the state duration density is also estimated.

### Index of dFC efficacy

SD1 simulation aimed to evaluate the ability of dFC methods to track the time-varying covariance structure. Thus, the efficacy for SD1 data was defined as the mean squared error (MSE) between the true correlation vector and the estimated correlation vector. Note that for metrices that are not framewise, the length of the estimated correlation vector is not 600. For SWFC and ISSWFC, the length is 578. For MTD and ISMTD, the length is 599.

In SD2, both spatial and temporal properties were predefined. The spatial efficacy was defined as the spatial Pearson correlation between the real state and the estimated state. We adopted two metrics to assess the temporal efficacy. Firstly, we calculated the Pearson correlation between the predefined state transition vector and the estimated state transition vector. The second method is to calculate the Fréchet distance between the true and estimated sojourn density curves. For the three datasets in SD2-2, the sojourn density curve of the estimated state transition vector was estimated by first calculating the sojourn times for each state and then fitting them against the Poisson distribution. The Fréchet distance is a measure of similarity between two curves or paths (Eiter and Mannila, 1994).

## Results

### Overall performance in SD1

In the first type of simulation, the covariance between the two-time series is time-varying. HMM and HSMM classically focus on the discretization of state sequences; therefore, we only present the results of the first five methods and their enhanced versions. From the MSE results of the nine datasets (three covariance structures x three levels of SNR), we can derive three basic conclusions: first, the enhanced dFC methods generally outperformed their original versions. Second, the efficacy depends on the covariance structure between the two time series. Overall. DCC and FLS based methods performed better than the other three methods. Third, as the standard deviation of random noise increases, the accuracy of the estimated correlation coefficient decreases rapidly, and the MSE increases sharply.

### Estimated time-varying structure of covariance for SD1

Figures 2 and 3 show the results on simulated datasets SD1-1-2. The ISDCC performs best, followed by the ISFLS and then ISGLKF. ISSWFC and ISMTD perform poorly. The original versions of all methods are slightly less efficient than the enhanced versions. Similar results were obtained on simulated datasets SD1-1-1 (Supplementary Figures 1 and 2) and SD1-1-3 (Supplementary Figures 3 and 4).

**Figure 2.**
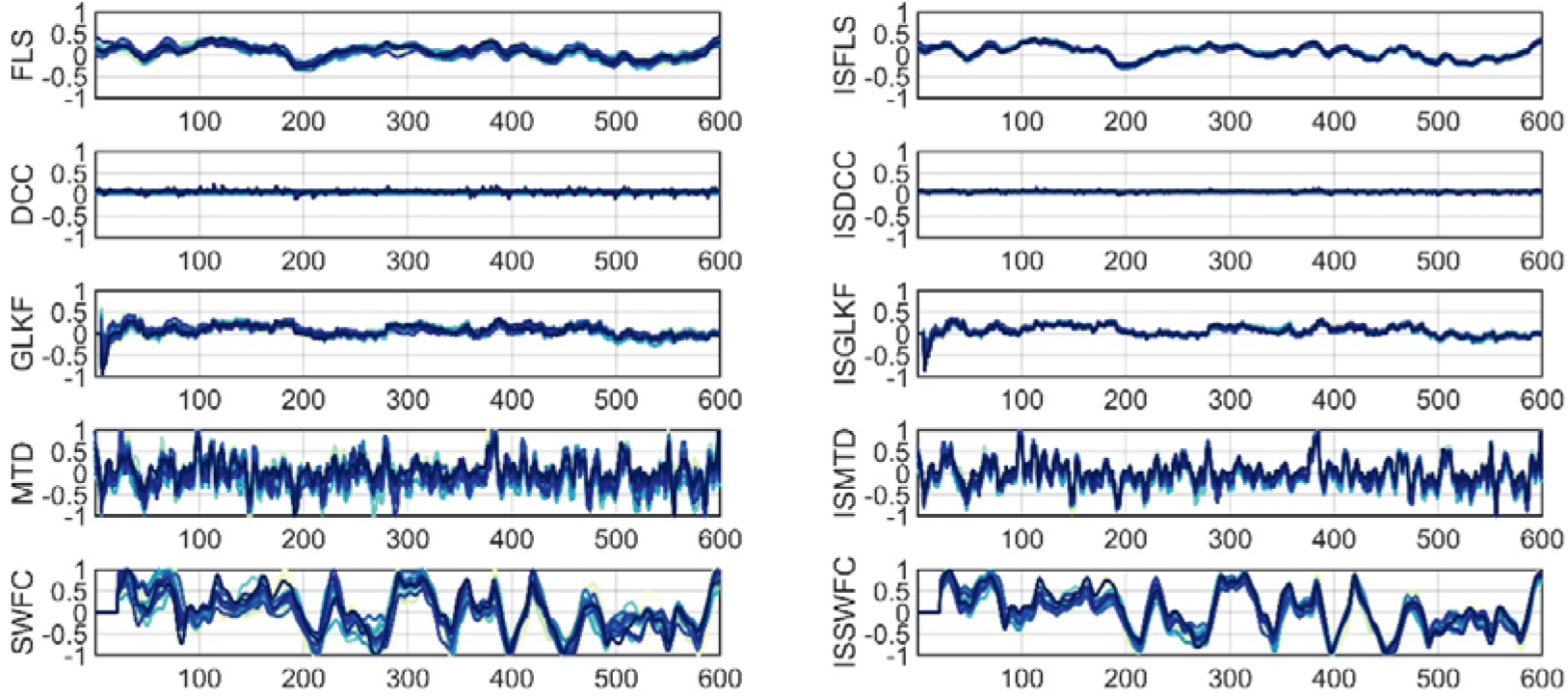
Example results of SD1-1-2. The estimated correlation coefficients of two time series with zero covariance from 20 simulations are presented.

**Figure 3.**
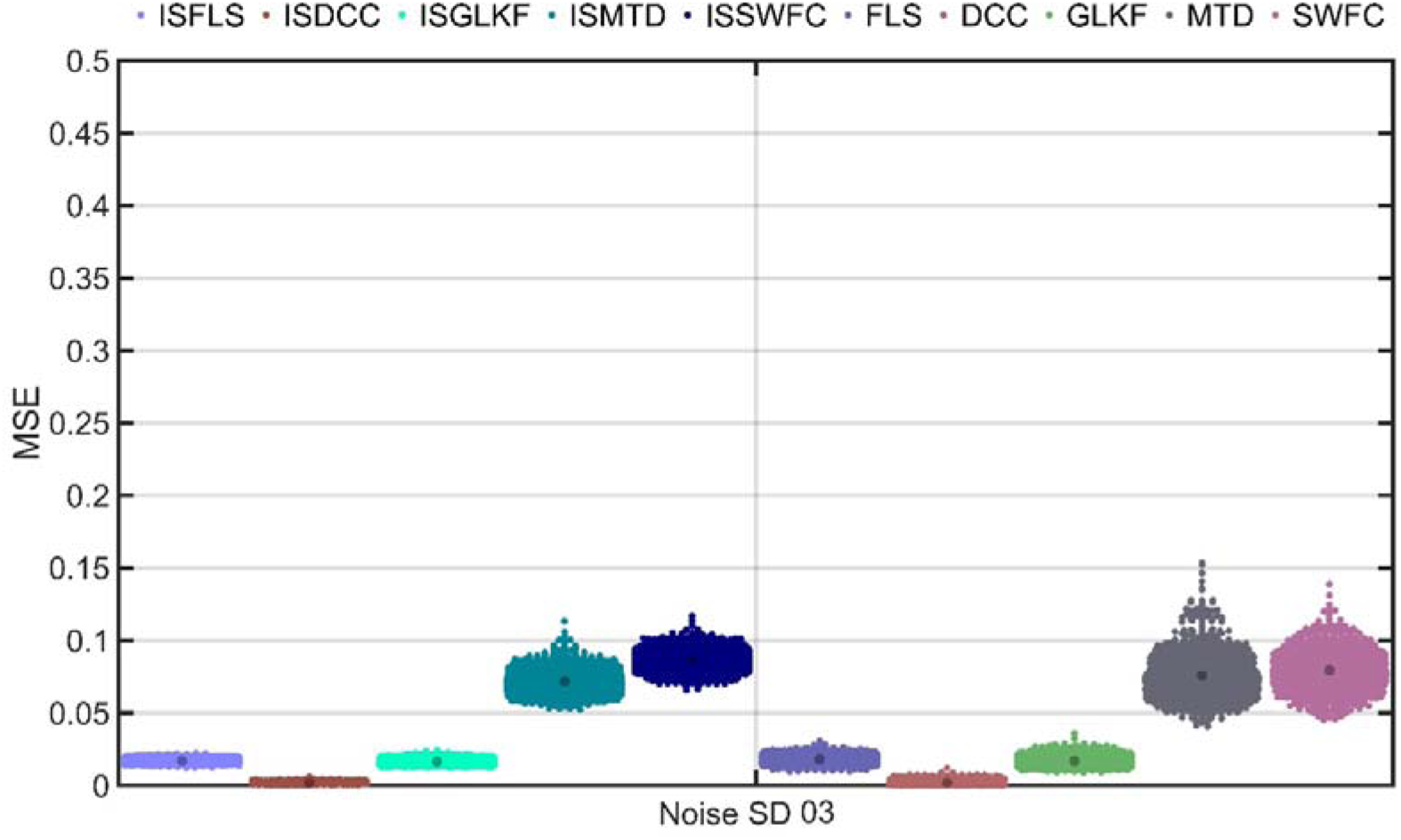
MSE results of SD1-1-2. The MSE between the estimated and real correlation coefficients of two time series with zero covariance from 1000 simulations are presented.

For the three simulated datasets of SD1-2 (Gaussion), ISFLS can accurately estimate the covariance structure with a Gaussian distribution, resulting in the smallest MSE.

ISDCC and ISGLKF were slightly worse. ISSWFC and ISMTD performed the worst (Figures 4 and 5, Supplementary Figures 5–8). For the three simulated datasets of SD1-3 (periodic), the best performers are still ISFLS and ISDCC, followed by ISSWFC and ISMTD, with ISGLKF performing the worst (Figures 6 and 7, Supplementary Figures 9–12).

**Figure 4.**
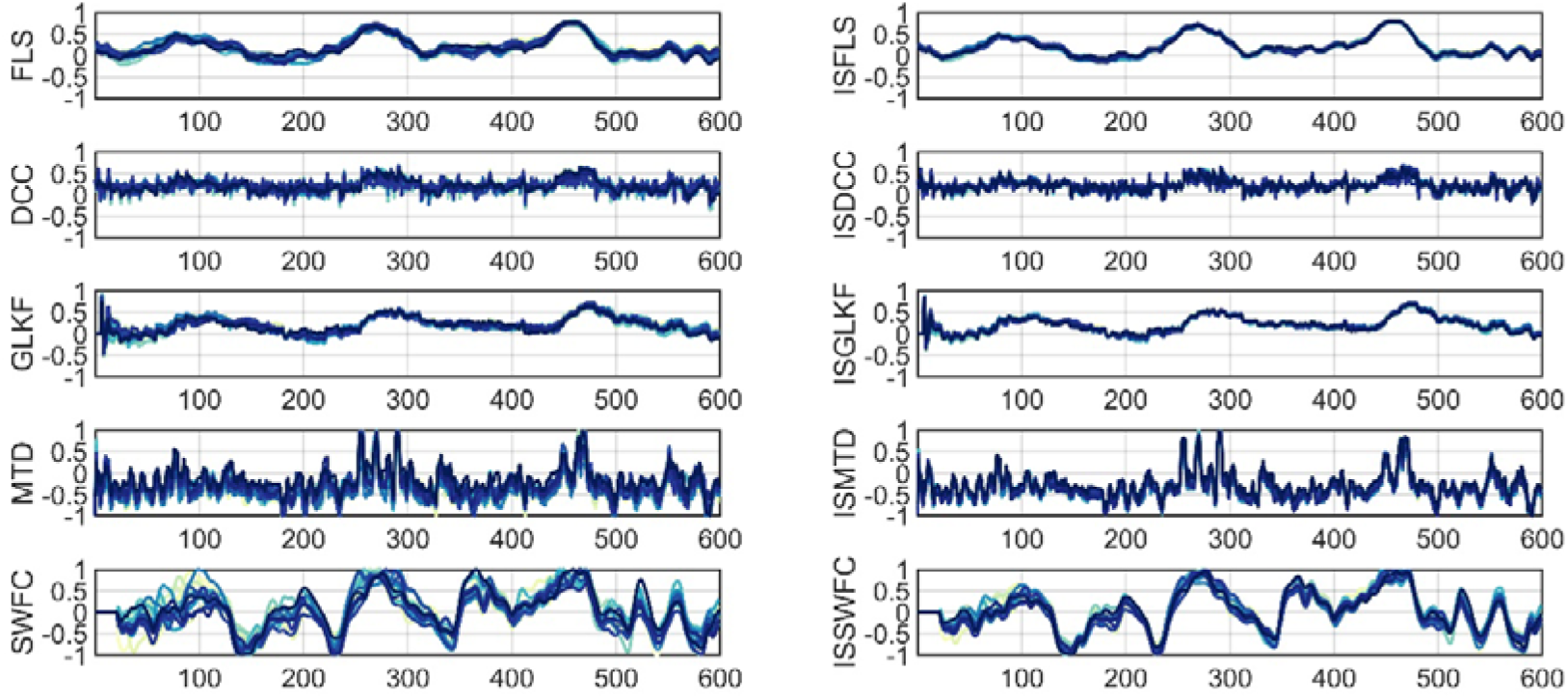
Example results of SD1-2-2. The estimated correlation coefficients of two time series with covariance structure of Gaussian distribution from 20 simulations are presented.

**Figure 5.**
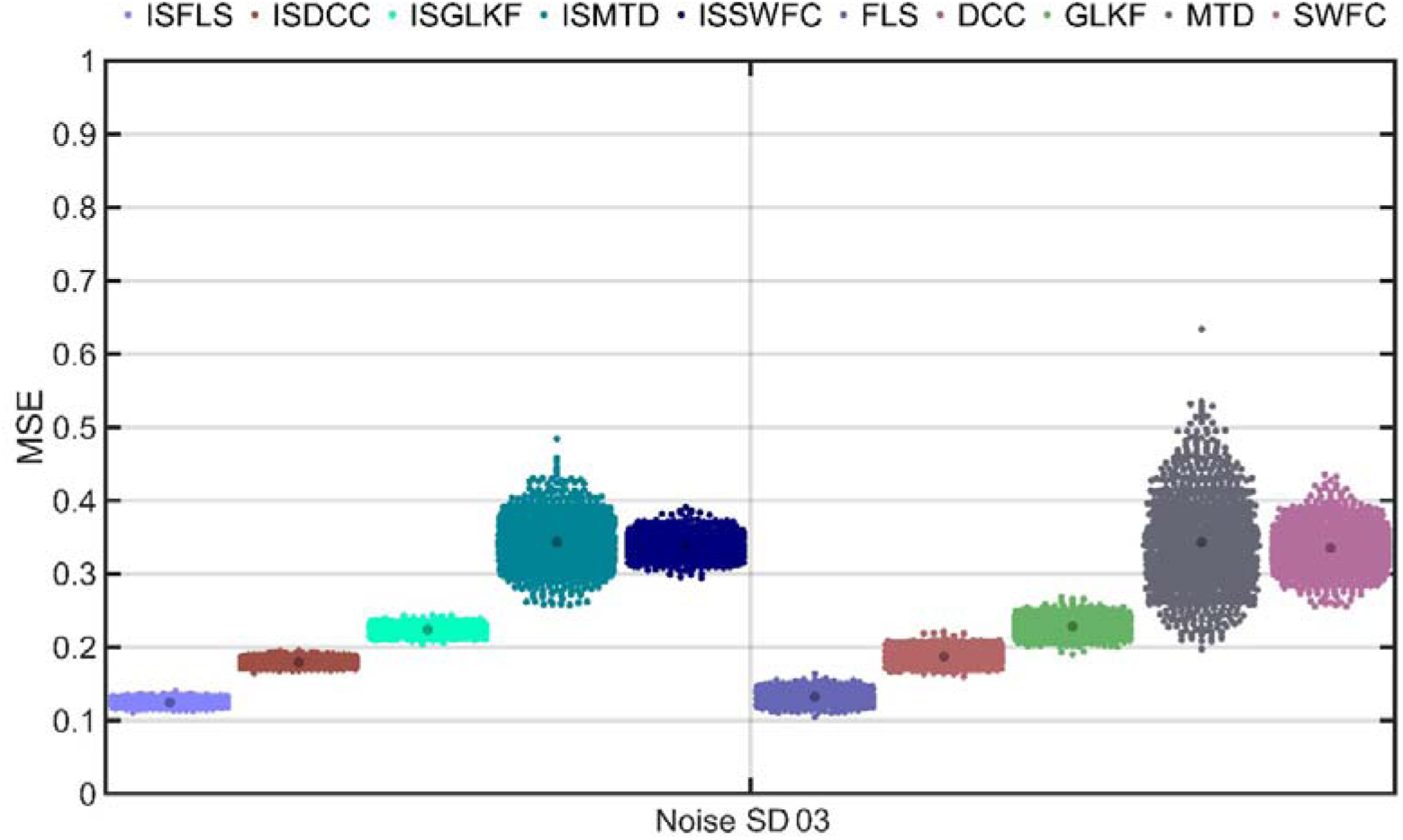
MSE results of SD1-2-2. The MSE between the estimated and the real correlation coefficients of two time series with covariance structure of Gaussian distribution from 1000 simulations are presented.

**Figure 6.**
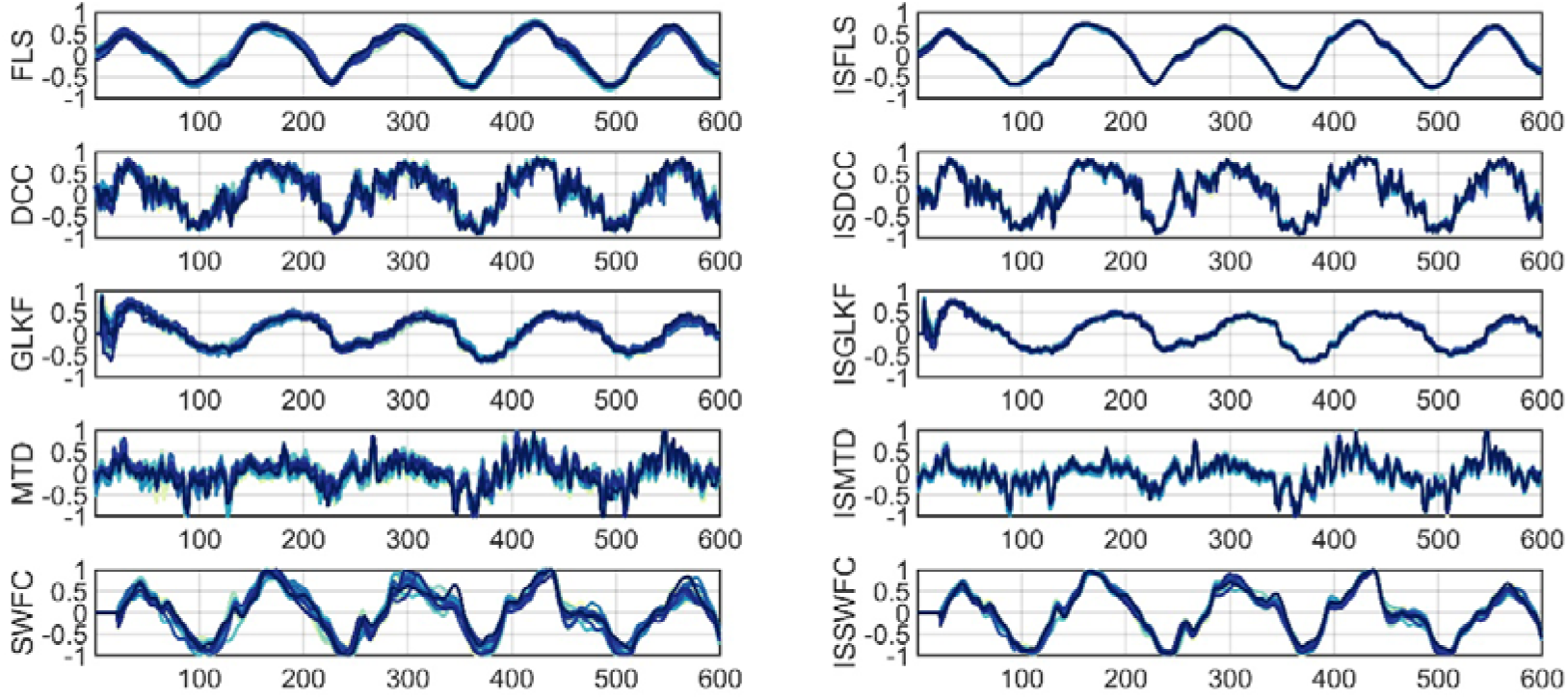
Example results of SD1-3-2. The estimated correlation coefficients of two time series with covariance structure of periodic distribution from 20 simulations are presented.

**Figure 7.**
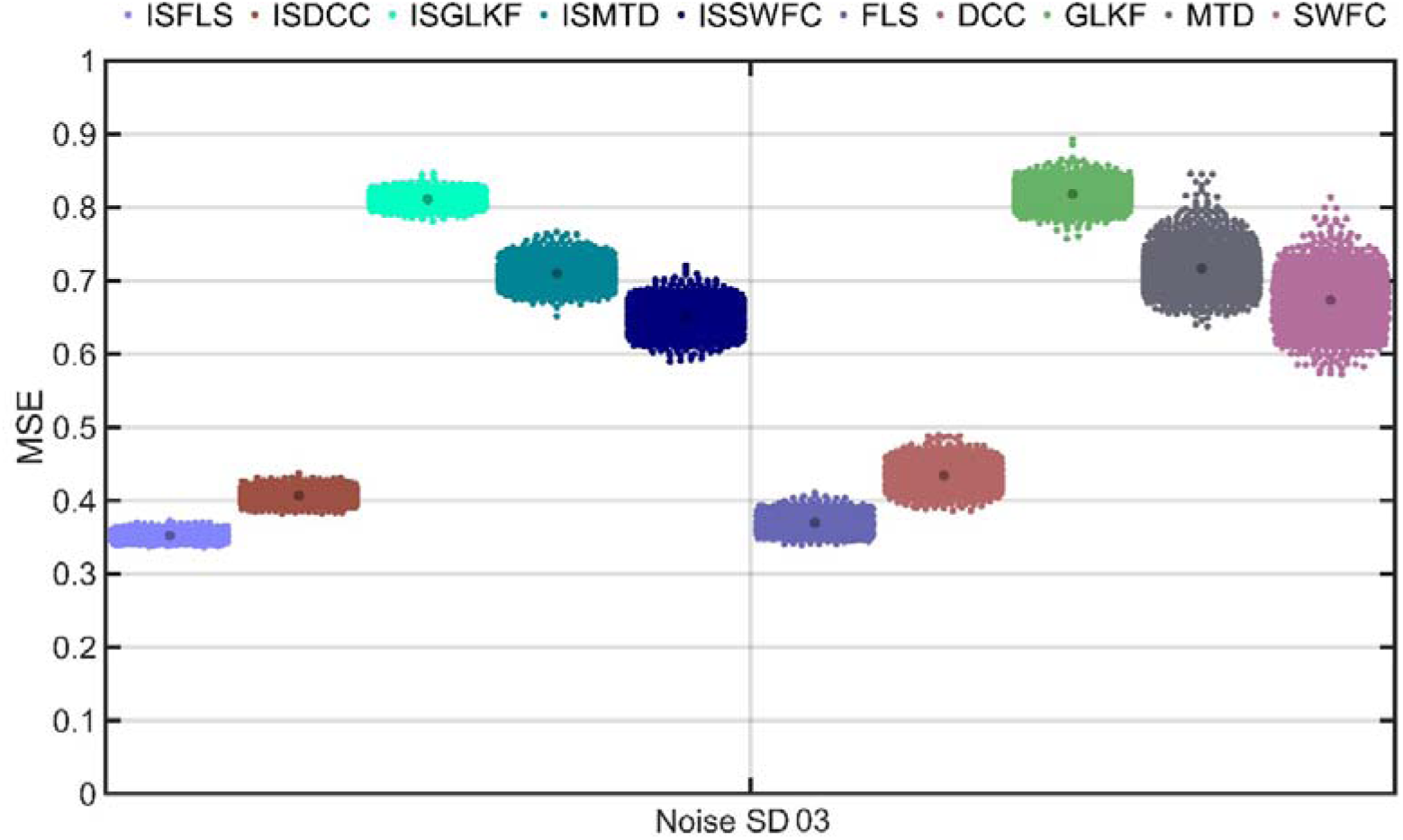
MSE results of SD1-3-2. The MSE between the estimated and the real correlation coefficients of two time series with covariance structure of periodic distribution from 1000 simulations are presented.

### Overall performance of SD2

For all datasets of the second type of simulation, we set the state number as 4. By comparing all results’ spatial and temporal correlations, we can draw the following conclusions: First, similar to SD1, the enhanced methods perform better than the original methods. All seven methods’ enhanced versions show higher inter-data consistency of state transition vectors than their original versions. Second, in all simulated datasets of SD2, ISHSMM, ISFLS, and ISSWFC performed well in estimating the four states and their sojourn times. However, among the remaining methods, either they could not accurately assess the connectivity patterns of the four states (e.g., ISGLKF), or they could not accurately estimate the state sojourn distribution (e.g., ISDCC and ISHMM). Third, as the standard deviation of random noise increases, the accuracy of spatial and temporal properties of all methods decreases. Fourth, the accuracy of different methods of spatial property estimation also seems state-dependent. Some methods can accurately estimate the connection patterns between nodes in certain states but fail to do so in other states.

### Estimated spatial maps and temporal transitions of SD2

Figures 8 and 9 present the estimated states and state transition vectors of SD2-1-2, and Figures 10 and 11 present the spatial and temporal correlation results of SD2-1-2. ISHMM and ISHSMM perform the best in estimating both spatial and temporal properties, as they had the highest spatial correlations of state maps (Figure 10) as well as the highest temporal correlations for the state transitions (Figure 11). They are followed by ISFLS and ISGLKF. Due to the inherent limitation of the sliding window approach, ISSWFC consistently exhibits latency in estimating the state transition. While it demonstrates robust performance in determining the spatial characteristics of states 3 and 4, its efficacy in capturing the spatial attributes of states 1 and 2 is notably less precise. ISDCC accurately estimates the spatial properties of states but has poor accuracy in estimating temporal properties. ISMTD performs the worst in estimating the spatiotemporal properties of states. Similar results were observed in SD2-11 (Supplementary Figures 13–16) and SD2-13 (Supplementary Figures 17–20).

**Figure 8.**
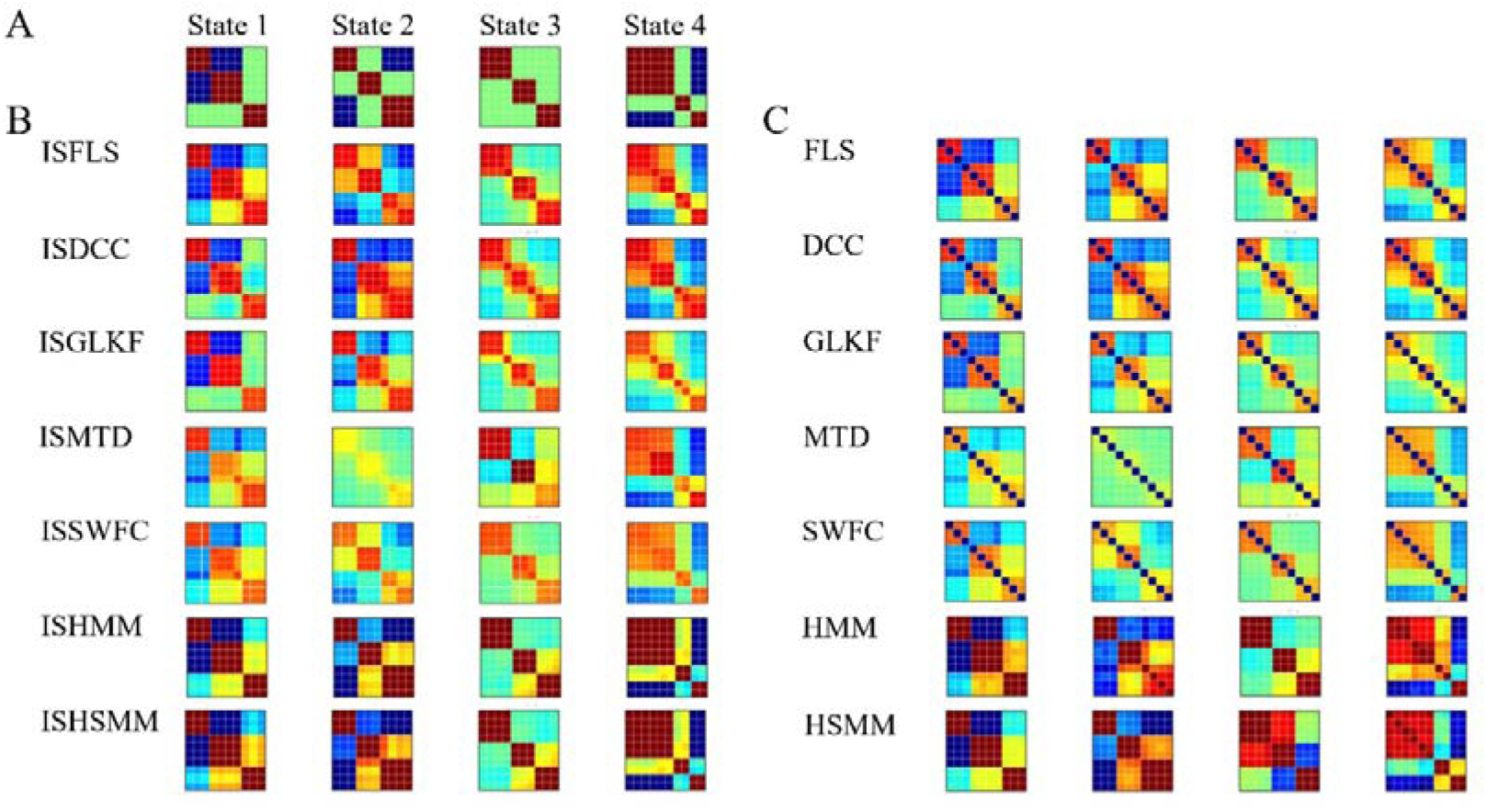
The predefined and estimated states of SD2-1-2 were determined using the 14 dFC methods.

**Figure 9.**
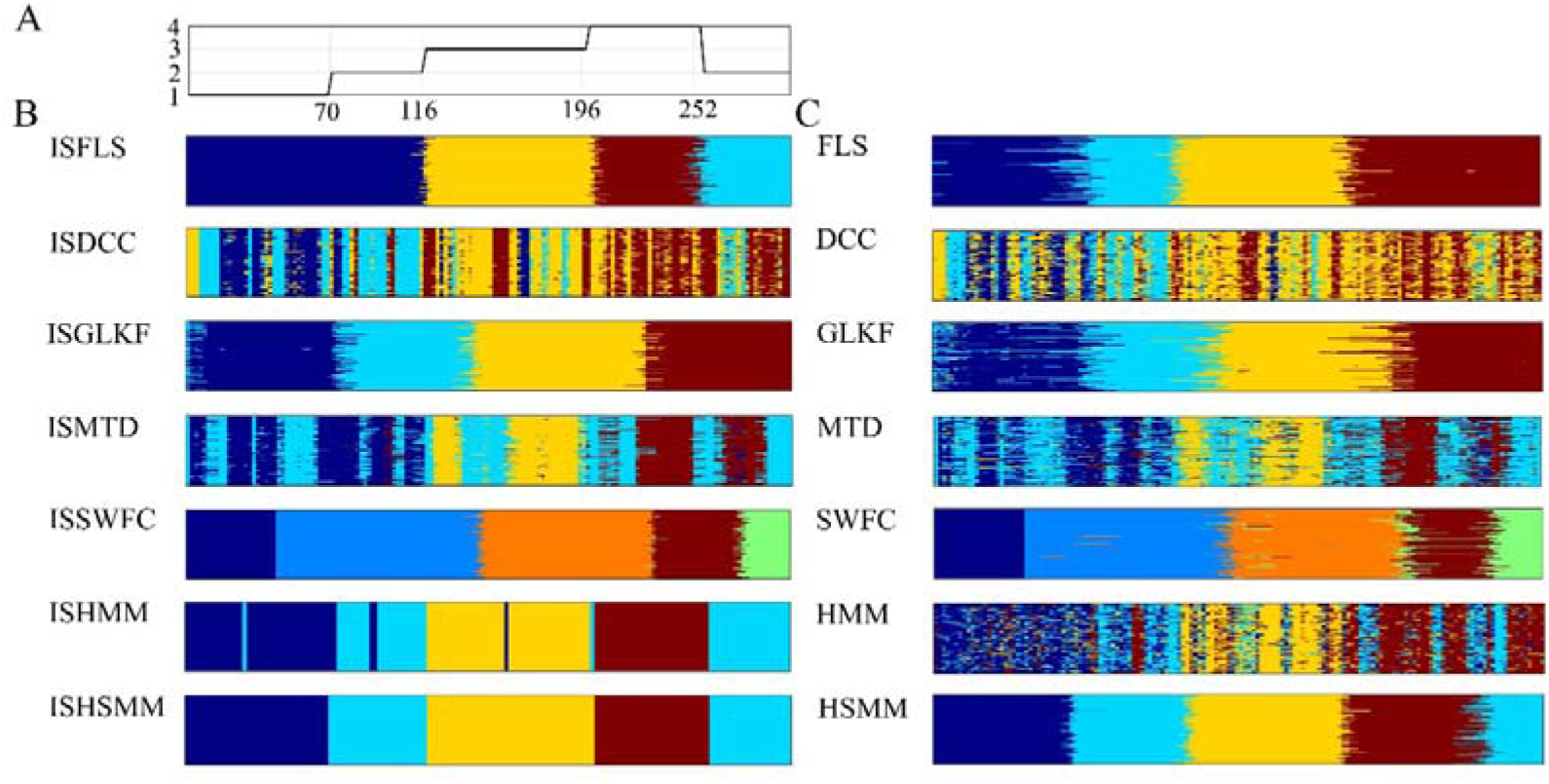
The estimated state transition vectors of SD2-1-2 by the 14 dFC methods. A: The predefined state transition vector of the four states. B & C: the color bar represents the state transition vectors for 100 simulated datasets, where rows represent the state transition vectors for each simulation and columns represent the number of simulations. For all methods except ISSWFC and SWFC, dark blue, light blue, yellow, and red correspond to states 1, 2, 3, and 4, respectively. For ISSWFC and SWFC, due to the sliding window, dark blue represents the null state, while light blue, orange, red, and light green corresponds to states 1, 2, 3, and 4, respectively.

**Figure 10.**
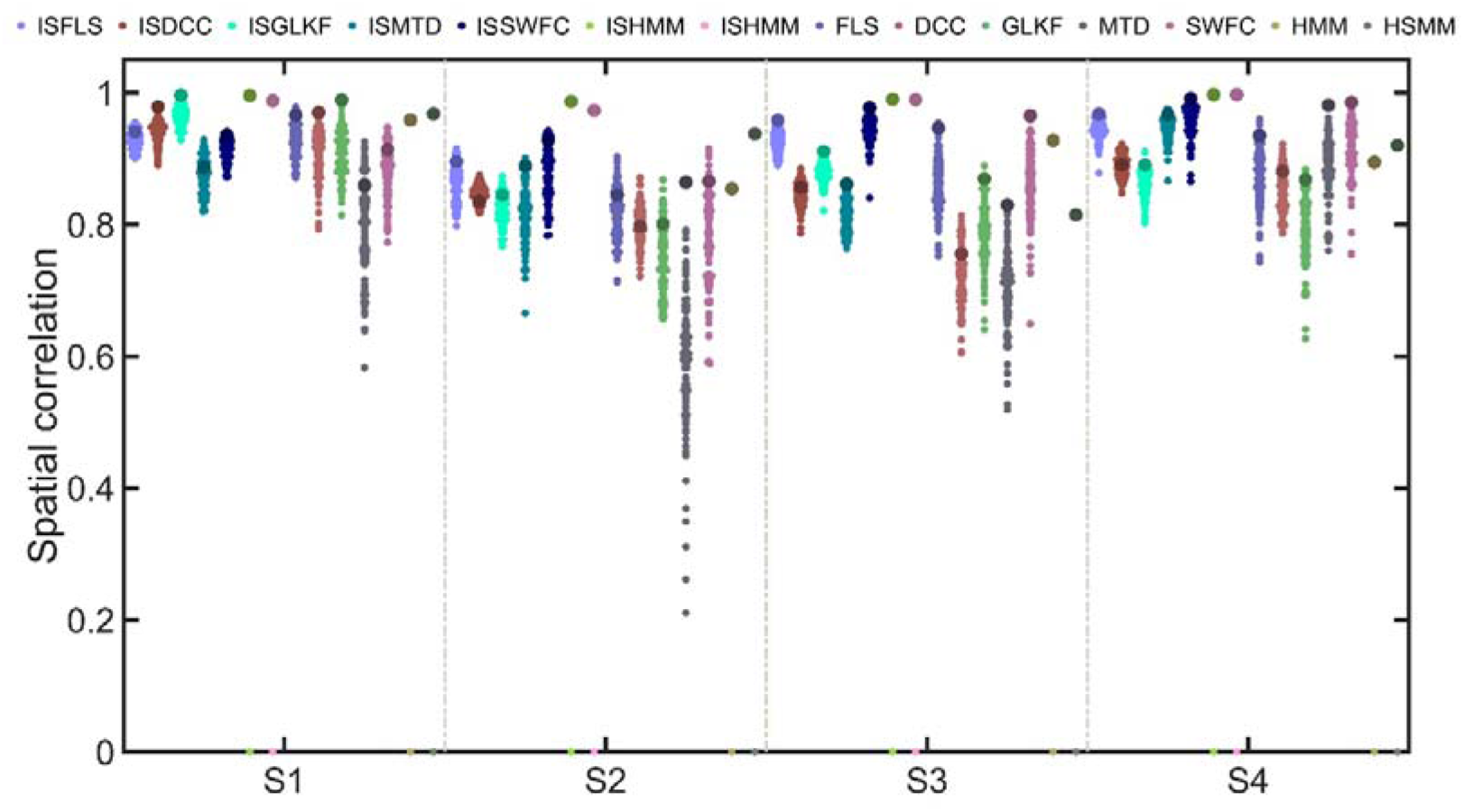
The spatial correlations of SD2-1-2 between the predefined and estimated states by the 14 dFC methods. There were no data-specific results for HMM and HSMM and their enhanced versions, ISHMM and ISHSMM.

**Figure 11.**
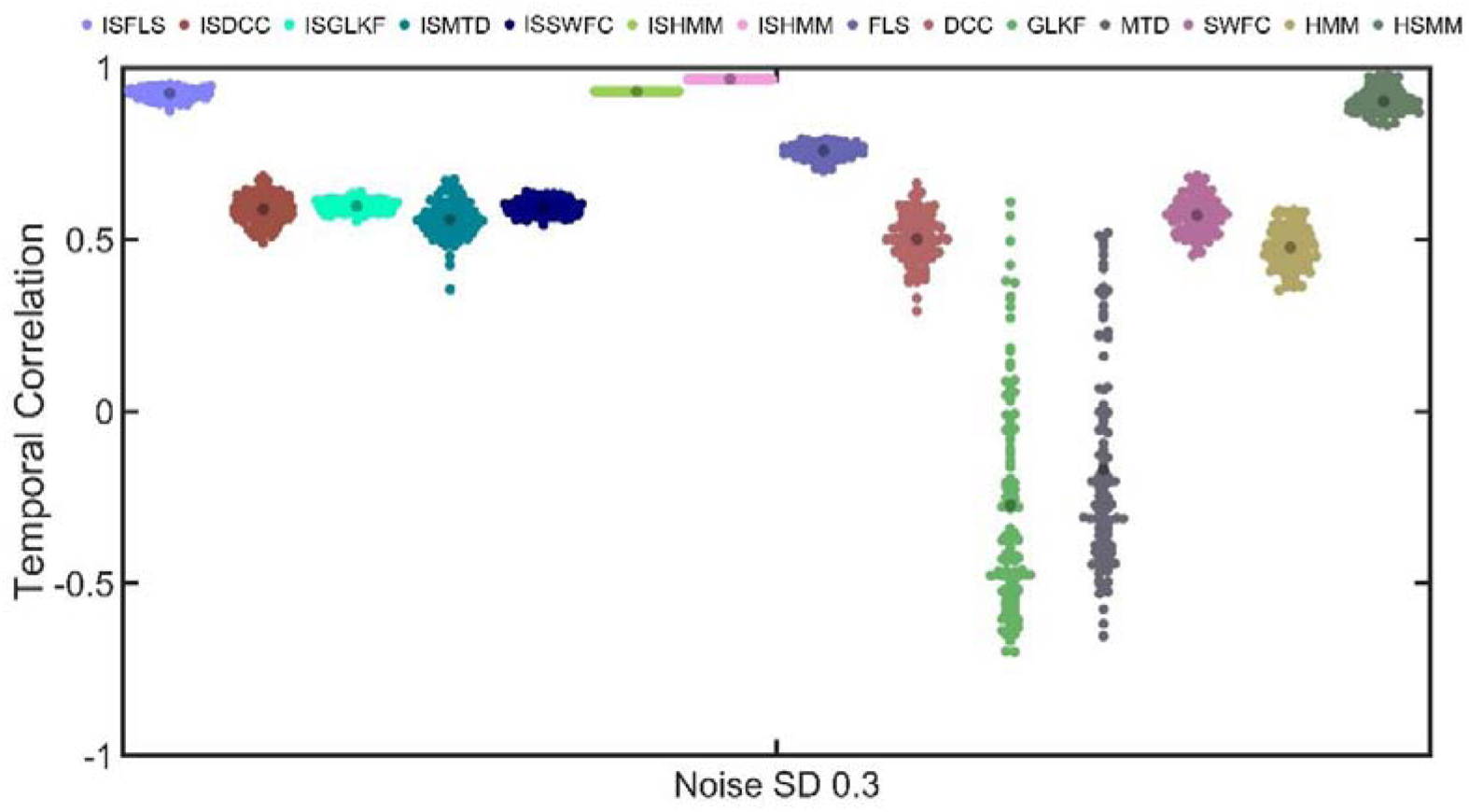
The temporal correlations of SD2-1-2 between the predefined state transition vectors and the estimated state transition vectors by the 14 dFC methods.

### Estimated distribution of sojourn time in SD2

Compared to SD2-1, the primary difference in dFC efficacy across the three simulated datasets of SD2-2 lies in estimating the distribution of state sojourn time. ISHSMM and ISFLS can accurately assess the Poisson distribution of state sojourn time. They are followed by ISSWFC and ISGLKF. ISHMM, ISDCC, and ISMTD perform the worst. The original versions of these methods perform similarly to the enhanced versions, but the state transition matrix exhibits poorer inter-data consistency.

Figures 12 and 13 present the estimated state maps and state transition vectors for SD2-2-2. Figure 14 shows the fitted state sojourn distribution for SD2-2-2. Figures 15 and 16 display the spatial and temporal correlation results for SD2-2-2, and Figure 17 presents the Frechet distances between the predefined and estimated state sojourn time density functions. In calculating the state maps, ISHSMM, ISHMM, and ISFLS perform the best, followed by ISDCC, ISSWFC, ISMTD, and ISGLKF. ISHSMM and ISFLS also perform best when estimating the temporal properties of state transitions, followed by ISSWFC and ISGLKF, with ISDCC, ISMTD, and ISHMM performing the worst. Similar results were observed for SD2-2-1 (Supplementary Figures 21–26) and SD2-2-3 (Supplementary Figures 27–32).

**Figure 12.**
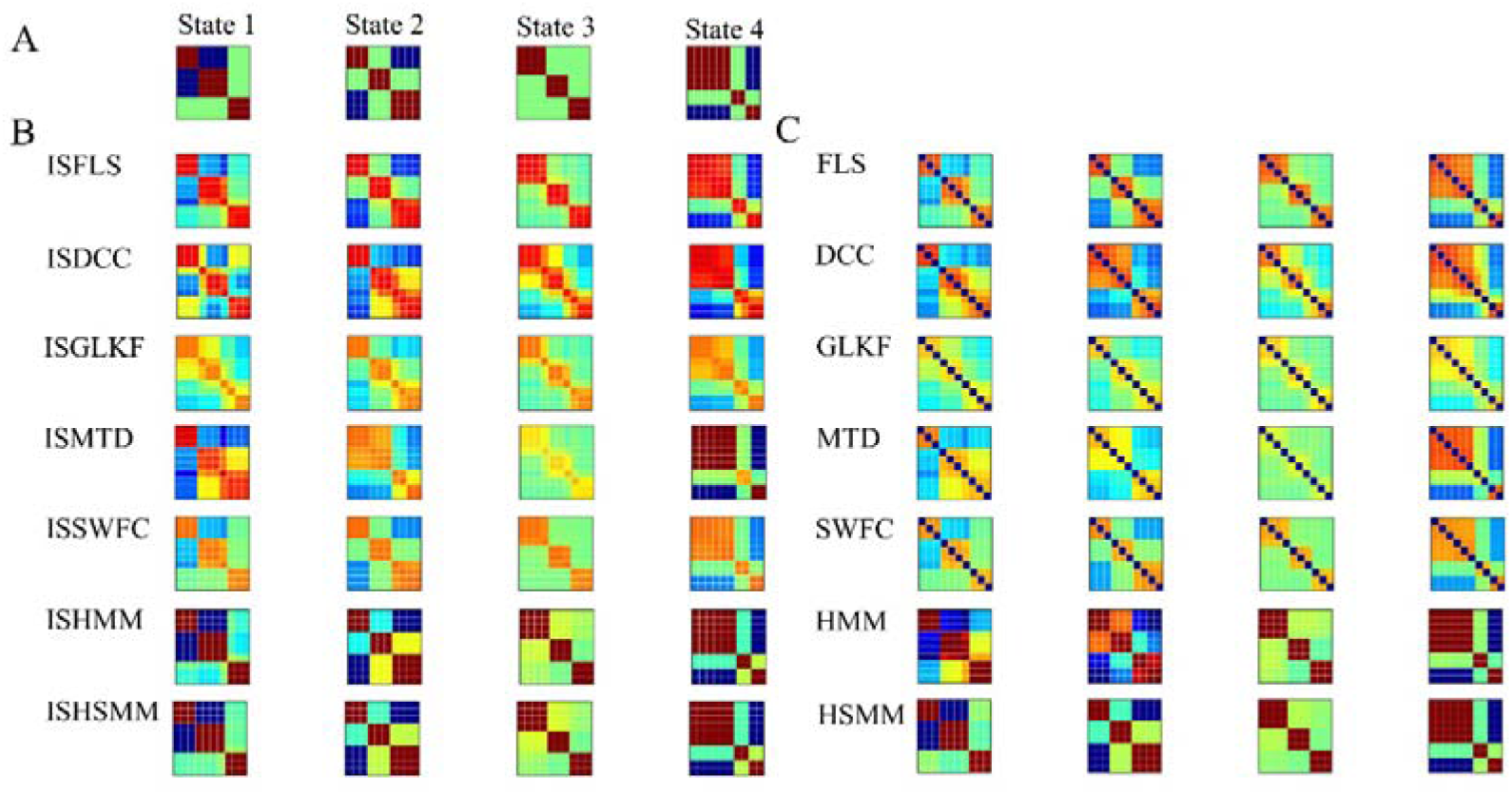
The predefined and estimated states of SD2-2-2 were determined using the 14 dFC methods.

**Figure 13.**
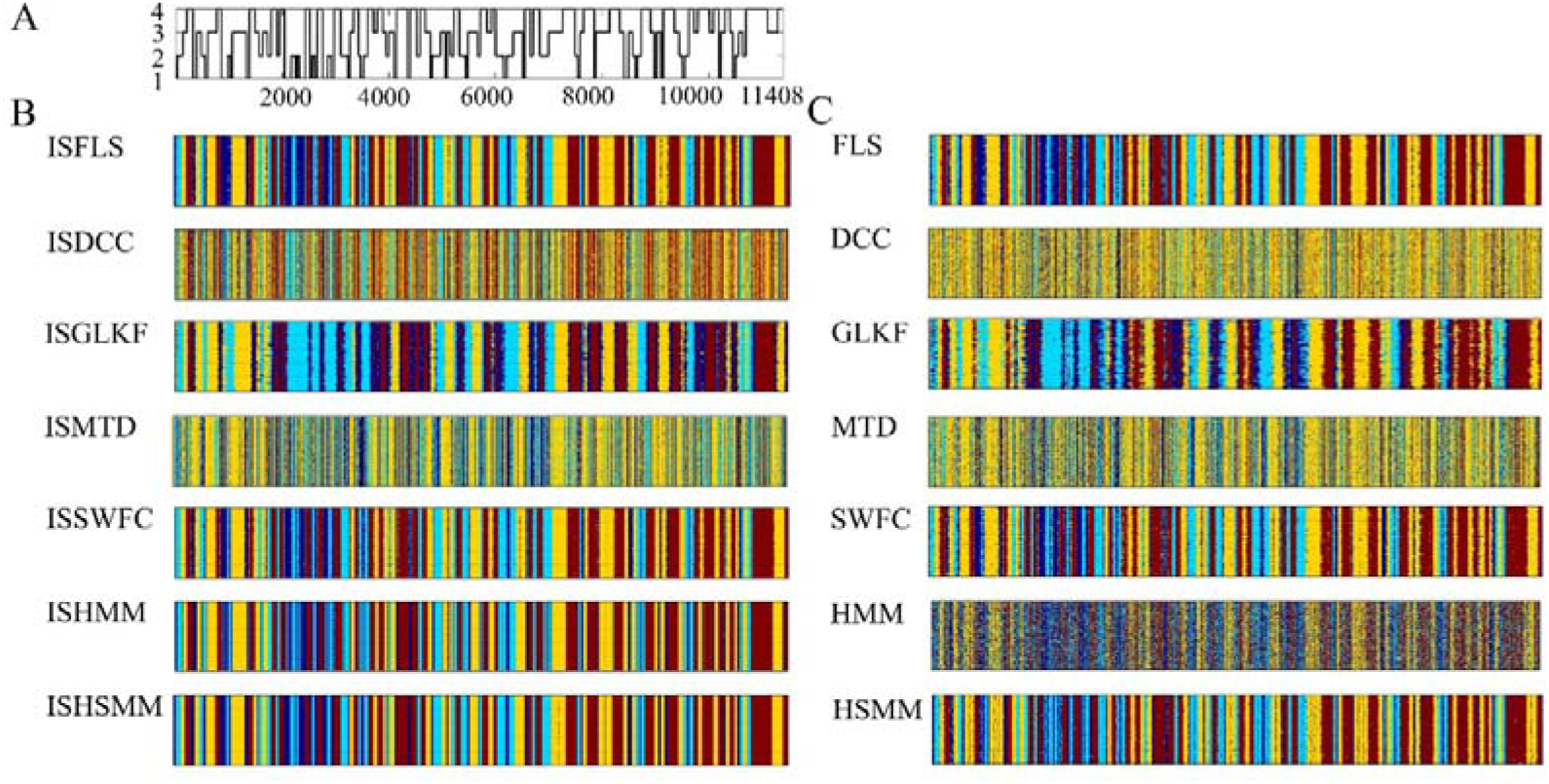
The estimated state transition vectors of SD2-2-2 by the 14 dFC methods. A: The predefined state transition vector of the four states.

**Figure 14.**
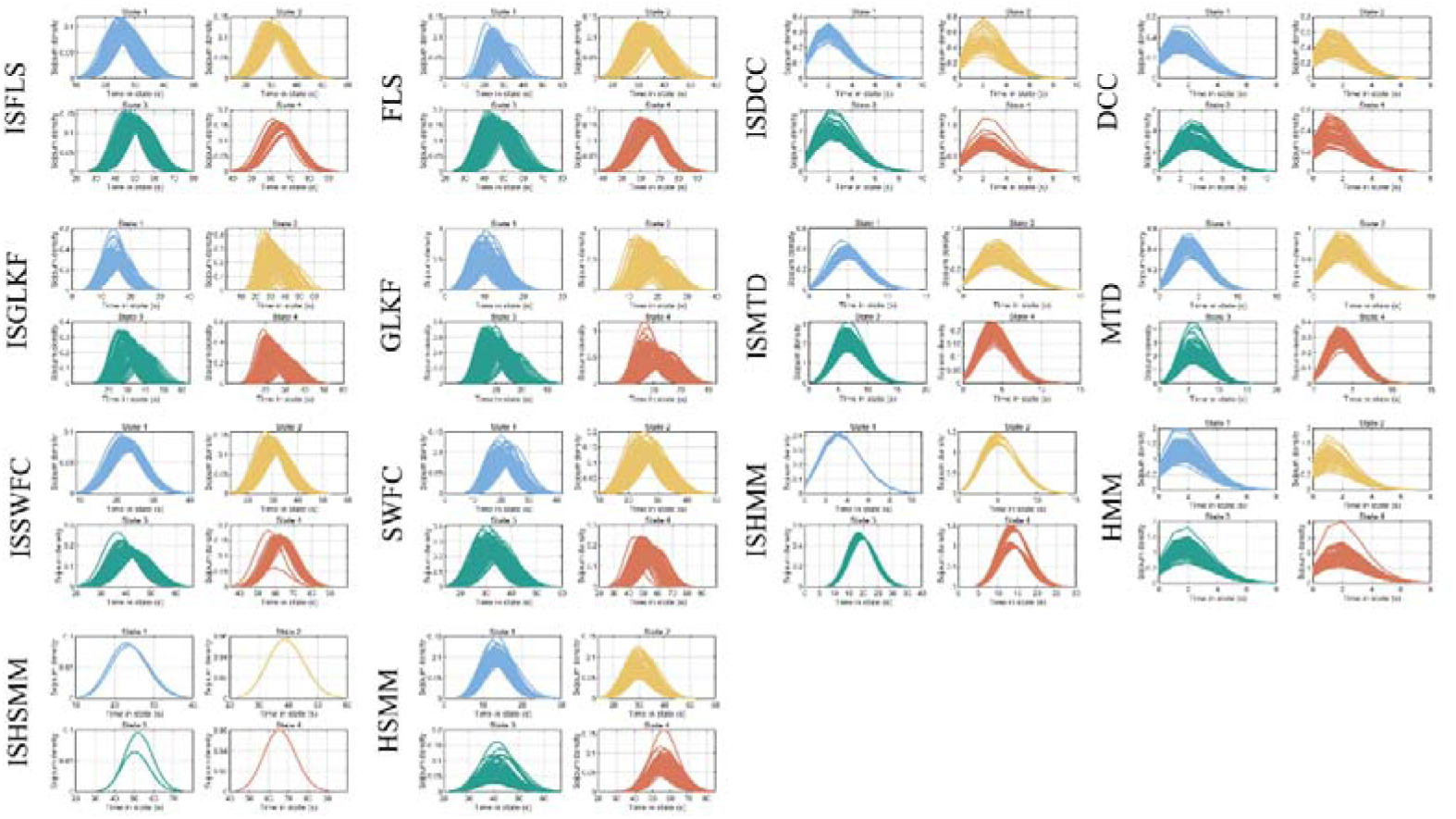
The distributions of state sojourn time of SD2-2-2 by the 14 dFC methods. In SD2-22, Poisson-distributed state sojourns (λ = 20, 30, 40, and 50 for states 1, 2, 3, and 4, respectively) with Gaussian noise (SD = 0.3) were simulated.

**Figure 15.**
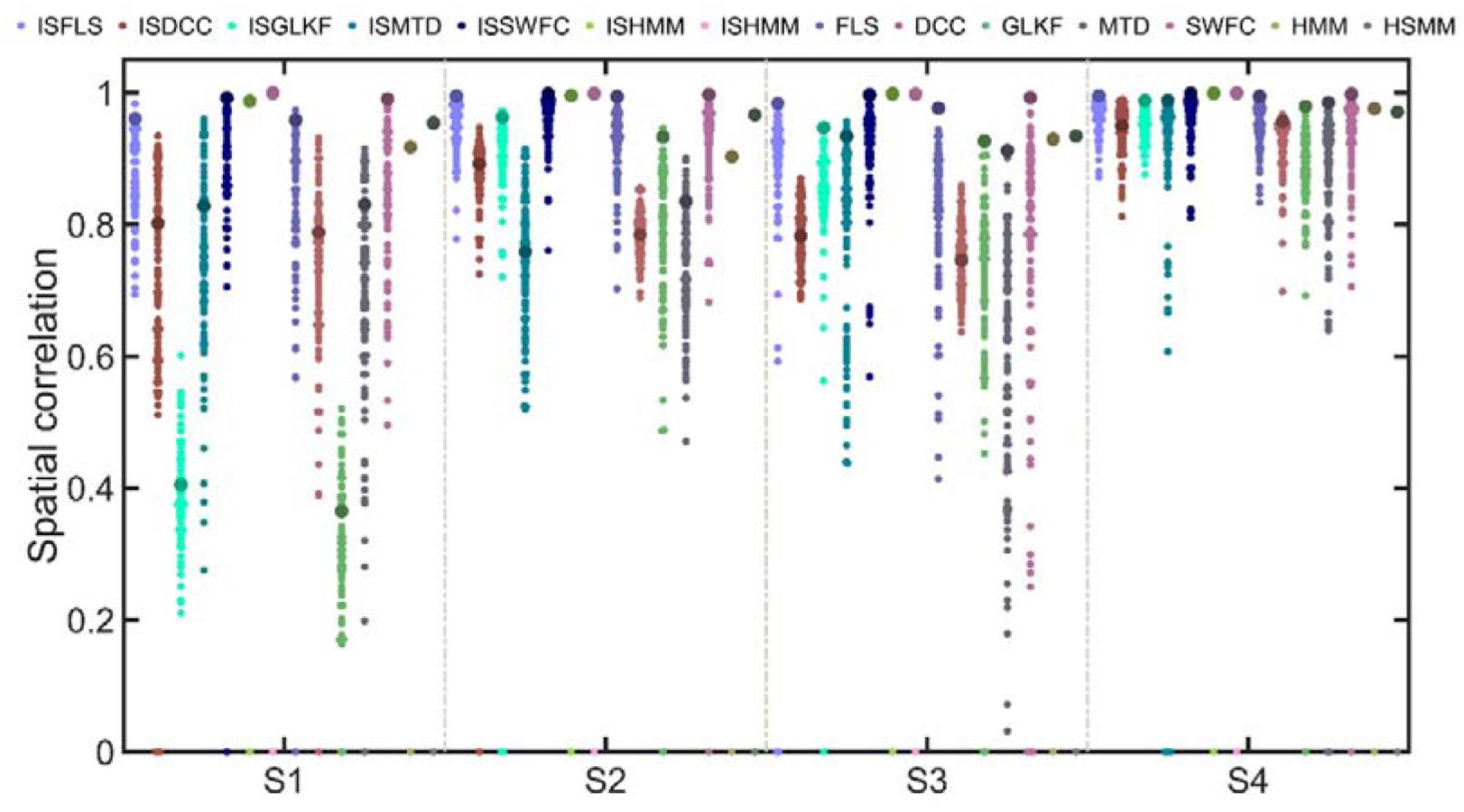
The spatial correlations of SD2-2-2 between the predefined and estimated states by the 14 dFC methods.

**Figure 16.**
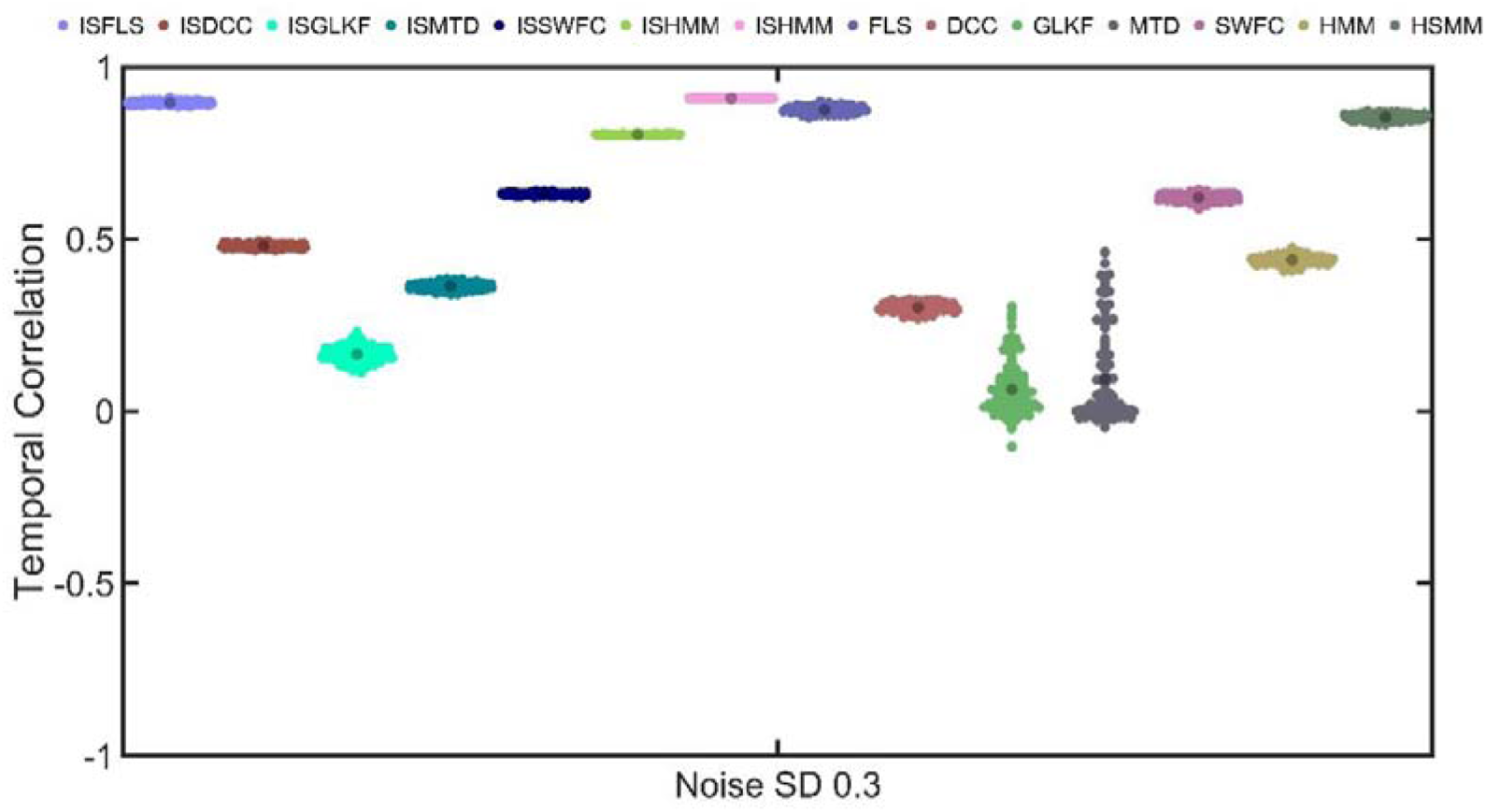
The temporal correlations of SD2-2-2 between the predefined state transition vectors and the estimated state transition vectors by the 14 dFC methods.

**Figure 17.**
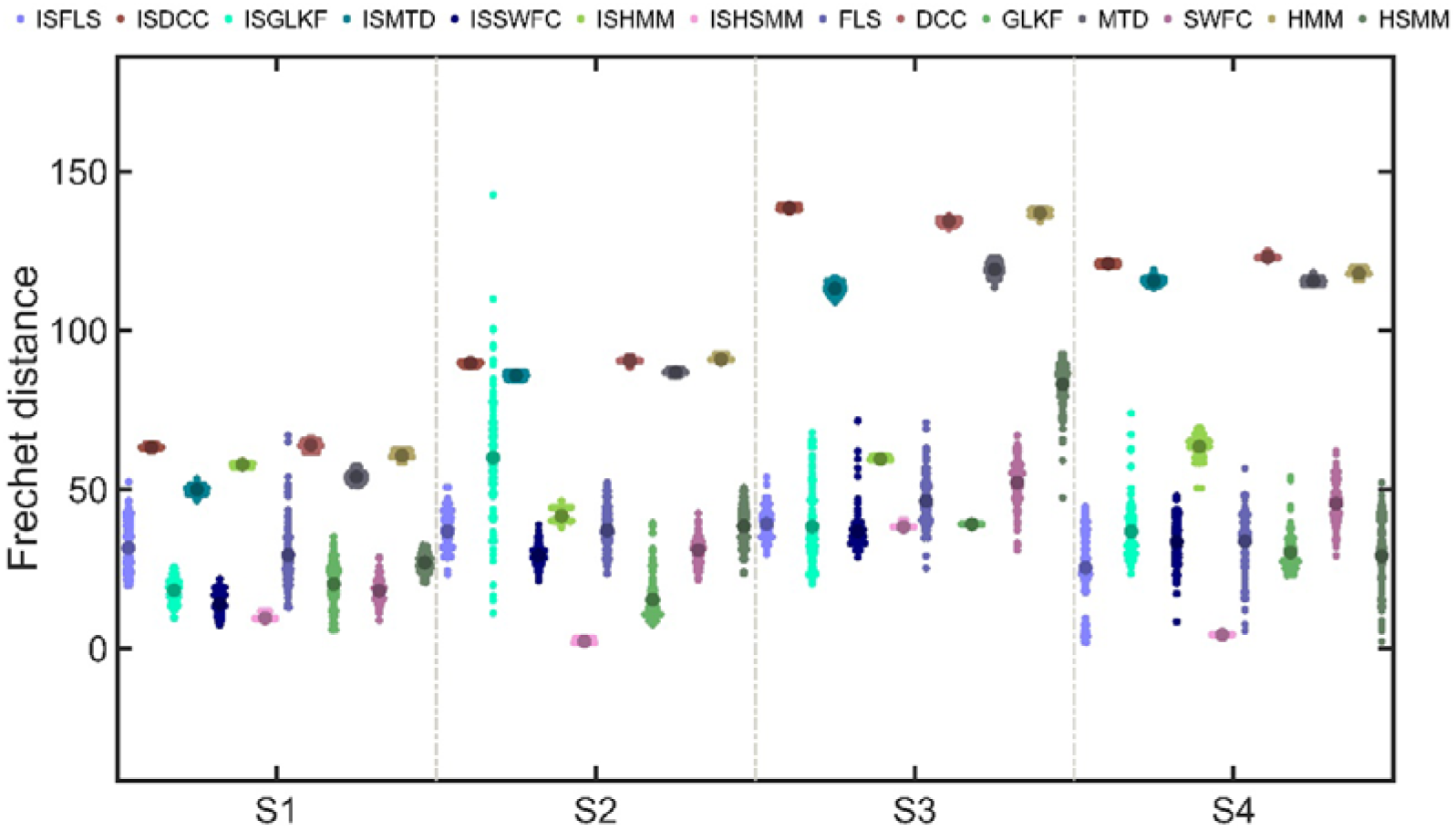
The Frechet distances between the predefined state sojourn time density curve and the estimated state sojourn time density curve for SD2-2-2, calculated using the 14 dFC methods.

### The effect of neurovascular effect in simulated data

As shown in Figures 9 and 13, DCC was sensitive to state transition. One possibility is that the procedure of convolution with HRF might have altered the covariance structure between the signals. To demonstrate this, we modified the simulation approach for SD2-1-2 and SD2-2-2 by not performing convolution after simulating signals with specific connectivity patterns and state durations. After adding Gaussian noise with a standard deviation of 0.3, we conducted dFC analysis on these new datasets (SD2-1-2-NoHRF and SD2-2-2-NoHRF) using ISDCC and DCC. The results showed that ISDCC accurately estimated the four connectivity states and their state sojourn times (Supplementary Figures 33 and 34), which are very close to those of ISHMM.

## Discussion

In this study, we presented a systemic comparison of seven dFC methods (and their enhanced versions) in tracking the time-varying connectivity. Given the absence of ground truth in real fMRI, we developed various simulation data with spatiotemporal benchmarks. Considering that several methods were initially proposed for non-fMRI-BOLD signals, thus both non-fMRI-BOLD and fMRI-BOLD signals with different covariance structures and sojourn distributions were simulated. For simulated non-fMRI-BOLD signals, the covariance of the two time series included three types of distributions: zero distribution, Gaussian distribution, and periodic distribution. For simulated fMRI-BOLD signals, four connectivity patterns among 10 regions were simulated with two types of state sojourn distributions (one with fixed sojourn duration and one with Poisson distribution. To make the simulated signals more realistic, we added spontaneous activity signals to the fMRI-BOLD signals. We also introduced Gaussian noise signals to the non-fMRI-BOLD and fMRI-BOLD signals. To suppress the effect of spontaneous activity signals and Gaussian noise signals and increase the sensitivity to the signal of interest, we got the enhanced versions of the seven methods by adopting ISA. The efficacy was defined as the spatial and temporal associations between the predefined and estimated covariance structures and connectivity states. To our knowledge, this is the most comprehensive study of dFC methodologies to date. Our results show that all factors we considered will affect the efficacy, including whether considering the neurovascular effect in simulated data, the covariance structure between two time series, state sojourn distribution, and levels of SNR.

Some of the results in this study are consistent with previous research. For example, consistent with the findings of Lindquist et al. (2014), DCC (and ISDCC) was able to accurately estimate the actual value for two uncorrelated non-fMRI-BOLD time series (SD1-1), while SWFC was often given poor estimation and was susceptible to rapid changes of covariance (SD1-1, SD1-2, SD1-3). Similarly, consistent with the findings of Shappell et al. (2019), HSMM and its enhanced version could accurately estimate state sojourn times with Poisson distributions (SD2-2) and were not susceptible to noise, while HMM failed due to the inherent limitation of geometrical distribution.

Our results indicate that the efficacy of dFC methods is related to various factors. In most cases, ISFSL and FSL performed excellently on non-fMRI-BOLD and fMRI-BOLD data but failed in SD1-1 and SD2-1. The superior performance of FSL can be attributed to its consideration of both residual measurement error (the discrepancy between the current observation and the value estimated by the current regression model) and residual dynamic error (the discrepancy between the current and previous regression coefficients). However, FSL is initially proposed for data generated by a linear regression model with coefficients that evolve slowly over time (Kalaba and Tesfatsion, 1989). Thus, when the two time series were uncorrelated (SD1-1), i.e., no linear regression model could be estimated, FSL may give poor estimation (Fig.2). When the covariance remained unchanged for a long period of time and presented an unanticipated shift at a time (SD2-1), the estimates of FSL would monotonically increase or decrease between the two coefficients of two specific times. The FSL estimates would firstly increase (or decrease) at an increasing rate over the initial time points, reach a maximum speed at the moment of covariance change, and a decreasing rate when reaching the final estimates (See Figure 2 in Kalaba and Tesfatsion, 1989). For this reason, in the data of SD2-1, FSL provided an incorrect state transition vector (Fig.9).

DCC and ISDCC performed significantly better on non-fMRI-BOLD data compared to fMRI-BOLD data. The convolution with HRF altered the covariance structure between the signals, leading to erroneous estimates. These findings explain why DCC method performs less favorably than other dFC methods in real fMRI data (Xie et al., 2019).

As the only one proposed for fMRI data, MTD (and ISMTD) performed worst in most cases and were susceptible to noise. Due to the differencing of the time series for MTD calculation, MTD has a high-pass filtering effect. Thus, it is insensitive to signal changes in the lower frequency bands and will underestimate the covariance below 0.05 Hz (see Supplementary Figure 3 in Shine et al., 2015). For all data in SD1, the power spectra of the two time series are evenly distributed between 0 and 0.5 Hz (Supplementary Figure 35), while for all data in SD2, similar to real fMRI-BOLD data, they exhibit low-frequency characteristics, with power spectra for all time series following an exponential distribution, primarily concentrated between 0 and 0.05 Hz (Supplementary Figure 36). These characteristics of simulated data cause the MTD method to erroneously estimate the covariance structure of the two time series.

Temporal attributes are crucial outcomes in dFC research. However, our findings reveal that some algorithms frequently produce erroneous estimates of state sojourn times (DCC and HMM), challenging the reliability of these results (Chen et al., 2022; Song et al., 2021b; Yuan et al., 2023a; Yuan et al., 2023b). Moreover, our study highlights the necessity of incorporating ISA method, which enhance dFC efficacy and reliability. In datasets with multiple components, such as naturalistic fMRI-BOLD data, these enhanced methods can effectively suppress irrelevant signals and increase sensitivity to signals of interest (Simony et al., 2016). Additionally, noise signals significantly impact dFC efficacy, diminishing the accuracy of state and temporal attribute estimates and increasing the coefficient of variation across subjects. This emphasizes the importance of implementing robust preprocessing techniques to isolate and eliminate noise signals (Yan et al., 2013; Yuan et al., 2016).

This study has several limitations. Firstly, although various datasets were simulated, the simulation method has inherent constraints. For instance, due to convolution with the hemodynamic response function, rapid changes in fMRI-BOLD data cannot be accurately simulated, potentially affecting the assessment of certain frame-wise methods. Additionally, convolution with HRF may alter the predefined covariance structures among signals and change the temporal resolution and signal amplitudes. Therefore, the findings of this study should be interpreted cautiously. Future research should prioritize validating the accuracy and generalizability of these results using real fMRI-BOLD data.

Secondly, other gold standard methods exist to assess dFC efficacy apart from simulation methods. These include coupling analysis of neural activity and hemodynamics (Ma et al., 2016) using multimodal neuroimaging data such as local field potential (via iEEG) and fMRI data (Berezutskaya et al., 2022; Keles et al., 2024), brain-body association analysis through synchronized physiological recording (e.g., skin conductance and eye movement data) with fMRI (Gamer et al., 2007; Williams et al., 2001), or brain-behavior association analysis through simultaneous recordings of subjects’ behavioral responses (e.g., attentional engagement) (Song et al., 2021a).

## Supporting information

Supplementary methods and results

## Data and code availability statement

Table 1 summarizes code and software availability of the seven methods. The in-house codes are available at https://github.com/yuanbinke/dFC_Simu.

## References

Allen, E.A., Damaraju, E., Plis, S.M., Erhardt, E.B., Eichele, T., Calhoun, V.D., 2014. Tracking whole-brain connectivity dynamics in the resting state. Cereb Cortex 24, 663–676.

Berezutskaya, J., Vansteensel, M.J., Aarnoutse, E.J., Freudenburg, Z.V., Piantoni, G., Branco, M.P., Ramsey, N.F., 2022. Open multimodal iEEG-fMRI dataset from naturalistic stimulation with a short audiovisual film. Scientific data 9, 91.

Calhoun, V.D., Miller, R., Pearlson, G., Adali, T., 2014. The chronnectome: time-varying connectivity networks as the next frontier in fMRI data discovery. Neuron 84, 262–274.

Chen, K., Li, C., Sun, W., Tao, Y., Wang, R., Hou, W., Liu, D.Q., 2022. Hidden Markov Modeling Reveals Prolonged "Baseline" State and Shortened Antagonistic State across the Adult Lifespan. Cereb Cortex 32, 439–453.

Choe, A.S., Nebel, M.B., Barber, A.D., Cohen, J.R., Xu, Y., Pekar, J.J., Caffo, B., Lindquist, M.A., 2017. Comparing test-retest reliability of dynamic functional connectivity methods. Neuroimage 158, 155–175.

Cohen, J.R., 2018. The behavioral and cognitive relevance of time-varying, dynamic changes in functional connectivity. Neuroimage 180, 515–525.

Damaraju, E., Tagliazucchi, E., Laufs, H., Calhoun, V.D., 2020. Connectivity dynamics from wakefulness to sleep. Neuroimage 220, 117047.

Di, X., Biswal, B.B., 2020. Intersubject consistent dynamic connectivity during natural vision revealed by functional MRI. Neuroimage 216, 116698.

Eavani, H., Satterthwaite, T.D., Gur, R.E., Gur, R.C., Davatzikos, C., 2013. Unsupervised learning of functional network dynamics in resting state fMRI. Information processing in medical imaging : proceedings of the … conference 23, 426–437.

Eiter, T., Mannila, H., 1994. Computing discrete Fréchet distance.

Engle, R., 2002. Dynamic conditional correlation: A simple class of multivariate generalized autoregressive conditional heteroskedasticity models. J Bus Econ Stat 20, 339–350.

Gamer, M., Bauermann, T., Stoeter, P., Vossel, G., 2007. Covariations among fMRI, skin conductance, and behavioral data during processing of concealed information. Hum Brain Mapp 28, 1287–1301.

Gonzalez-Castillo, J., Hoy, C.W., Handwerker, D.A., Robinson, M.E., Buchanan, L.C., Saad, Z.S., Bandettini, P.A., 2015. Tracking ongoing cognition in individuals using brief, whole-brain functional connectivity patterns. Proc Natl Acad Sci U S A 112, 8762–8767.

Guédon, Y., 2003. Estimating hidden semi-Markov chains from discrete sequences. Journal of computational and graphical statistics 12, 604–639.

Hutchison, R.M., Womelsdorf, T., Allen, E.A., Bandettini, P.A., Calhoun, V.D., Corbetta, M., Della Penna, S., Duyn, J.H., Glover, G.H., Gonzalez-Castillo, J., Handwerker, D.A., Keilholz, S., Kiviniemi, V., Leopold, D.A., de Pasquale, F., Sporns, O., Walter, M., Chang, C., 2013. Dynamic functional connectivity: promise, issues, and interpretations. Neuroimage 80, 360–378.

Kalaba, R., Tesfatsion, L., 1989. Time-varying linear regression via flexible least squares. Computers & Mathematics with Applications 17, 1215–1245.

Kang, J., Wang, L., Yan, C., Wang, J., Liang, X., He, Y., 2011. Characterizing dynamic functional connectivity in the resting brain using variable parameter regression and Kalman filtering approaches. Neuroimage 56, 1222–1234.

Keles, U., Dubois, J., Le, K.J.M., Tyszka, J.M., Kahn, D.A., Reed, C.M., Chung, J.M., Mamelak, A.N., Adolphs, R., Rutishauser, U., 2024. Multimodal single-neuron, intracranial EEG, and fMRI brain responses during movie watching in human patients. Scientific data 11, 214.

Liao, W., Wu, G.R., Xu, Q., Ji, G.J., Zhang, Z., Zang, Y.F., Lu, G., 2014. DynamicBC: a MATLAB toolbox for dynamic brain connectome analysis. Brain connectivity 4, 780–790.

Lindquist, M.A., Meng Loh, J., Atlas, L.Y., Wager, T.D., 2009. Modeling the hemodynamic response function in fMRI: efficiency, bias and mis-modeling. Neuroimage 45, S187–198.

Lindquist, M.A., Xu, Y., Nebel, M.B., Caffo, B.S., 2014. Evaluating dynamic bivariate correlations in resting-state fMRI: a comparison study and a new approach. Neuroimage 101, 531–546.

Ma, Y., Shaik, M.A., Kozberg, M.G., Kim, S.H., Portes, J.P., Timerman, D., Hillman, E.M., 2016. Resting-state hemodynamics are spatiotemporally coupled to synchronized and symmetric neural activity in excitatory neurons. Proc Natl Acad Sci U S A 113, E8463–E8471.

Milde, T., Leistritz, L., Astolfi, L., Miltner, W.H., Weiss, T., Babiloni, F., Witte, H., 2010. A new Kalman filter approach for the estimation of high-dimensional time-variant multivariate AR models and its application in analysis of laser-evoked brain potentials. Neuroimage 50, 960–969.

O’Connell, J., Højsgaard, S., 2011. Hidden semi markov models for multiple observation sequences: The mhsmm package for R. Journal of Statistical Software 39, 1–22.

Satterthwaite, T.D., Elliott, M.A., Gerraty, R.T., Ruparel, K., Loughead, J., Calkins, M.E., Eickhoff, S.B., Hakonarson, H., Gur, R.C., Gur, R.E., Wolf, D.H., 2013. An improved framework for confound regression and filtering for control of motion artifact in the preprocessing of resting-state functional connectivity data. Neuroimage 64, 240–256.

Shakil, S., Lee, C.H., Keilholz, S.D., 2016. Evaluation of sliding window correlation performance for characterizing dynamic functional connectivity and brain states. Neuroimage 133, 111–128.

Shappell, H., Caffo, B.S., Pekar, J.J., Lindquist, M.A., 2019. Improved state change estimation in dynamic functional connectivity using hidden semi-Markov models. Neuroimage 191, 243–257.

Shine, J.M., Koyejo, O., Bell, P.T., Gorgolewski, K.J., Gilat, M., Poldrack, R.A., 2015. Estimation of dynamic functional connectivity using Multiplication of Temporal Derivatives. Neuroimage 122, 399–407.

Simony, E., Honey, C.J., Chen, J., Lositsky, O., Yeshurun, Y., Wiesel, A., Hasson, U., 2016. Dynamic reconfiguration of the default mode network during narrative comprehension. Nature communications 7, 12141.

Song, H., Finn, E.S., Rosenberg, M.D., 2021a. Neural signatures of attentional engagement during narratives and its consequences for event memory. Proc Natl Acad Sci U S A 118.

Song, H., Park, B.Y., Park, H., Shim, W.M., 2021b. Cognitive and Neural State Dynamics of Narrative Comprehension. J Neurosci 41, 8972–8990.

Thompson, W.H., Fransson, P., 2018. A common framework for the problem of deriving estimates of dynamic functional brain connectivity. Neuroimage 172, 896–902.

Thompson, W.H., Richter, C.G., Plaven-Sigray, P., Fransson, P., 2018. Simulations to benchmark time-varying connectivity methods for fMRI. PLoS computational biology 14, e1006196.

Vergara, V.M., Mayer, A.R., Damaraju, E., Calhoun, V.D., 2017. The effect of preprocessing in dynamic functional network connectivity used to classify mild traumatic brain injury. Brain and behavior 7, e00809.

Vidaurre, D., Smith, S.M., Woolrich, M.W., 2017. Brain network dynamics are hierarchically organized in time. Proc Natl Acad Sci U S A 114, 12827–12832.

Weissenbacher, A., Kasess, C., Gerstl, F., Lanzenberger, R., Moser, E., Windischberger, C., 2009. Correlations and anticorrelations in resting-state functional connectivity MRI: a quantitative comparison of preprocessing strategies. Neuroimage 47, 1408–1416.

Williams, L.M., Phillips, M.L., Brammer, M.J., Skerrett, D., Lagopoulos, J., Rennie, C., Bahramali, H., Olivieri, G., David, A.S., Peduto, A., Gordon, E., 2001. Arousal dissociates amygdala and hippocampal fear responses: evidence from simultaneous fMRI and skin conductance recording. Neuroimage 14, 1070–1079.

Xie, H., Zheng, C.Y., Handwerker, D.A., Bandettini, P.A., Calhoun, V.D., Mitra, S., Gonzalez-Castillo, J., 2019. Efficacy of different dynamic functional connectivity methods to capture cognitively relevant information. Neuroimage 188, 502–514.

Yan, C.G., Cheung, B., Kelly, C., Colcombe, S., Craddock, R.C., Di Martino, A., Li, Q., Zuo, X.N., Castellanos, F.X., Milham, M.P., 2013. A comprehensive assessment of regional variation in the impact of head micromovements on functional connectomics. Neuroimage 76, 183–201.

Yu, S.-Z., 2010. Hidden semi-Markov models. Artificial intelligence 174, 215–243.

Yuan, B., Xie, H., Gong, F., Zhang, N., Xu, Y., Zhang, H., Liu, J., Chen, L., Li, C., Tan, S., Lin, Z., Hu, X., Gu, T., Cheng, J., Lu, J., Liu, D., Wu, J., Yan, J., 2023a. Dynamic network reorganization underlying neuroplasticity: the deficits-severity-related language network dynamics in patients with left hemispheric gliomas involving language network. Cereb Cortex.

Yuan, B., Xie, H., Wang, Z., Xu, Y., Zhang, H., Liu, J., Chen, L., Li, C., Tan, S., Lin, Z., Hu, X., Gu, T., Lu, J., Liu, D., Wu, J., 2023b. The domain-separation language network dynamics in resting state support its flexible functional segregation and integration during language and speech processing. Neuroimage 274, 120132.

Yuan, B.K., Zang, Y.F., Liu, D.Q., 2016. Influences of Head Motion Regression on High-Frequency Oscillation Amplitudes of Resting-State fMRI Signals. Frontiers in human neuroscience 10, 243.

