## Supplementary methods and results for "A systematic evaluation of dynamic functional connectivity methods using simulation data"

Sliding-window functional connectivity with L1-regularization (SWFC)

Given two windowed time series $x_{i}(t)$ and $y_{i}(t)$, the SWFC is:

$SWFC=\frac{\sum_{T-wl/2}^{T+wl/2} (x_{i}(t)-\bar{x_{i}})(y_{i}(t)-\bar{y_{i}})}{\sqrt{\sum_{T-wl/2}^{T+wl/2} {(x_{i}(t)-\bar{x_{i}})}^{2}}\sqrt{\sum_{T-wl/2}^{T+wl/2} {(y_{i}(t)-\bar{y_{i}})}^{2}}}$	(1)

Where$\bar{x_{i}}$ and $\bar{y_{i}}$ are the sample mean of windowed time series, and $wl$ denotes the window length.

Similar to other studies, we used a Gaussian tapered window by convolving a rectangle with a Gaussian kernel ($\sigma= 3$).

The small samples per window may lead to instable parameter estimate. L1-regularization with graphical LASSO is often adopted with assumption that the estimated functional connectivity matrix is sparse. Instead of assuming correlation

matrices being sparse, graphical LASSO assumes the inverse covariance matrix $\Phi$ to be sparse. Graphical LASSO gets the sparse solution of the inverse covariance matrix $\Phi$ by maximizing the following log-likelihood function:

$L_{1}=log det \Phi-tr(s\Phi)- \lambda\left\| \Phi\right\|_{1}$	(2)

where $\det$ denotes the determinant, $\mathrm{tr}$ denotes the trace which is the sum of all elements on the main diagonal, s is the empirical covariance matrix, $\lambda$ is the regularization parameter, and $\left\| \Phi\right\|_{1}$ denotes the L1 penalty on $\Phi$. The optimal $\lambda$ is selected through cross-validation, which estimates the optimal $\lambda$ for each subject by evaluating how well the inverse covariance matrix of a training set estimated with a given $\lambda$ describes a test set from the same subject.

Multiplication of temporal derivatives (MTD)

We first calculate the temporal derivative (*dt*) of each time series by performing a first-order differencing:

${dt}_{x}=x_{t}{-x}_{t-1}$	(3)

The MTD score at each time point is equal to the product of the *dt_x_* and *dt_y_*, and then normalized it by dividing each *dt* by the standard deviation (*σ*) of the *dt*:

$MTD=\frac{{dt}_{x} *{dt}_{y}}{\sigma_{x}*\sigma_{y}}$	(4)

To decrease the susceptibility to noise, a simple moving average method is adopted by averaging MTD scores surrounding a point in time within a window:

$SMAMTD=\frac{1}{2\omega+1}\sum_{t-\omega}^{t-\omega} MTD$	(5)

Dynamic conditional correlation (DCC)

Dynamic conditional correlation (DCC) is based on the generalized autoregressive conditional heteroscedastic model (GARCH). Bollerslev (Bollerslev, 1986) proposed the GARCH model, which was a natural generalization of ARCH (Engle, 1982). The main idea of this method is to represent the conditional variance of a single time series at some moment as a linear combination of the past conditional variance and the values of the past time series. For a GARCH (*p*, *q*) process, we assume that $y_{t}$ is a univariate time series with a conditional variance.

$y_{t}\sim N(0,\delta_{t}^{2})$	(6)

where the conditional variance $\delta_{t}^{2}$ is expressed as:

$\delta_{t}^{2}=\alpha_{0}+\sum_{i=1}^{q} \alpha_{i}y_{t-i}^{2}+\sum_{i=1}^{p} \beta_{i}\delta_{t-i}^{2} {where \alpha}_{0}>0,\alpha,\beta\geq0,\alpha+\beta<1$	(7)

The parameter $\alpha_{0}$ is a constant, $\alpha_{i}$ controls the impact of the past values of the time series, and $\beta_{i}$ controls the effects of the past conditional variance of the time series. Then GARCH(1,1) can be written as:

$\delta_{t}^{2}=\alpha_{0}+\alpha y_{t-1}^{2}+\beta\delta_{t-1}^{2}$	(8)

To illustrate the DCC model, we assume that $y_{t}$ is a bivariate time series with a covariance matrix $\Sigma_{t}$. $y_{t}$ contains two univariate time series.

$y_{t}\sim N(0,\Sigma_{t})$	(9)

The DCC method consists of two steps. First, the univariate GARCH(1, 1) model is fit to the case of two univariate time series of $y_{t}$. In the first step, Eq.(10) ~ Eq.(11) are used to estimate the standard residuals $\epsilon_{t}$.

$\delta_{i,t}^{2}=\alpha_{i,0}+\alpha y_{i,t-1}^{2}+\beta\delta_{i,t-1}^{2} for i=1, 2$	(10)

$D_{t}=diag\{\delta_{1,t},\delta_{2,t}\}$	(11)

$\epsilon_{t}={D_{t}}^{-1}y_{t}$	(12)

Second, the standardized residuals $\epsilon_{t}$(Eq.(12)) were used to estimate the covariance matrix to calculate the dynamic correlation. The $\overline{Q}$ is the non-conditional covariance matrix of $\epsilon_{t}$ (Eq.(13)). The covariance matrix is estimated after a simple scaling. The upper right corner of the covariance matrix is the correlation coefficient of the two univariate time series.

$Q_{t}=\left( 1-\theta_{1}-\theta_{2} \right)\overline{Q}+\theta_{1}\epsilon_{t-1}\epsilon_{t-1}^{T}+\theta_{2}Q_{t-1} where 0<\theta_{1}+\theta_{2}<1$	(13)

$R_{t}=diag{\{Q_{t}\}}^{-1/2}Q_{t}{\{Q_{t}\}}^{-1/2}$	(14)

$\Sigma_{t}=D_{t}R_{t}D_{t}$	(15)

DCC has been proven to outperform other dynamic methods to unveil reliable dynamic functional correlations (Choe et al., 2017; Lindquist et al., 2014), and has been used to investigate the spatiotemporal properties of language network in resting state (Yuan et al., 2023b, 2023a).

Flexible Least Squares (FLS)

Given two windowed time series $x_{i}(t)$ and $y_{i}(t)$ , FLS supposes that the linear regression coefficients, $\beta(t)$ changes over time,

$y_{i}(t)=x_{i}(t)\beta_{t}+\varepsilon(t)$	(16)

The idea of FLS in estimating $\beta(t)$ is minimizing two types of errors, the residual measurement error and the residual dynamic error. The residual measurement error is defined as:

$r_{m}^{2}(\beta)=\sum_{t=1}^{T} {(y_{i}(t)-x_{i}(t)\beta_{t})}^{2}$	(17)

Where *T* is the length of the time series.

The residual dynamic error is defined as:

$r_{D}^{2}(\beta)=\sum_{t=1}^{T-1} {(\beta(t+1)-\beta(t))}^{T}(\beta(t+1)-\beta(t))$	(18)

To find the residual efficiency frontier, an incompatibility cost is defined as:

$C=\mu r_{D}^{2}(\beta)$+$r_{m}^{2}(\beta)$	(19)

The incompatibility cost function $C$ generalizes the goodness-of-fit criterion function for ordinary least squares estimation by permitting the coefficient vector $\beta(t)$ to vary over time.

General linear Kalman filter

GLKF is an estimator of a system’s state space and covariance. It can be used to reconstruct the set of linearly independent hidden variables that regulate the evolution of the system over time:

$z(t)=(t-1)z(t-1)+\varepsilon(t)$	(20)

In Eq 20, the hidden state $z(t)$ at time t has a deterministic component given by the propagation of the previous state $z(t-1)$ through a transition matrix , and a stochastic component given by the zero-mean white noise sequence $\varepsilon(t)$ of covariance *Q(t)*.

$x_{i}(t)=H(t-1)z(t)+v(t)$	(21)

In Eq 21, the observed data $x_{i}(t)$ at time t are expressed as a linear combination of the state variable x with projection measurement matrix H, in the absence of noise. The term $v(t)$ is a random white noise perturbation (zero mean, covariance *R(t)*) corrupting the measurements.

To recursively estimate the hidden state x at each time (t = t_1_,..,t_T_), the Kalman filter alternates between two steps, the prediction and the update step. In the prediction step, the state and the error covariance are extrapolated as:

$\hat{{z(t)}^{-}}=(t-1)\hat{{z(t-1)}^{+}}$	(22)

${P(t)}^{-}=(t-1){P(t-1)}^{+}{(t)}^{T}+Q(t-1)$	(23)

where $\hat{{z(t)}^{-}}$ and ${P(t)}^{-}$ are the a priori or predicted state and the error covariance at time t, based on the propagation of the previous estimated state and covariance $\hat{{z(t-1)}^{+}}$ and ${P(t-1)}^{+}$ through the transition matrix . The superscript *T* denotes matrix transposition.

In the update step, a posteriori estimates of the state and error covariance are refined

according to:

$K(t)={P(t)}^{-}{(H(t-1){P(t)}^{-}+R(t))}^{-1}$	(24)

$\hat{{z(t)}^{+}}=\hat{{z(t)}^{-}}+K(t)(x_{i}(t)-H(t)\hat{{z(t)}^{-}})$	(25)

${P(t)}^{+}=(I-K(t)H(t)){P(t)}^{-}$	(26)

where *I* is the identity matrix, and $K(t)$ is the Kalman Gain matrix reflecting the relationship between uncertainty in the prior estimate and uncertainty in the measurements. The Kalman Gain thus quantifies the relative reliability of measurements and predictions and determines which one should be given more weight during the update step.

The optimal behavior of the Kalman filter in fMRI and physiological time series is not known. The transition matrix is usually replaced by an identity matrix I, such that the state in Eq (20) follows a first-order random walk model.

GLKF derives *R* recursively from measurement innovations and approximate *Q* as a diagonal weight matrix that determines the rate of change of ${P(t)}^{-}$. $\hat{R}$ is initialized as *I [d x d*] and adaptively updated from the measurement innovations (the pre-update residuals) as:

$\sum_{r} =\frac{{(x_{i}(t)-H(t)\hat{{z(t)}^{-}})}^{T}(x_{i}(t)-H(t)\hat{{z(t)}^{-}})}{N-1}$	(27)

$\hat{R(t)}=\hat{R(t-1)+c(\sum_{r} -\hat{R(t-1)})}$	(28)

where $\sum_{r}$ is the covariance of measurement innovations, *N* is the total number of trials and c (0 < *c* < 1) is a constant across time that regulates the adaptation speed for $\hat{R(t)}$. The constant c determines the trade-off between fast adaptation and smoothness: a small c value adds inertia to the system, reducing the ability to track and to recover from dynamic changes in the true state while a large c value increases the contribution of measurements to each update and the uncertainty associated with ${P(t)}^{-}$.

$\hat{R(t)}$ is computed before the Kalman update to replace the unknown $R(t)$ in the Kalman Gain with

$K(t)={P(t)}^{-}{H(t)}^{T}{(H(t){P(t)}^{-}{H(t)}^{T}+tr(\hat{R(t)})I_{N})}^{-1}$	(29)

where $tr$ denotes the trace of a matrix and $I_{N}$ is the identity matrix [N x N].

Hidden Markov models (HMM) and Hidden semi-Markov models (HSMM)

An HMM describes observed time series as a sequence of hidden states, and the vector of latent network states follows a first order Markov chain:

$f(\tilde{z})=f(z_{1})\prod_{t=2}^{T} f\left( z_{t} | z_{t-1} \right)$	(30)

Where $z_{t}$ is a set of latent states, $\tilde{z}$ is vector of latent segment variables.

The observed BOLD signal vectors at each time point are conditionally independent given the latent process:

$f\left( x_{t} | z_{1:T}, x_{1:t-1},x_{1:t+1} \right)=f\left( x_{t} | z_{t} \right)$	(31)

The complete data log-likelihood:

${(\mu}_{1:K},\sum_{1:K} ,P,\pi)=logf(\tilde{z},\tilde{s})$	(32)

$logf(\tilde{z},\tilde{s})=\sum_{t=1}^{T} logf\left( x_{t} | z_{t} \right)+\sum_{t=2}^{T} logf\left( z_{t} | z_{t-1} \right)+logf(z_{1})$	(33)

The conditional distribution of $x_{t}$ in the given hidden state $z_{t}$ is Gaussian:

$f\left( x_{t} | z_{t}=k \right)\sim N(\mu_{k},\sum k)$	(34)

In HMM, the sojourn time (i.e. number of consecutive time points in a state) distribution for a given state is implicitly geometrically distributed. In HSMM, the underlying latent process is a semi-Markov chain, and the sojourn time is explicitly defined in the model and able to be estimated.

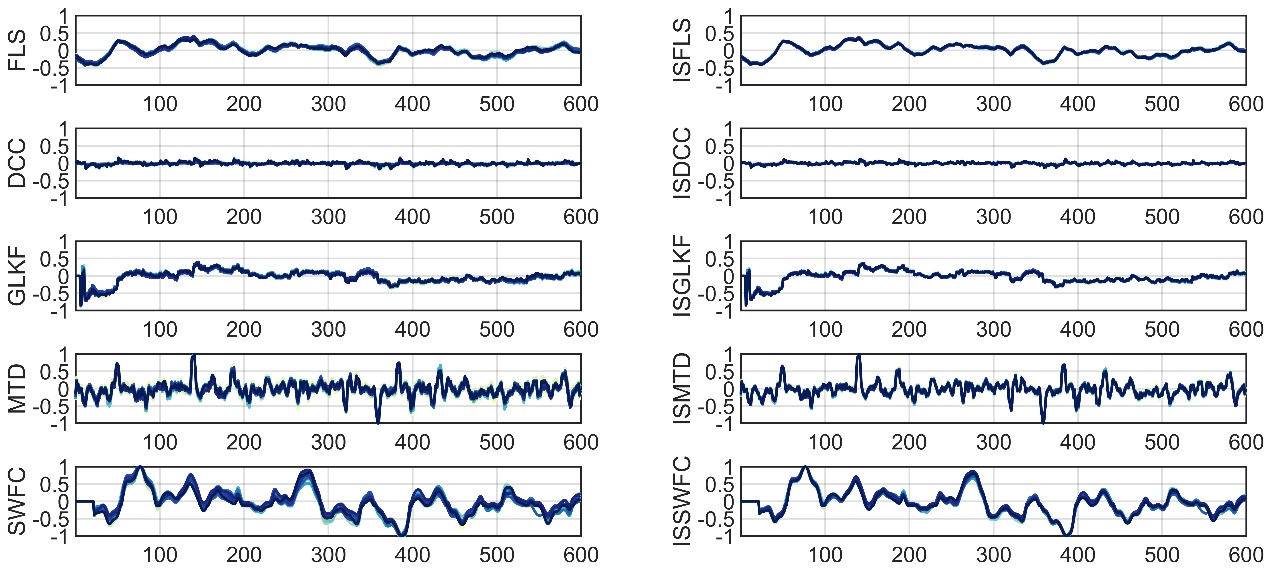

Supplementary Figure 1. Example results of SD1-1-1. The estimated correlation coefficients of two time series with zero covariance from 20 simulations are presented.

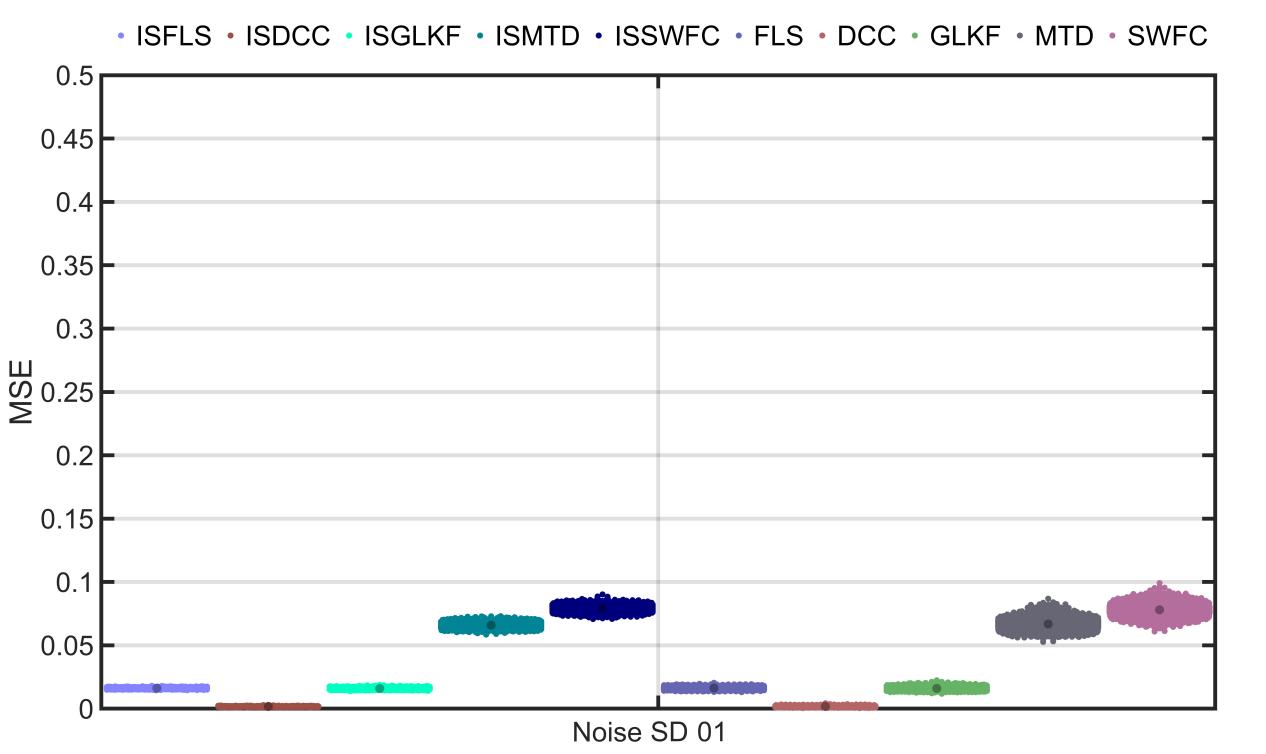

Supplementary Figure 2. MSE results of SD1-1-1. The MSE between the estimated and real correlation coefficients of two time series with zero covariance from 1000 simulations are presented.

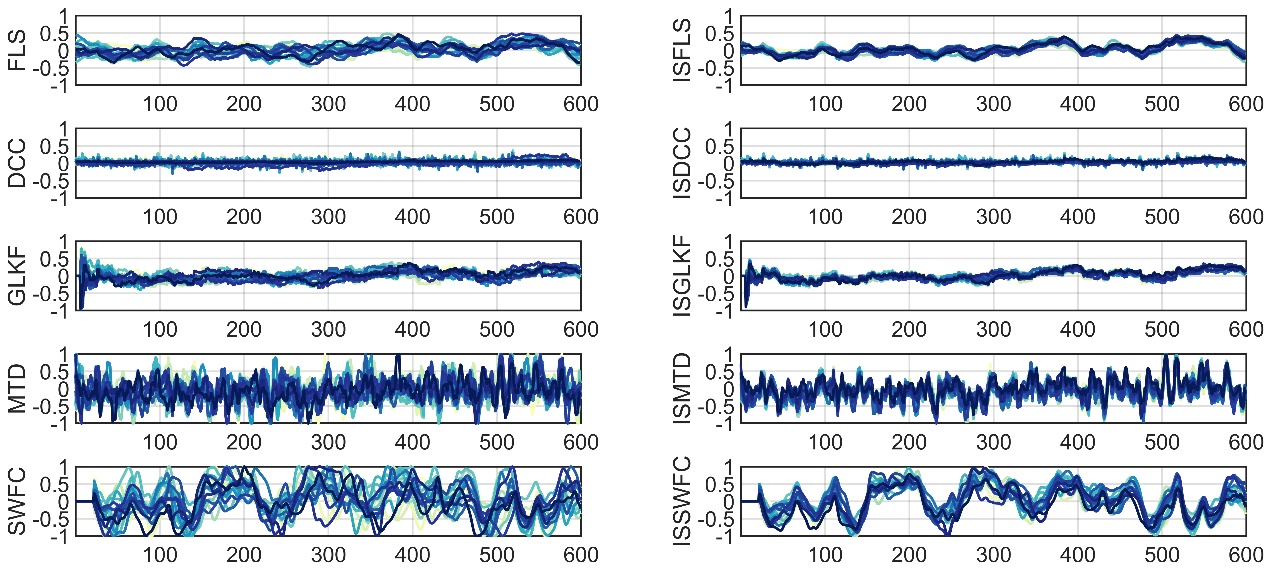

Supplementary Figure 3. Example results of SD1-1-3. The estimated correlation coefficients of two time series with zero covariance from 20 simulations are presented.

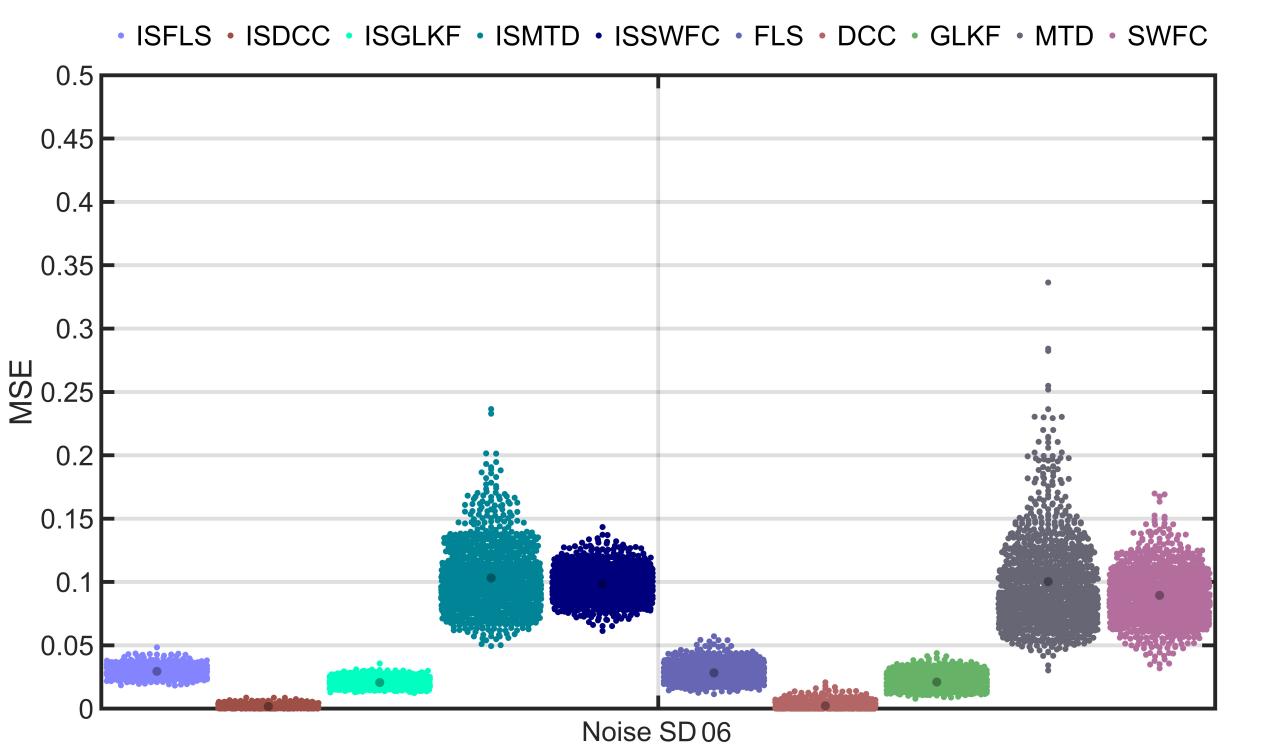

Supplementary Figure 4. MSE results of SD1-1-3. The MSE between the estimated and real correlation coefficients of two time series with zero covariance from 1000 simulations are presented.

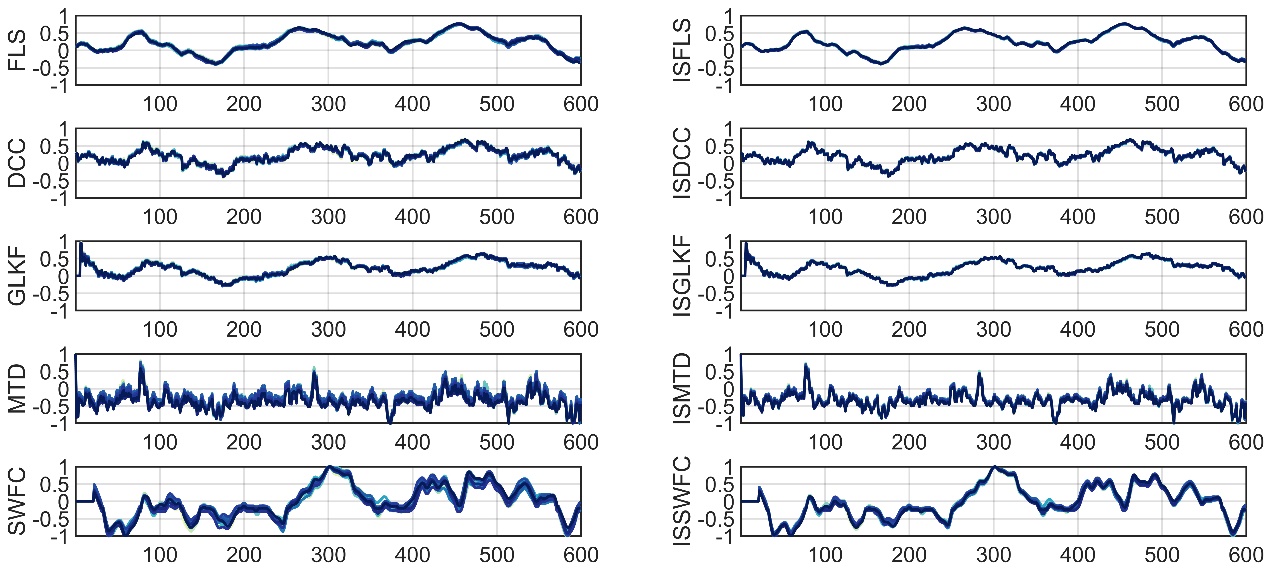

Supplementary Figure 5. Example results of SD1-2-1. The estimated correlation coefficients of two time series with covariance structure of Gaussian distribution from 20 simulations are presented.

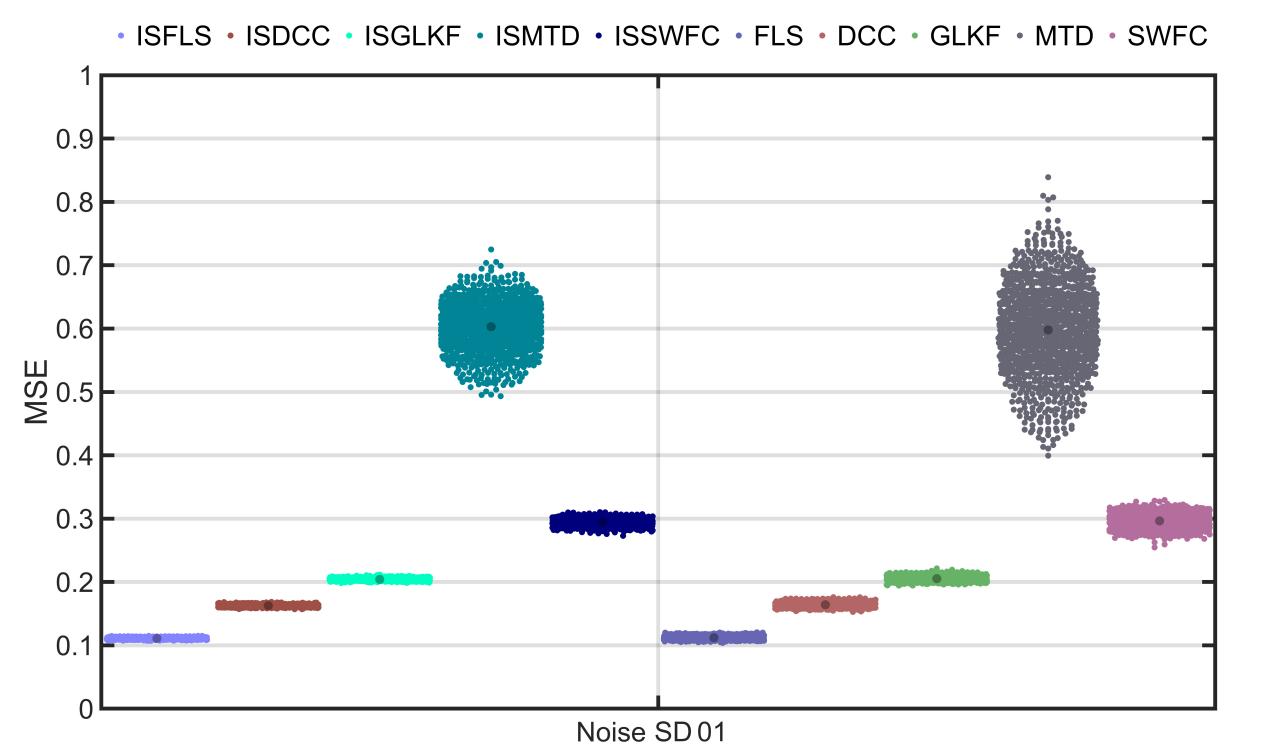

Supplementary Figure 6. MSE results of SD1-2-1. The MSE between the estimated and the actual correlation coefficients of two time series with covariance structure of Gaussian distribution from 1000 simulations are presented.

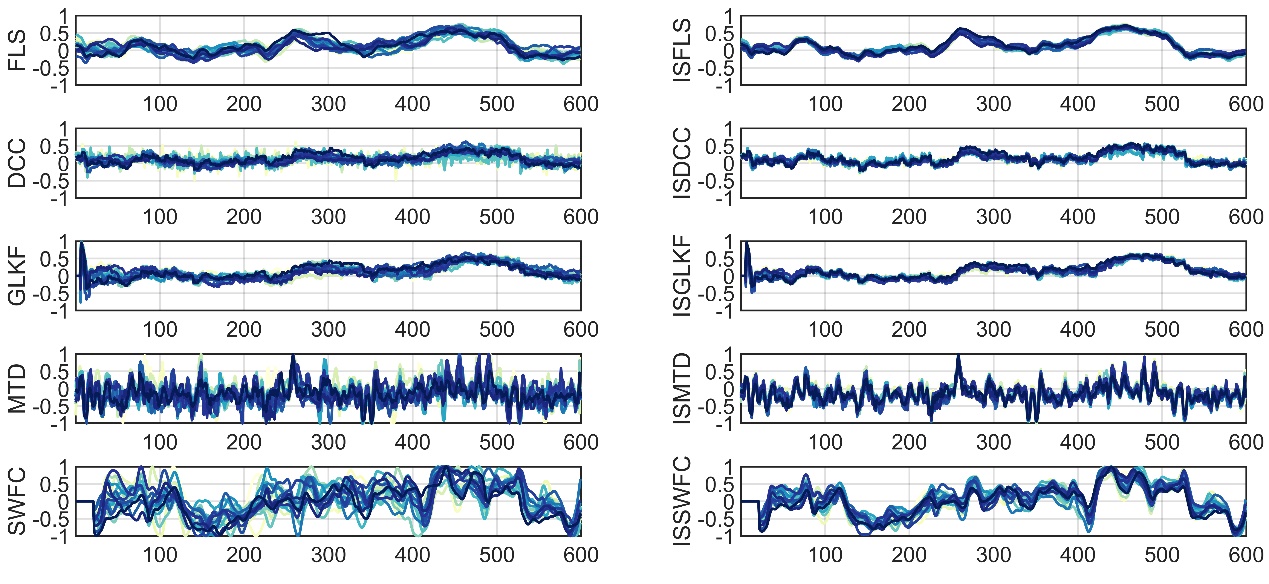

Supplementary Figure 7. Example results of SD1-2-3. The estimated correlation coefficients of two time series with covariance structure of Gaussian distribution from 20 simulations are presented.

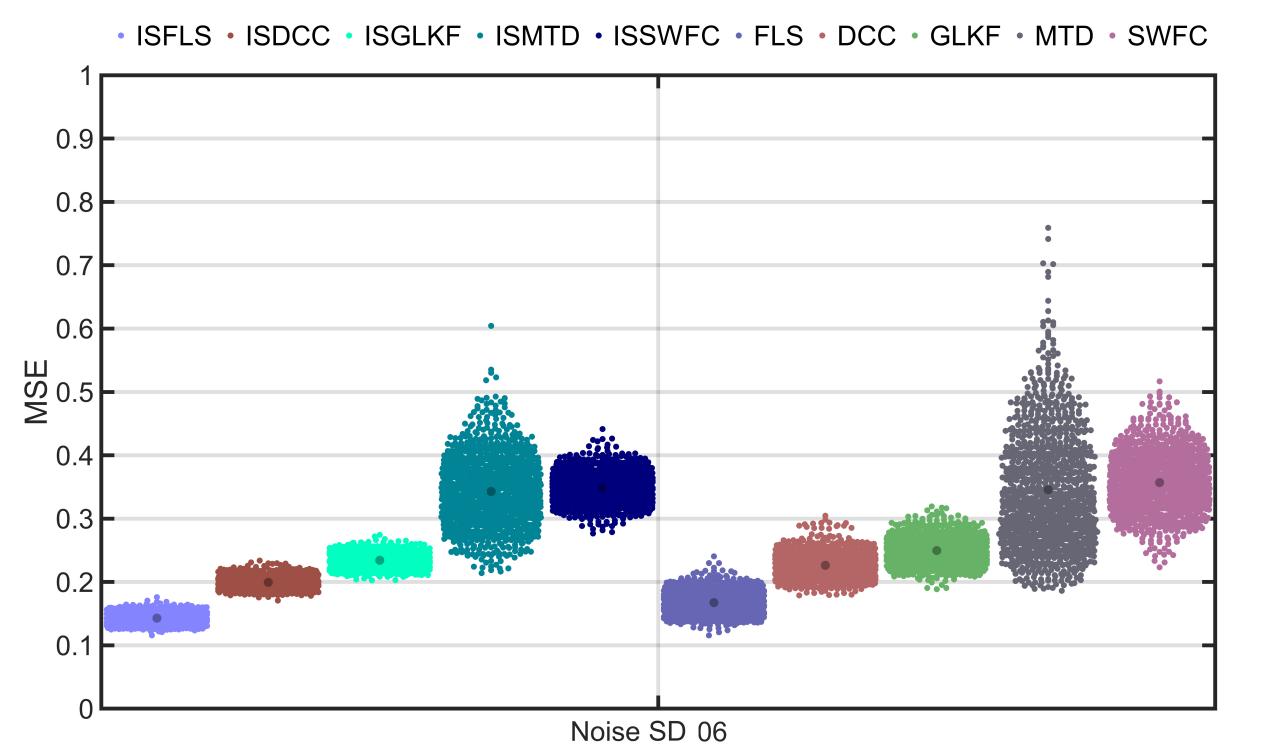

Supplementary Figure 8. MSE results of SD1-2-3. The MSE between the estimated and the actual correlation coefficients of two time series with covariance structure of Gaussian distribution from 1000 simulations are presented.

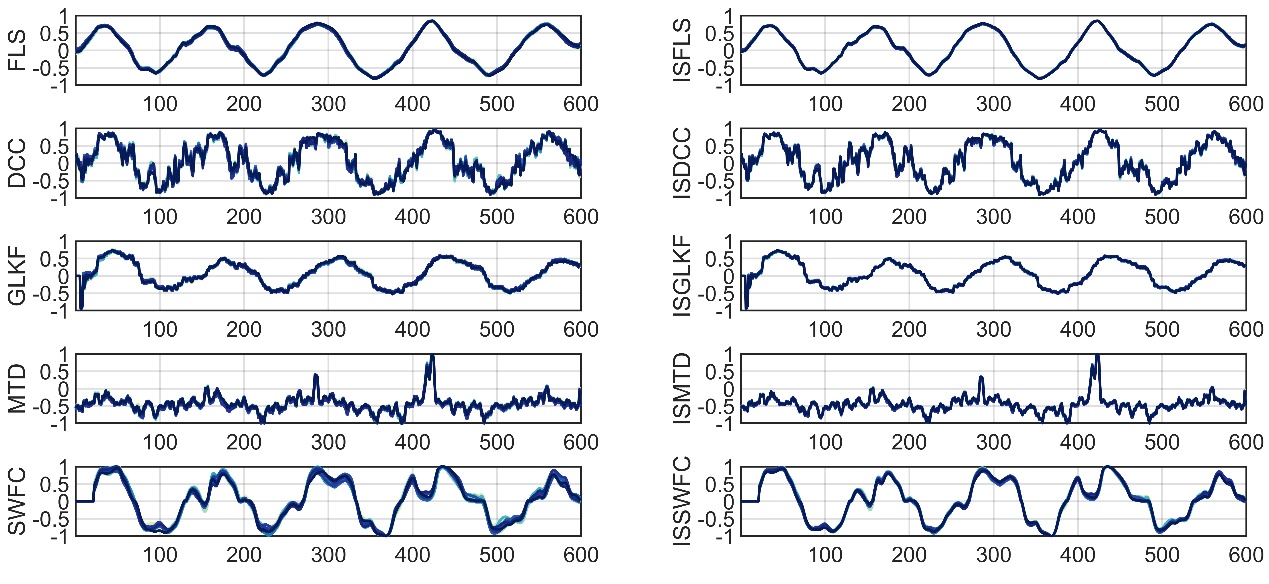

Supplementary Figure 9. Example results of SD1-3-1. The estimated correlation coefficients of two time series with covariance structure of periodic distribution from 20 simulations are presented.

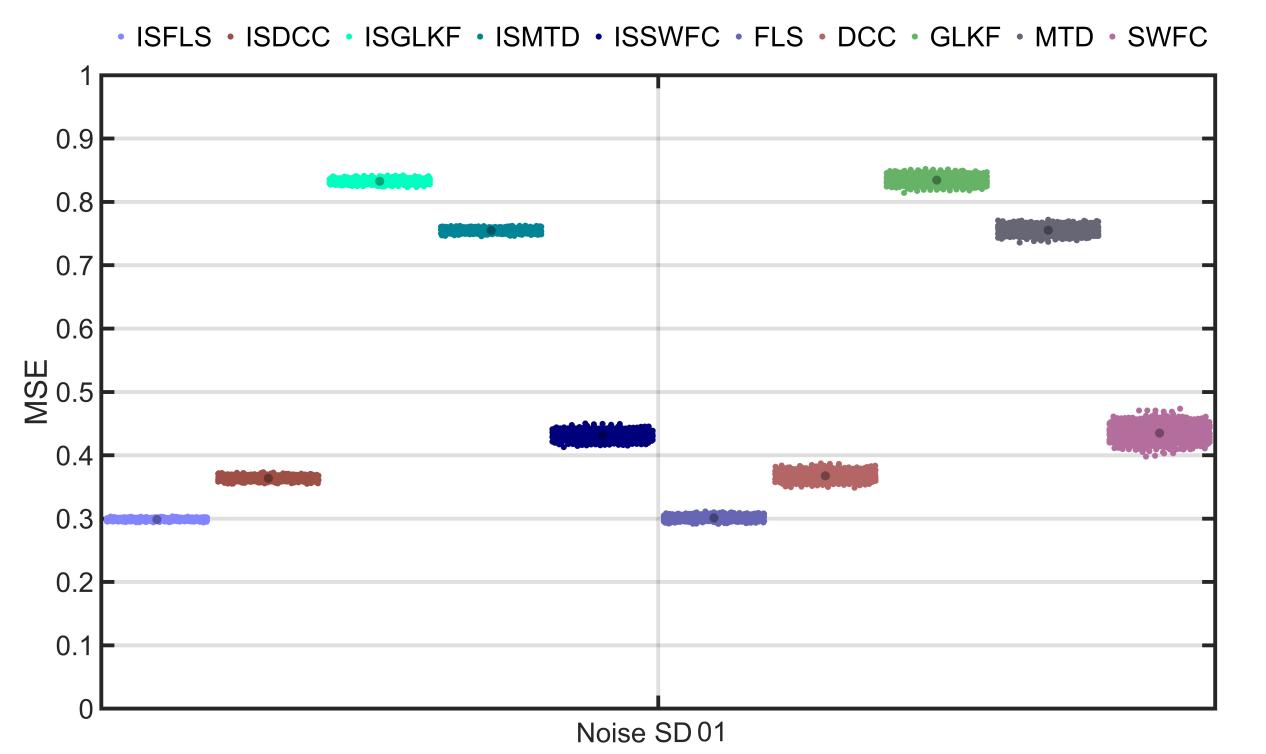

Supplementary Figure 10. MSE results of SD1-3-1. The MSE between the estimated and the actual correlation coefficients of two time series with covariance structure of periodic distribution from 1000 simulations are presented.

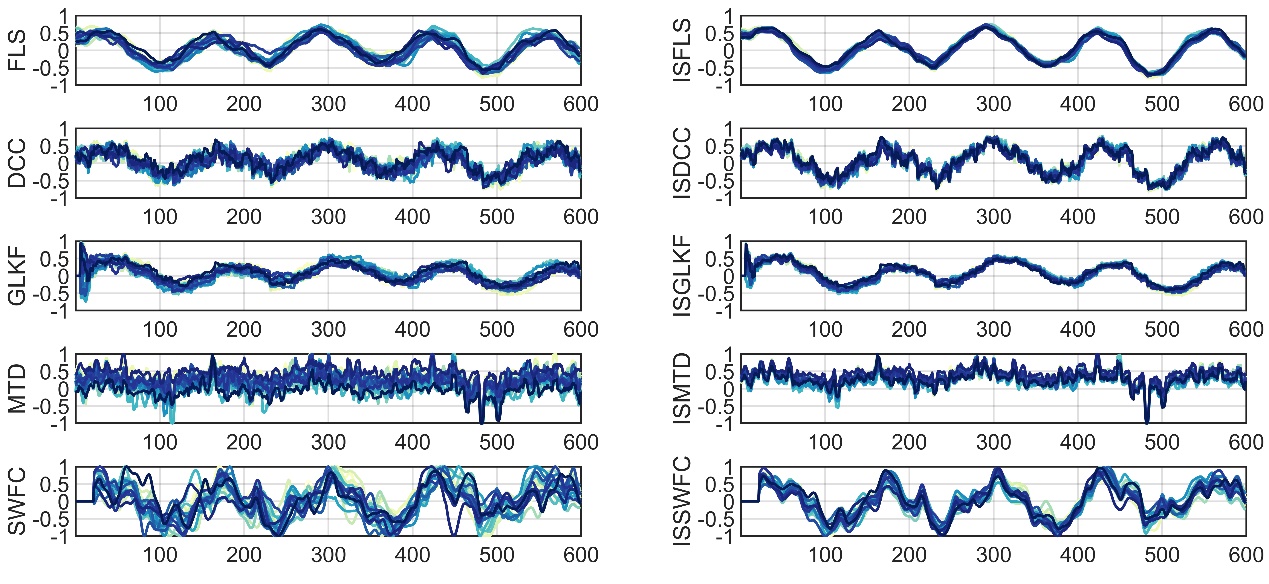

Supplementary Figure 11. Example results of SD1-3-3. The estimated correlation coefficients of two time series with covariance structure of periodic distribution from 20 simulations are presented.

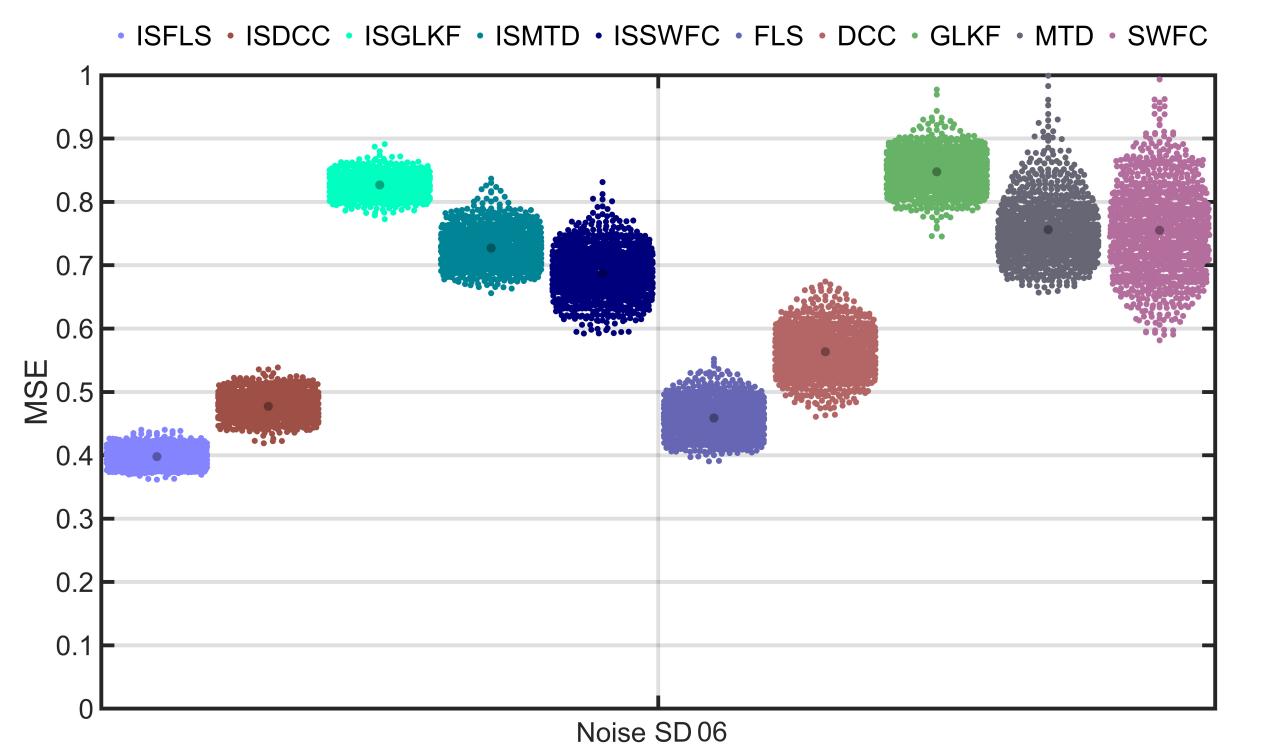

Supplementary Figure 12. MSE results of SD1-3-3. The MSE between the estimated and the actual correlation coefficients of two time series with covariance structure of periodic distribution from 1000 simulations are presented.

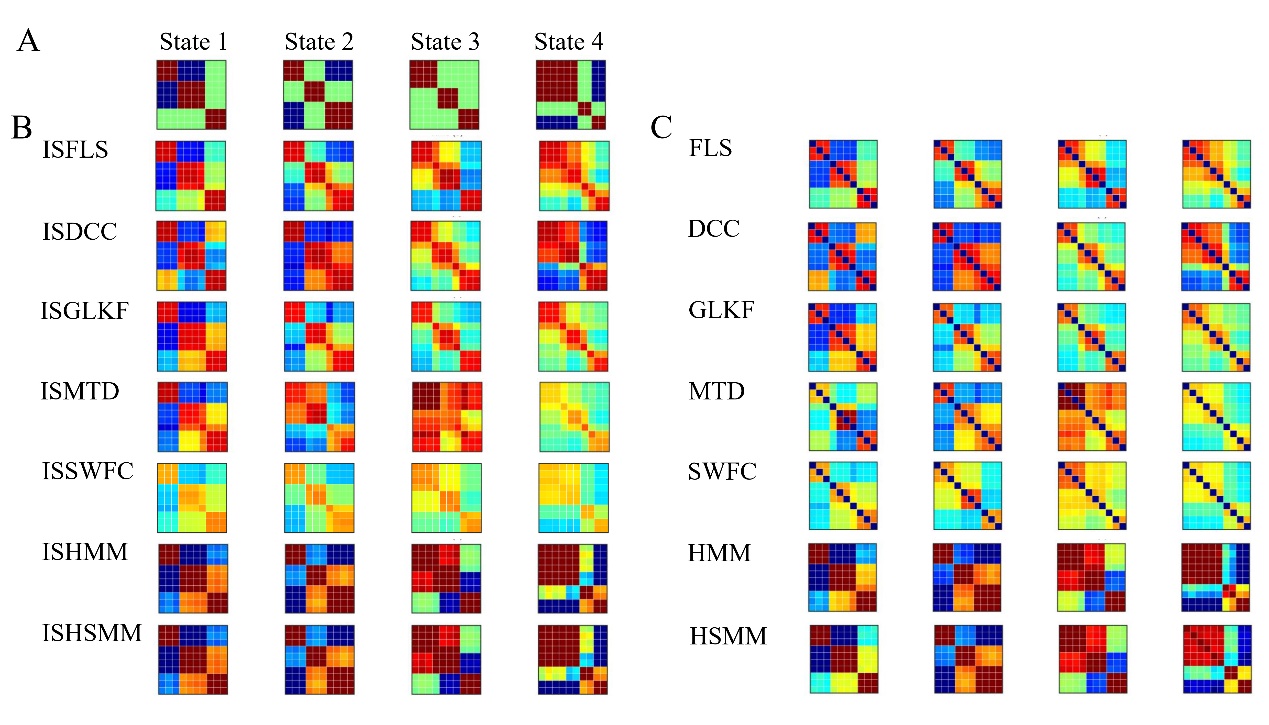

Supplementary Figure 13. The predefined and estimated states of SD2-1-1 by the 14 dFC methods.

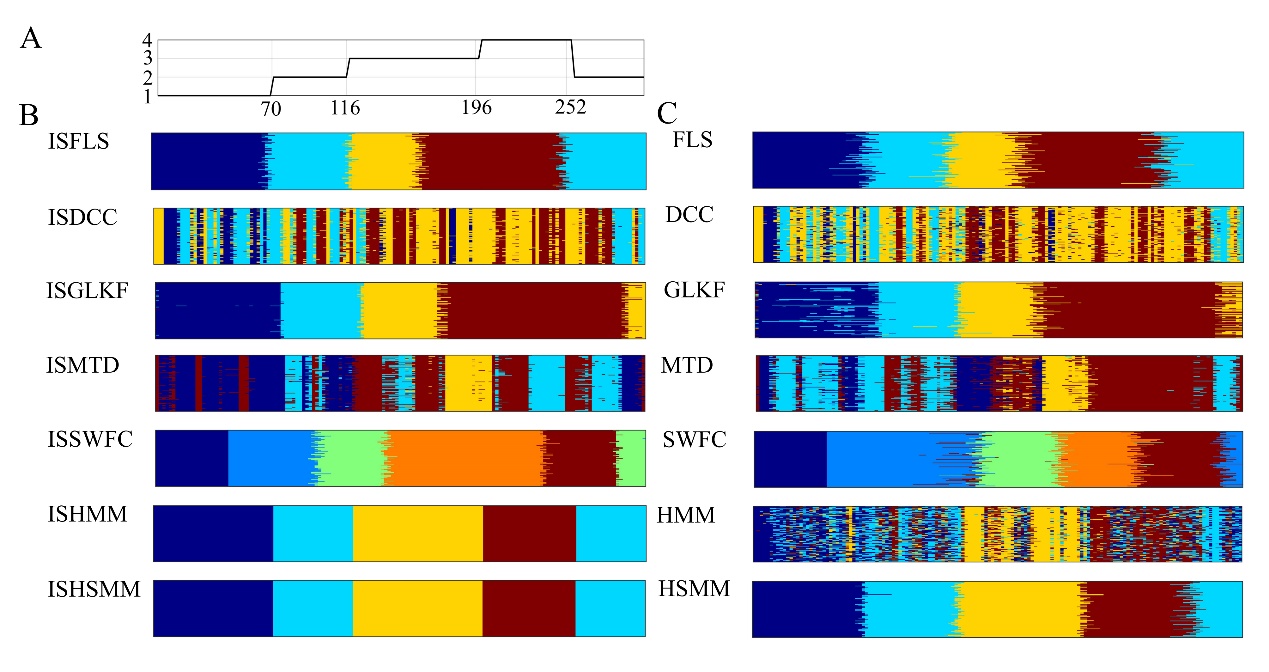

Supplementary Figure 14. The estimated state transition vectors of SD2-1-1 by the 14 dFC methods. A: The predefined state transition vector of the four states.

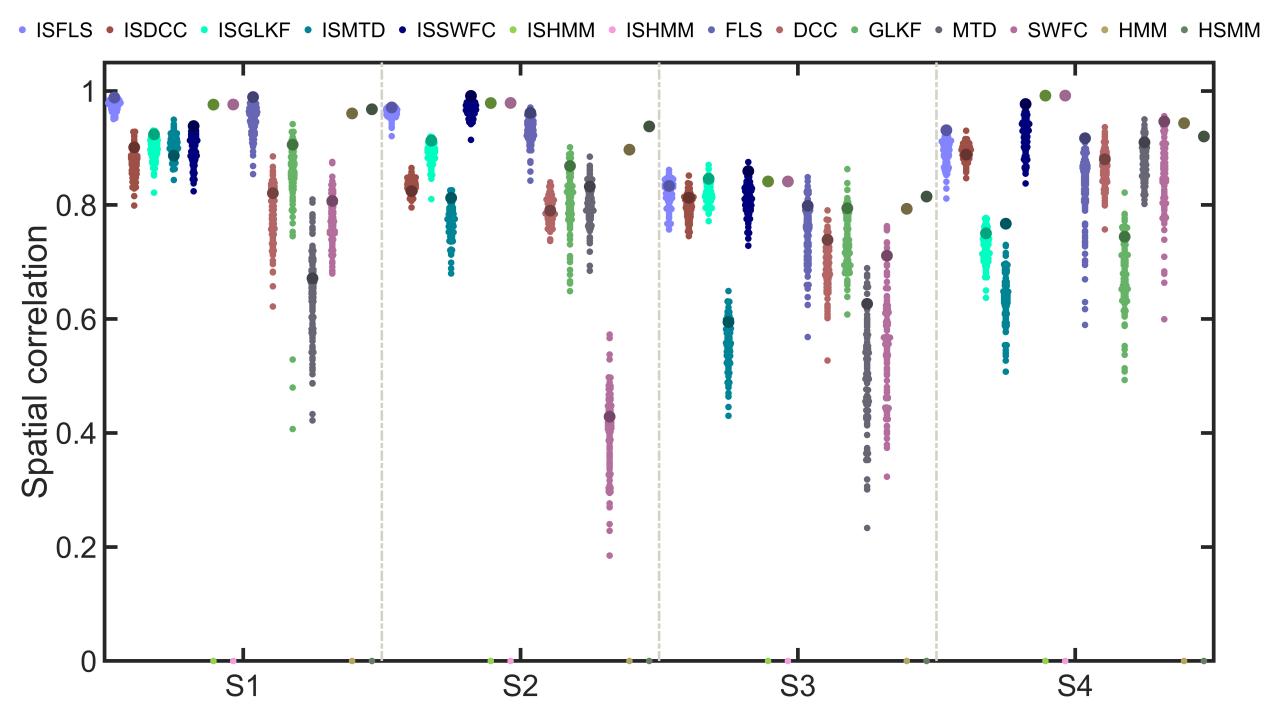

Supplementary Figure 15. Spatial correlations of SD2-1-1 between the predefined and estimated states by the 14 dFC methods.

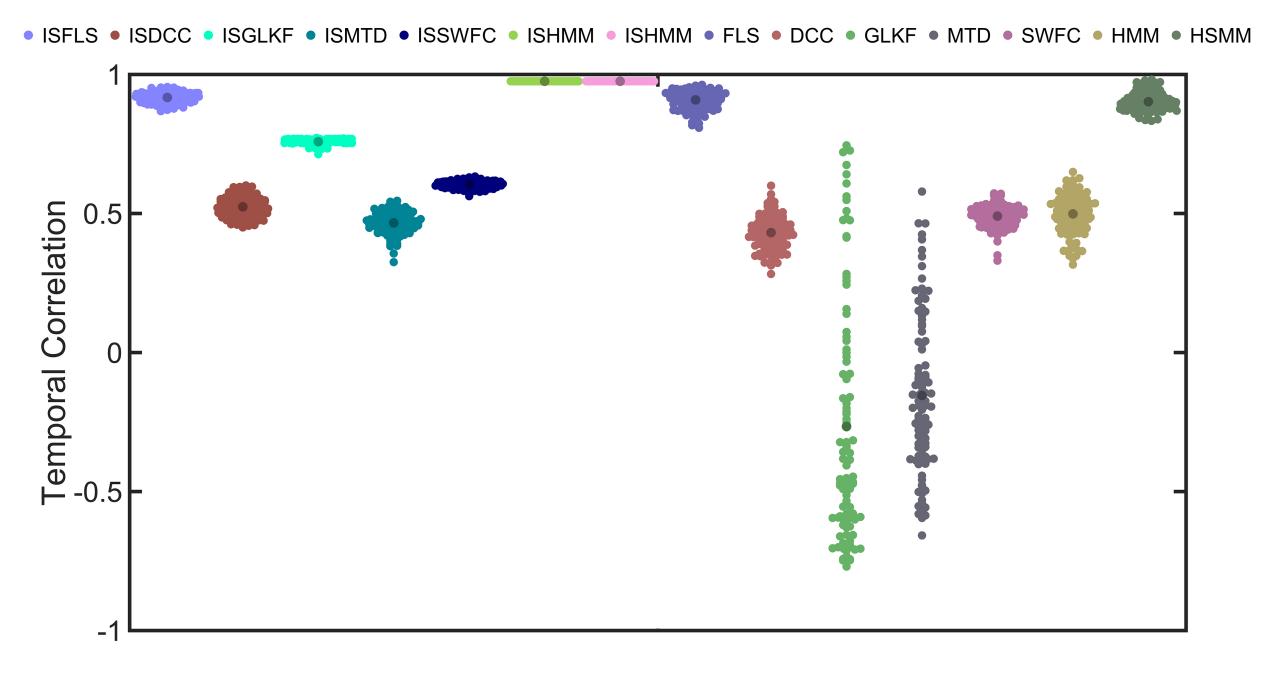

Supplementary Figure 16. Temporal correlations of SD2-1-1 between the predefined state transition vector and the estimated state transition vector by the 14 dFC methods.

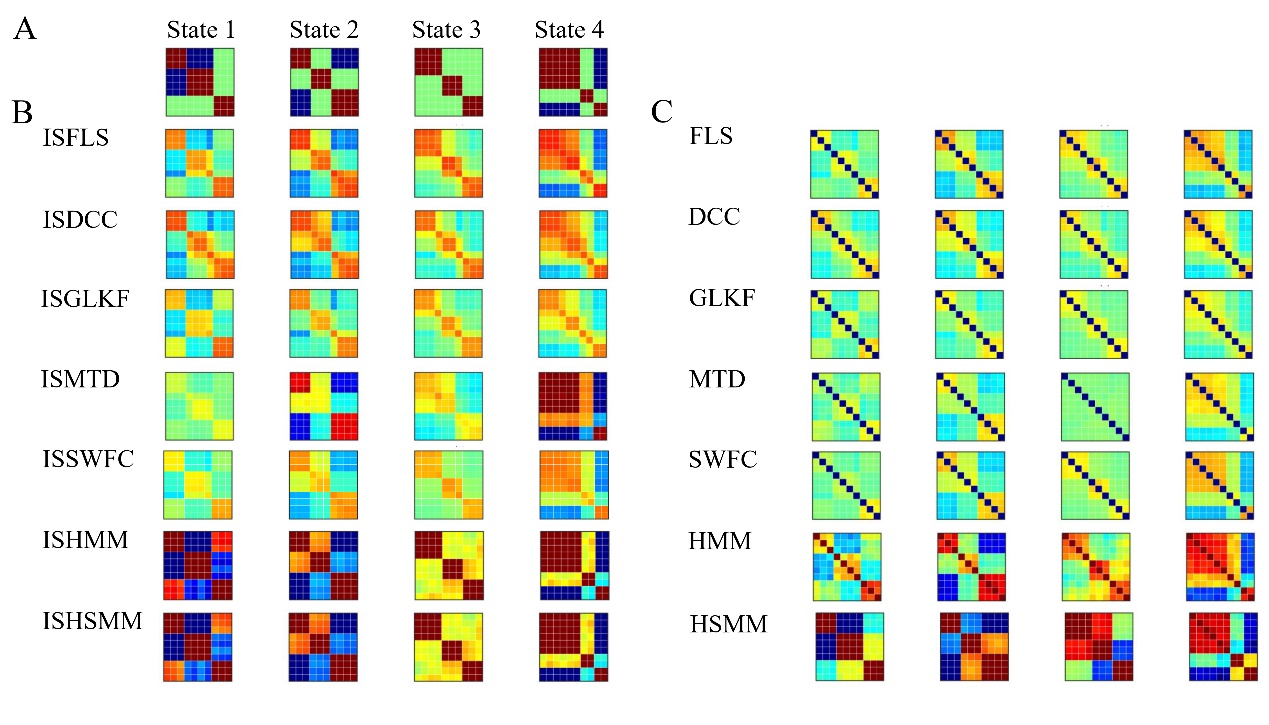

Supplementary Figure 17. The predefined and estimated states of SD2-1-3 by the 14 dFC methods.

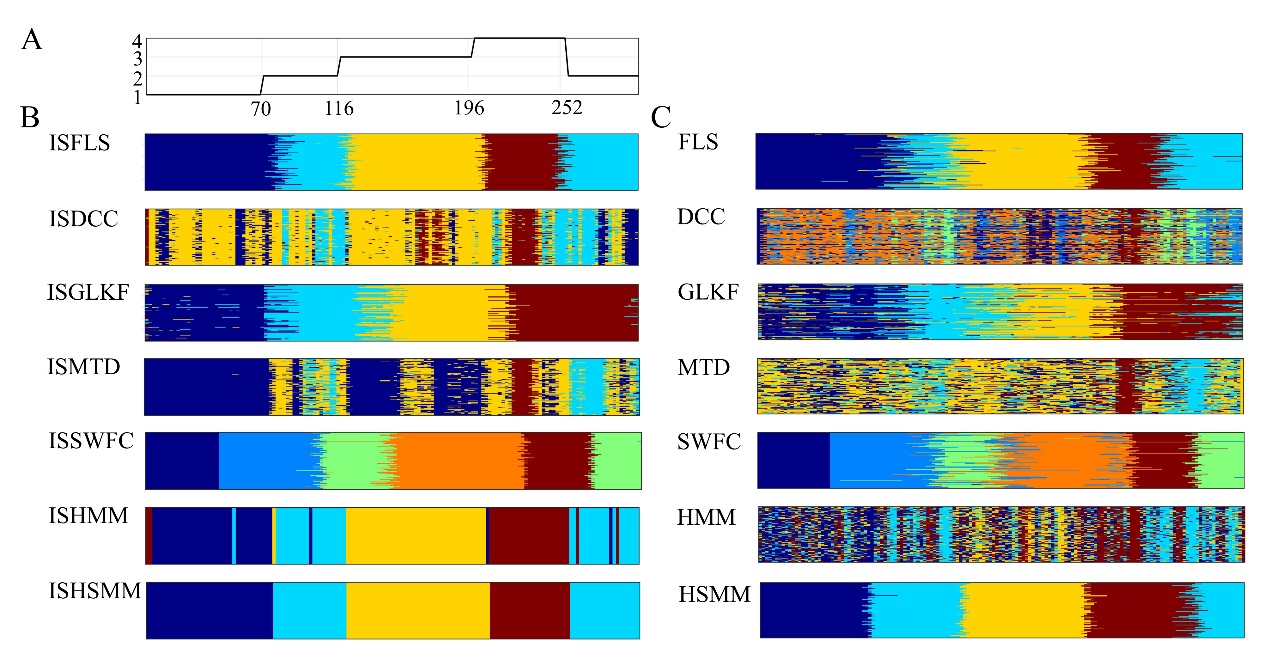

Supplementary Figure 18. The estimated state transition vectors of SD2-1-3 by the 14 dFC methods. A: The predefined state transition vector of the four states. B & C: the estimated state transition vectors.

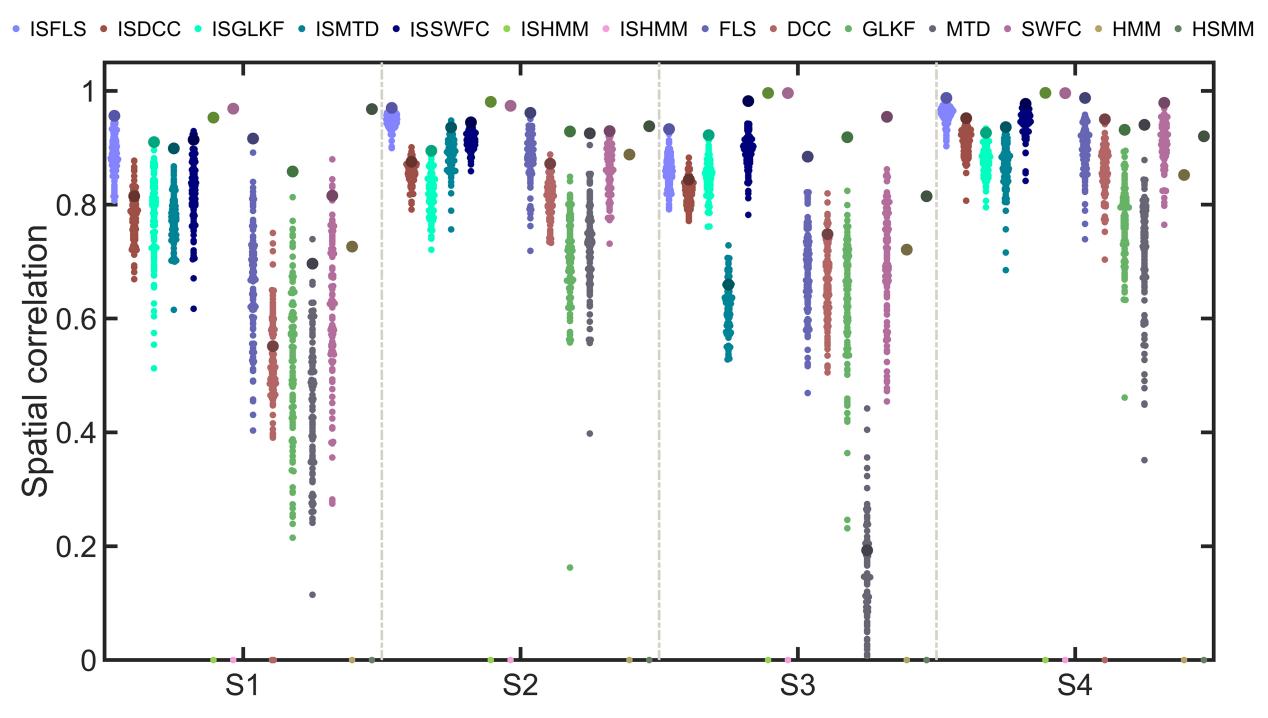

Supplementary Figure 19. Spatial correlations of SD2-1-3 between the predefined states and the estimated states by the 14 dFC methods.

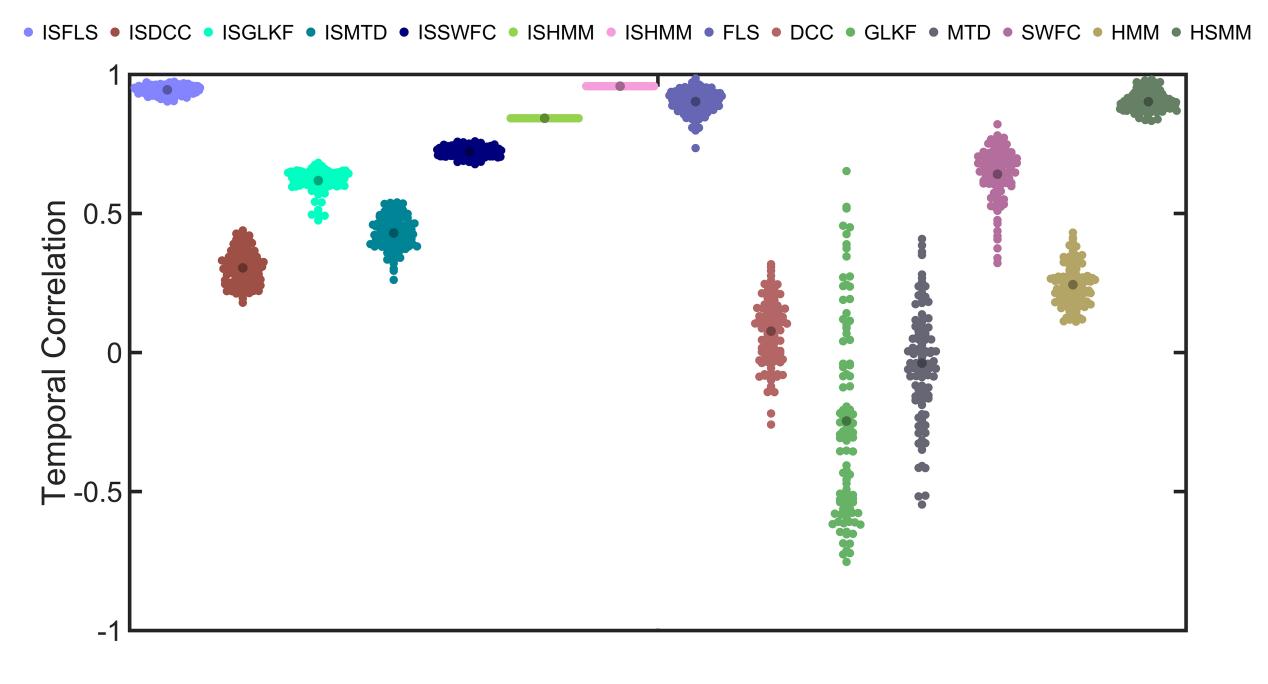

Supplementary Figure 20. Temporal correlations of SD2-1-3 between the predefined state transition vector and the estimated state transition vector by the 14 dFC methods.

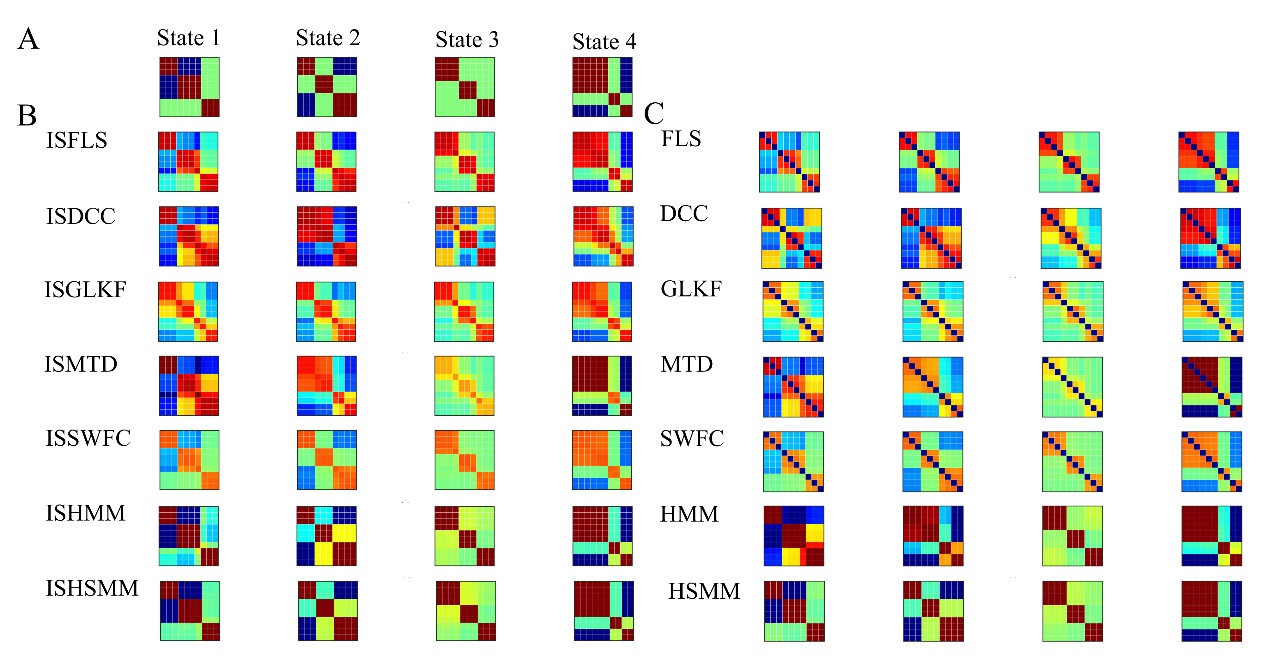

Supplementary Figure 21. The predefined and estimated states of SD2-2-1 were determined using the 14 dFC methods.

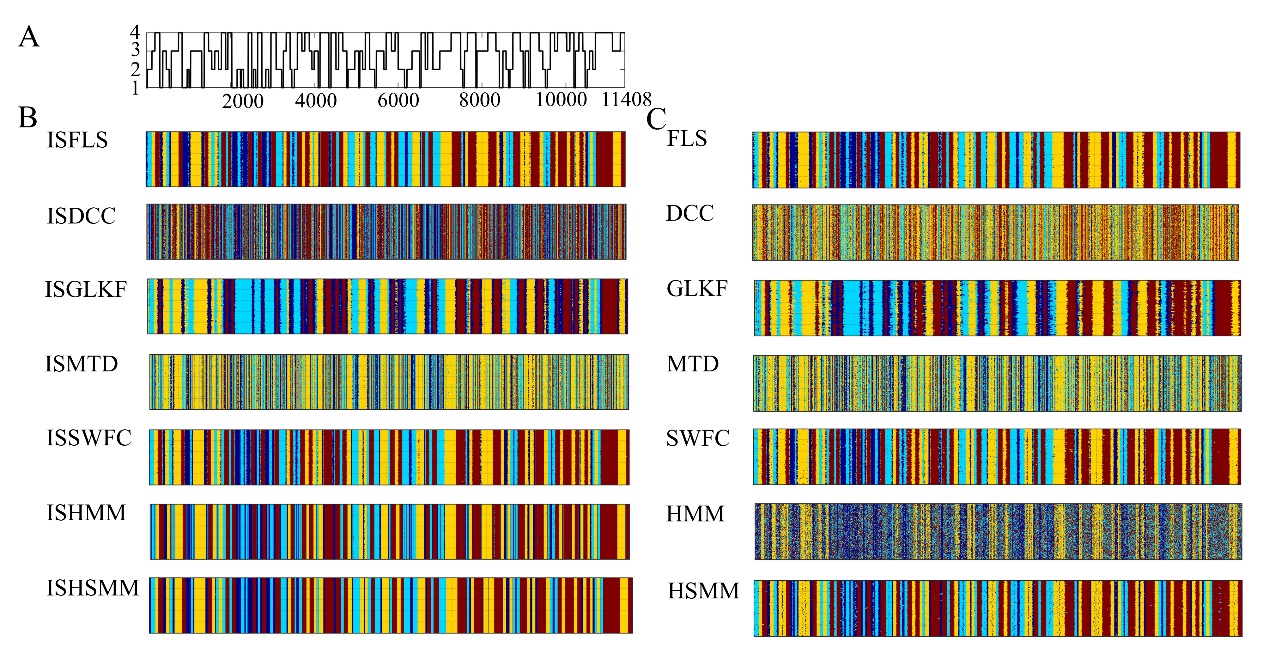

Supplementary Figure 22. The estimated state transition vectors of SD2-2-1 by the 14 dFC methods. A: The predefined state transition vector of the four states. B & C: the estimated state transition vectors.

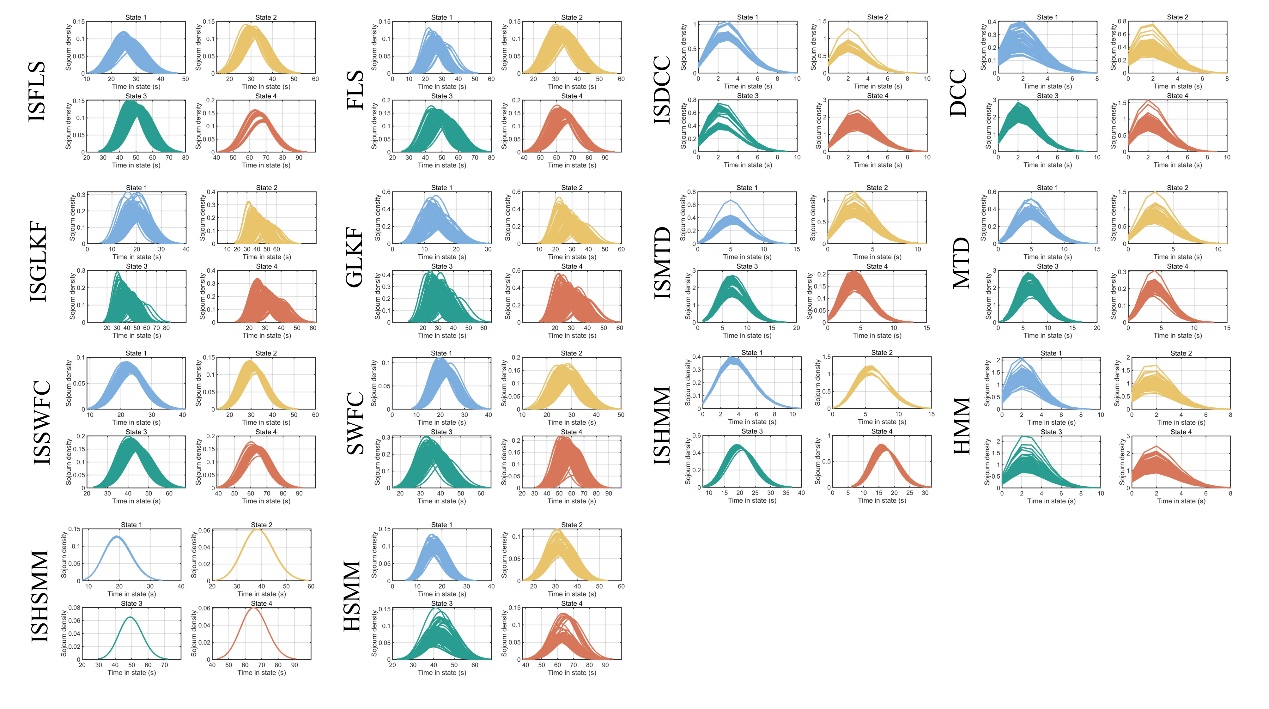

Supplementary Figure 23. The fitted distribution of state sojourn time of SD2-2-1 by the 14 dFC methods.

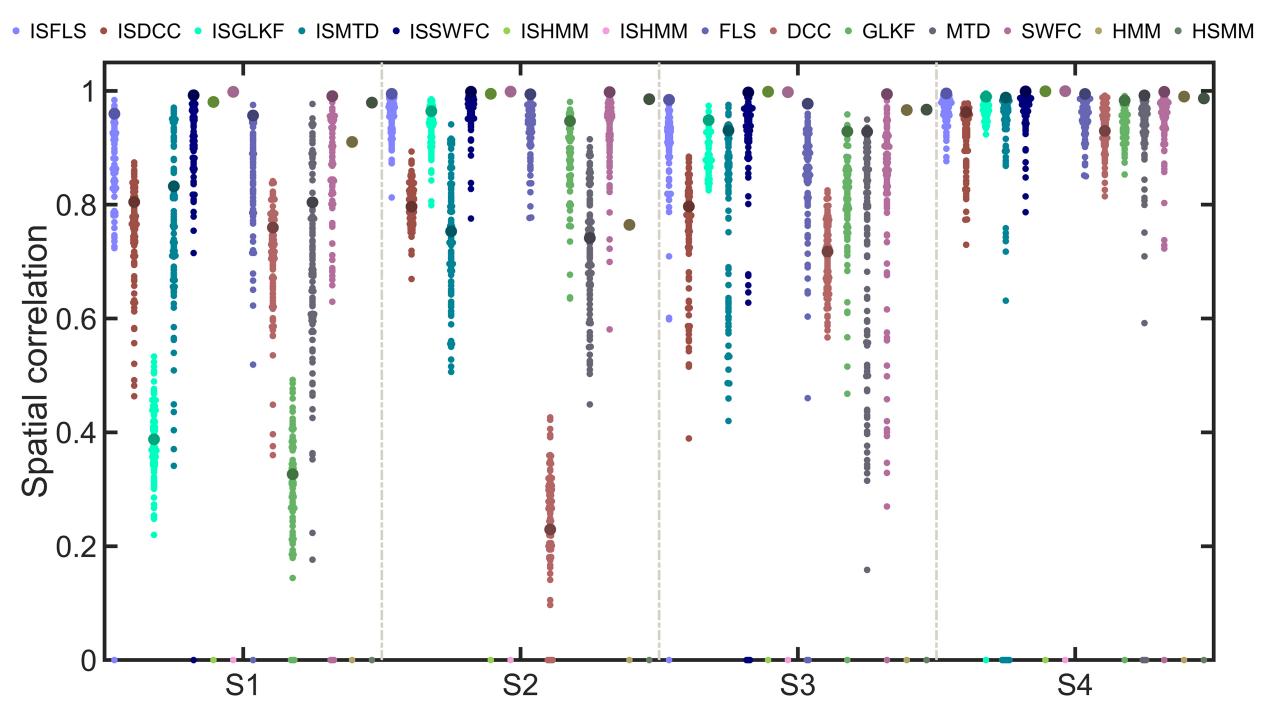

Supplementary Figure 24. Spatial correlations of SD2-2-1 between the predefined states and the estimated states by the 14 dFC methods.

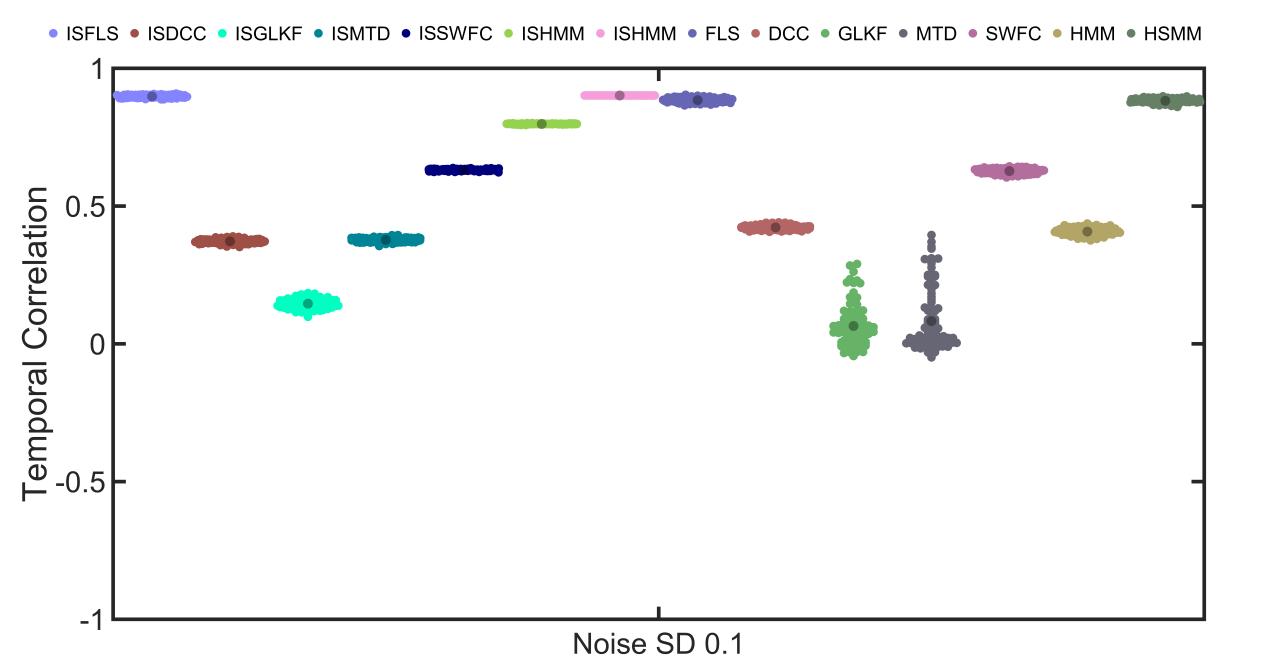

Supplementary Figure 25. Temporal correlations of SD2-2-1 between the predefined state transition vector and the estimated state transition vector by the 14 dFC methods.

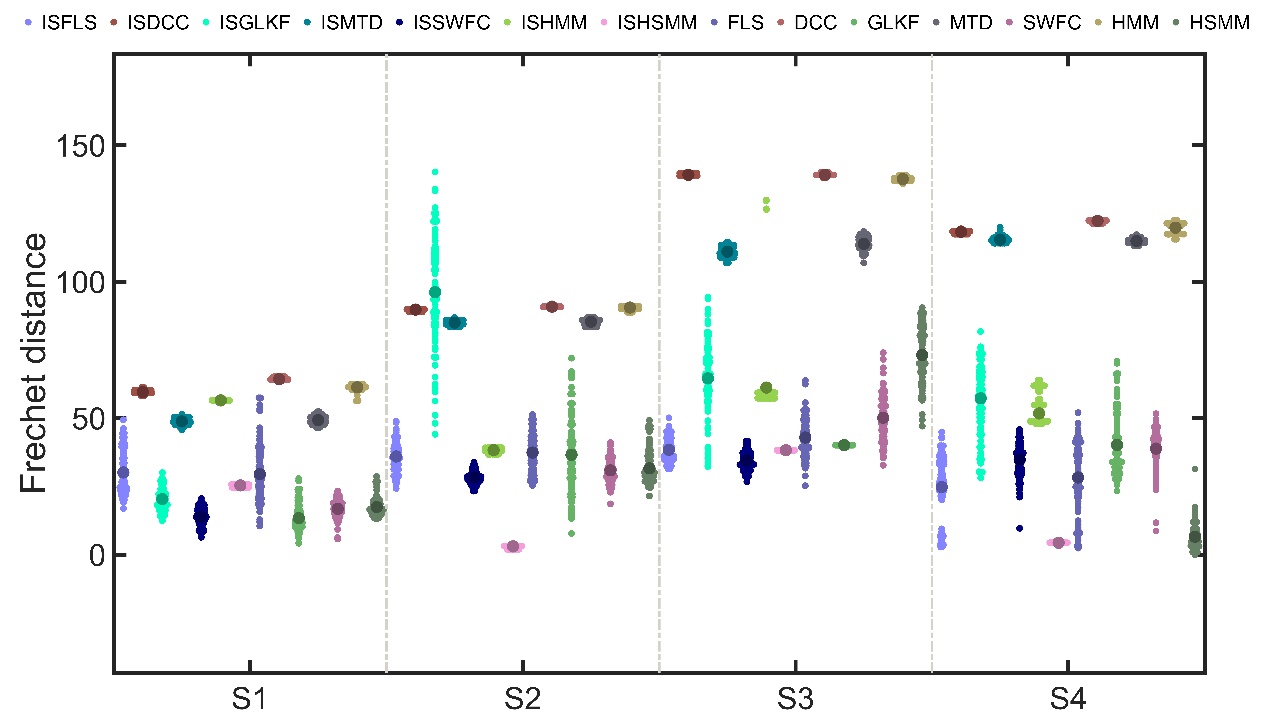

Supplementary Figure 26. The Frechet distance between the curve of predefined state sojourn time distributions and the curve of estimated state sojourn time distributions for SD2-2-1, calculated using the 14 dFC methods.

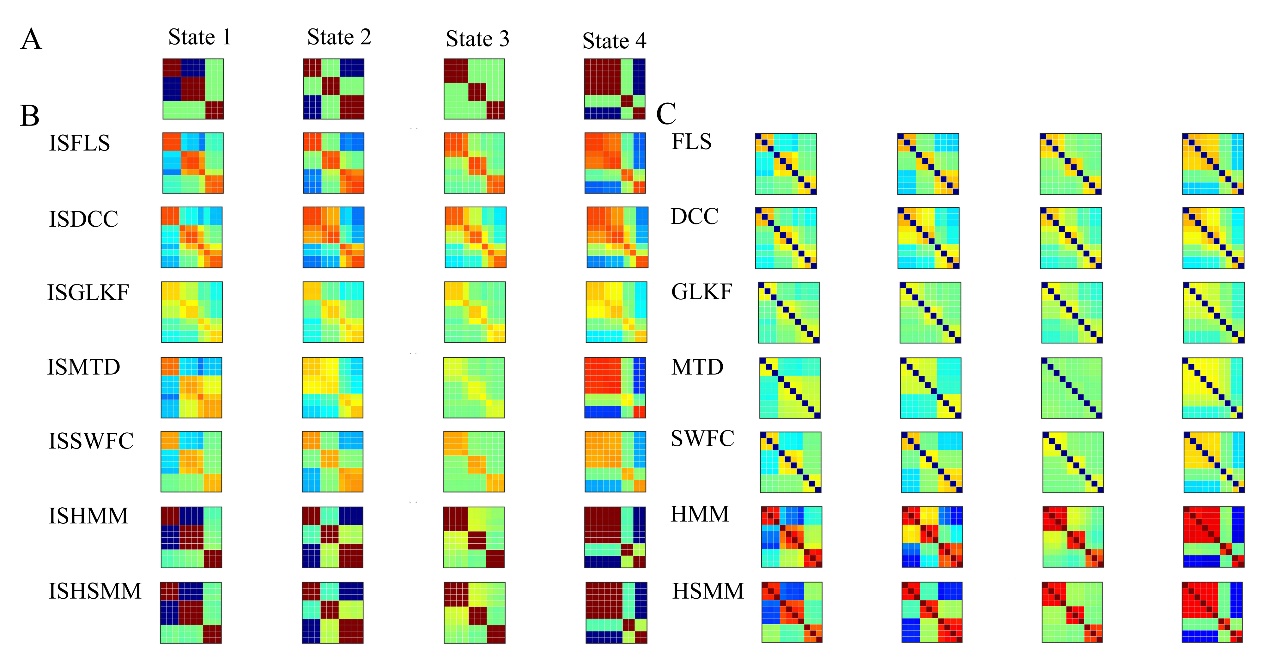

Supplementary Figure 27. The predefined and estimated states of SD2-2-3 by the 14 dFC methods.

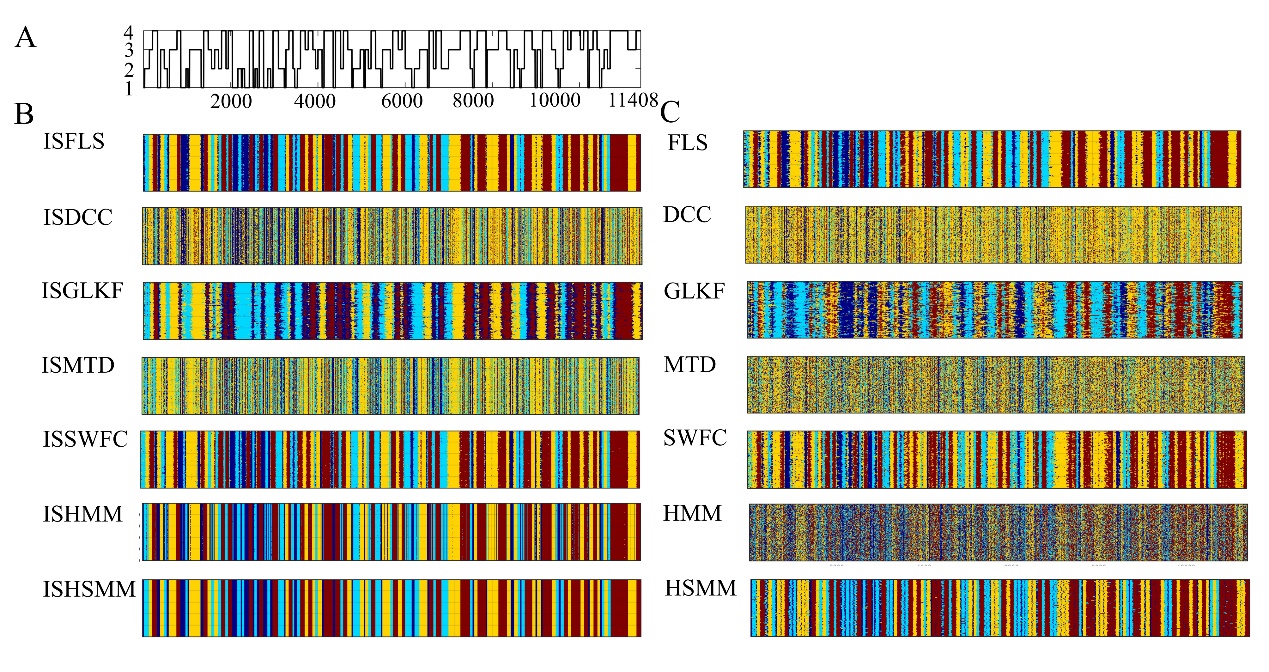

Supplementary Figure 28. The estimated state transition vectors of SD2-2-3 by the 14 dFC methods. A: The predefined state transition vector of the four states. B & C: the estimated state transition vectors.

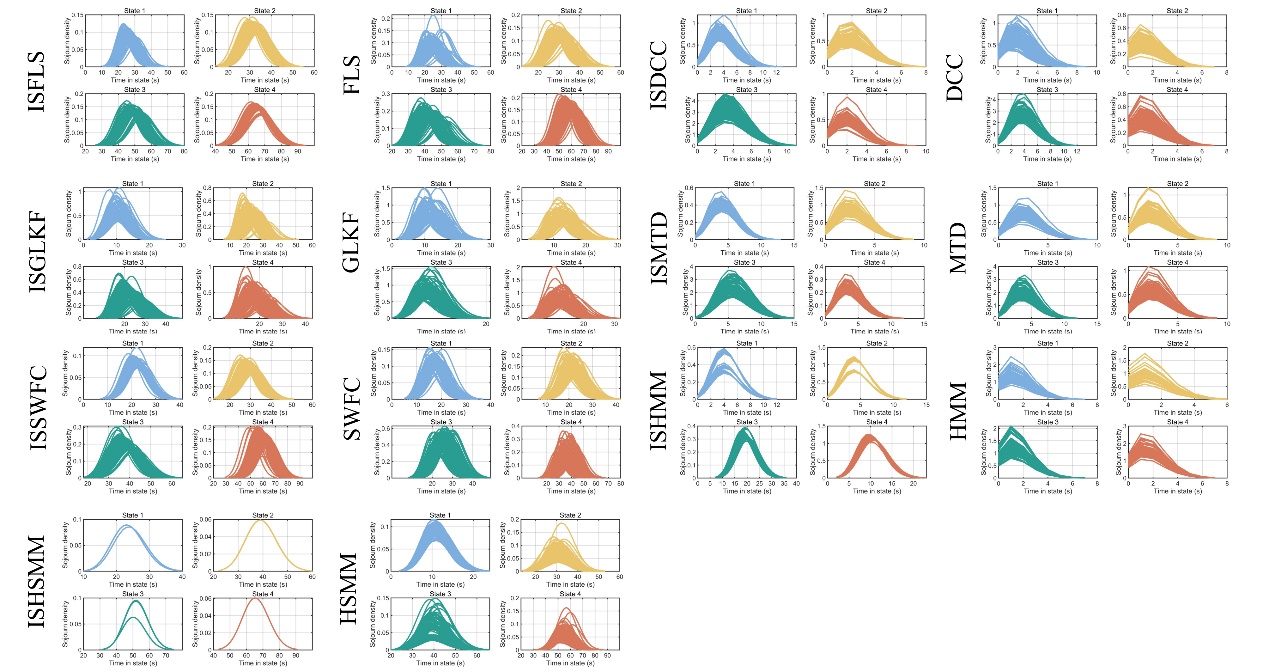

Supplementary Figure 29. The fitted distribution of state sojourn time of SD2-2-3 by the 14 dFC methods.

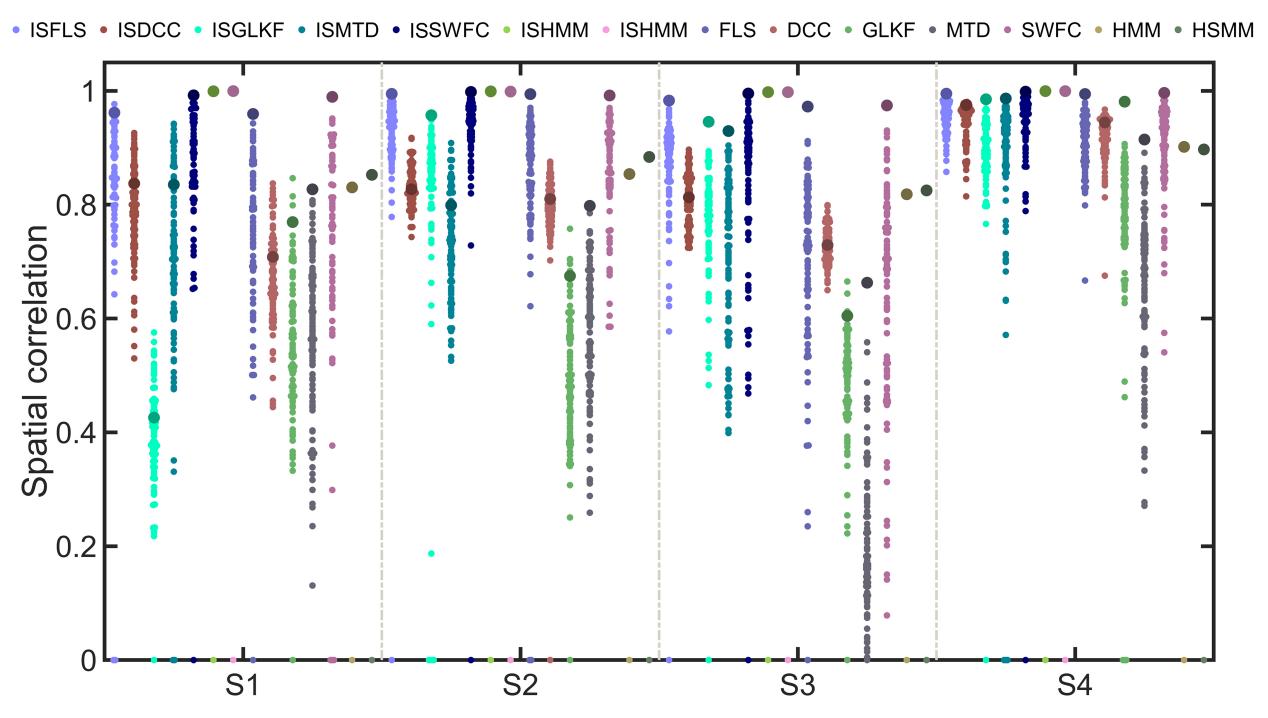

Supplementary Figure 30. Spatial correlations of SD2-2-3 between the predefined states and the estimated states by the 14 dFC methods.

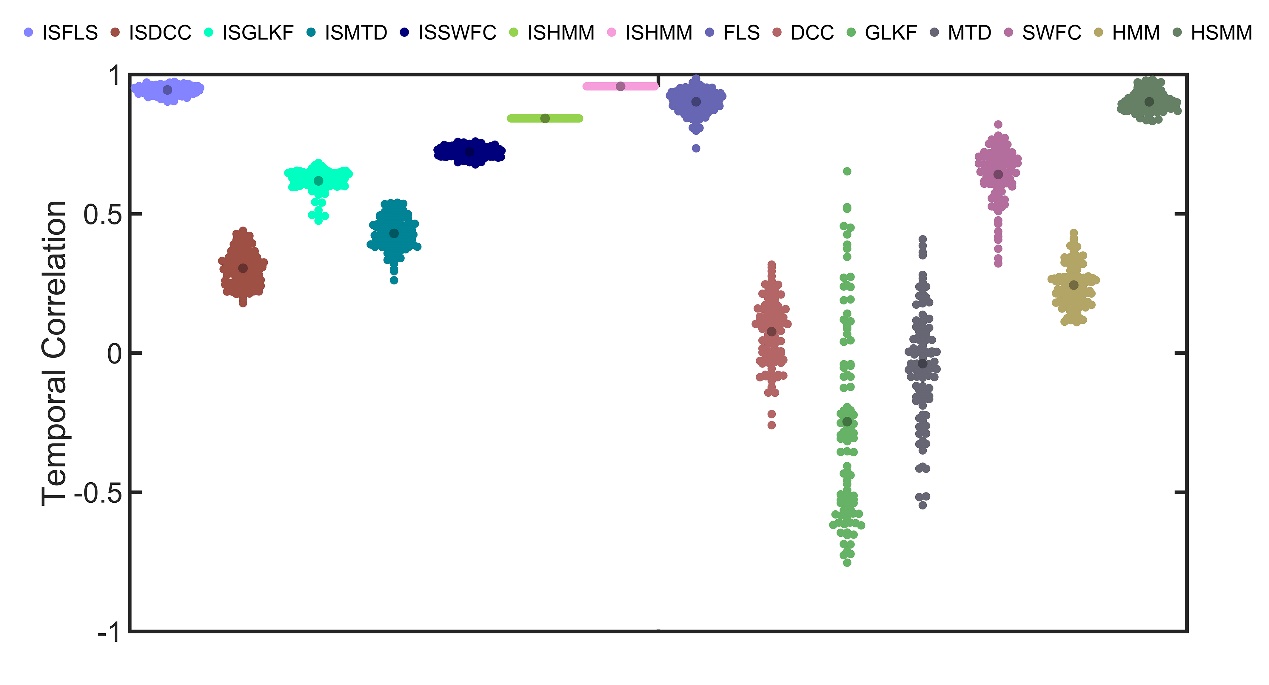

Supplementary Figure 31. Temporal correlations of SD2-2-3 between the predefined state transition vector and the estimated state transition vector by the 14 dFC methods.

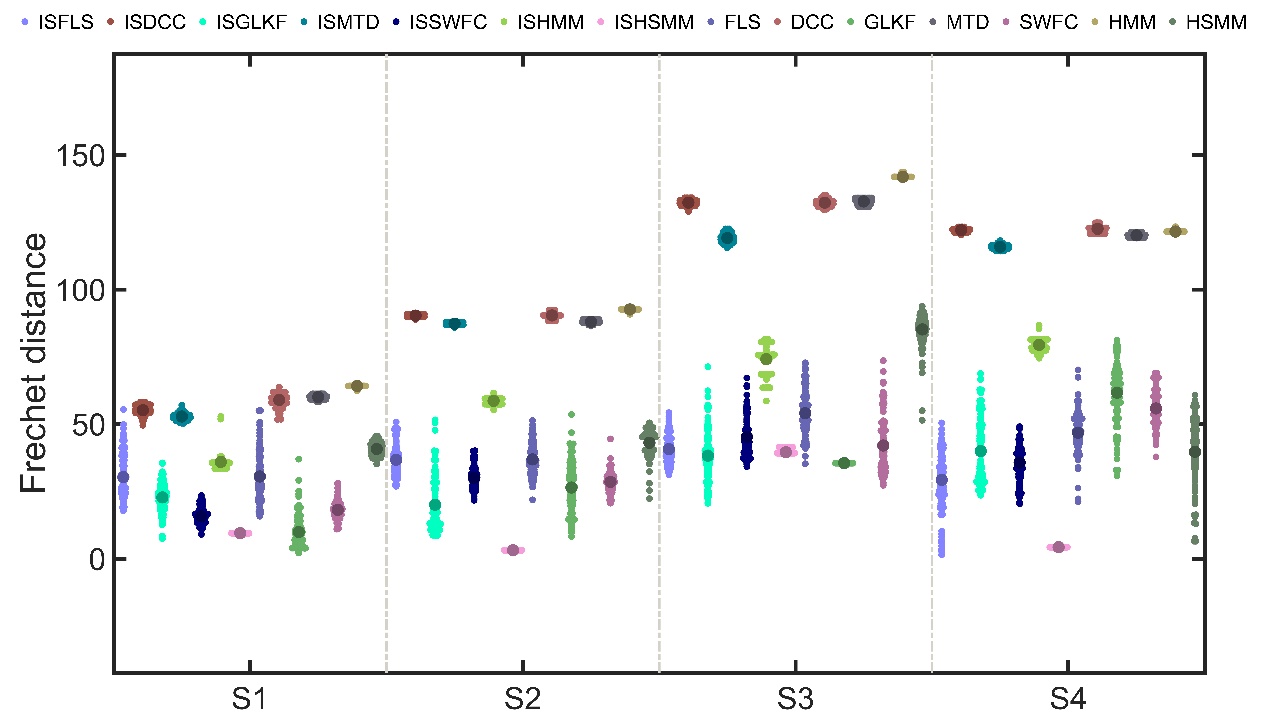

Supplementary Figure 32. The Frechet distance between the curve of predefined state sojourn time distributions and the curve of estimated state sojourn time distributions for SD2-2-2, calculated using the 14 dFC methods.

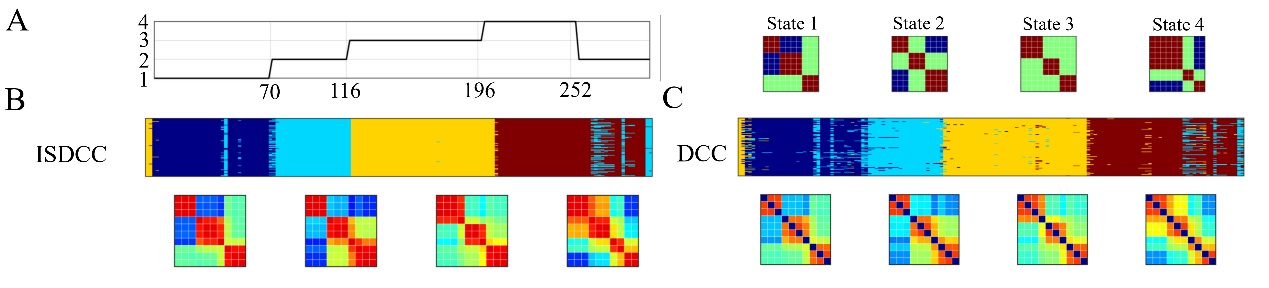

Supplementary Figure 33. The true and estimated results of ISDCC and DCC for a new SD2-1-2 dataset, without HRF convolution (SD2-1-2-NoHRF). Ten time series with four connectivity states and predefined state sojourn times were simulated 100 times. A: the four connectivity states and predefined state sojourn times. B&C: the estimated connectivity states and state transition vectors by ISDCC and DCC.

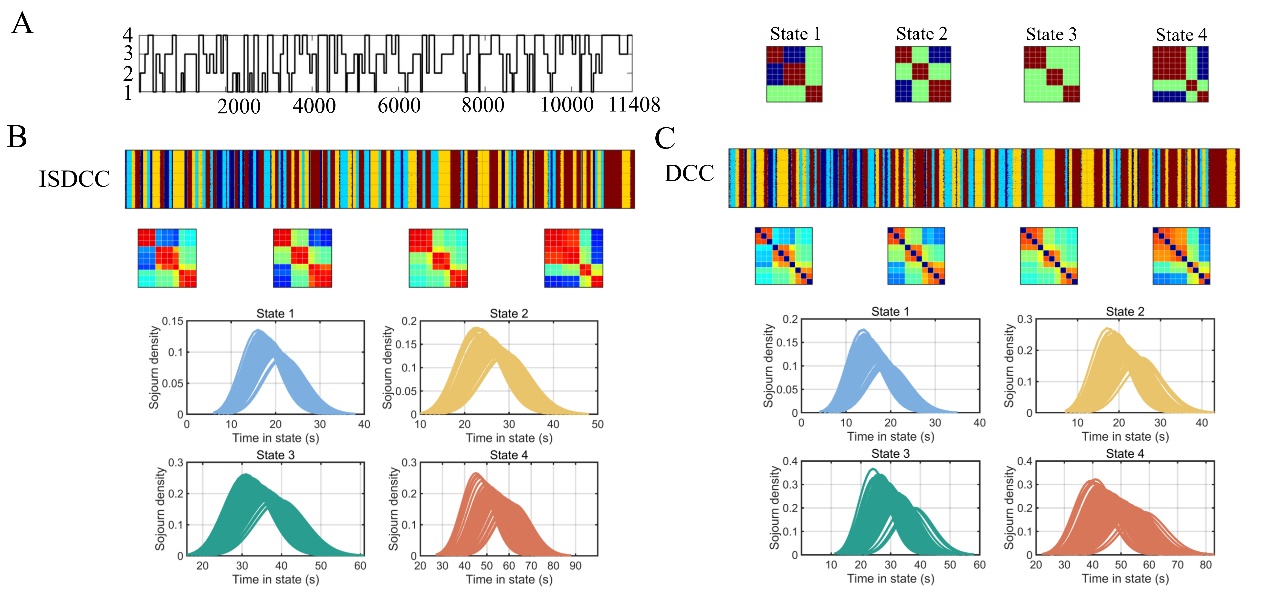

Supplementary Figure 34. The true and estimated results of ISDCC and DCC for a new SD2-2-2 dataset, without HRF convolution (SD2-2-2-NoHRF). Ten time series with four connectivity states and predefined state sojourn times were simulated 100 times. The state sojourn times were A: the four connectivity states and predefined state sojourn times. B&C: the estimated connectivity states and state transition vectors by ISDCC and DCC.

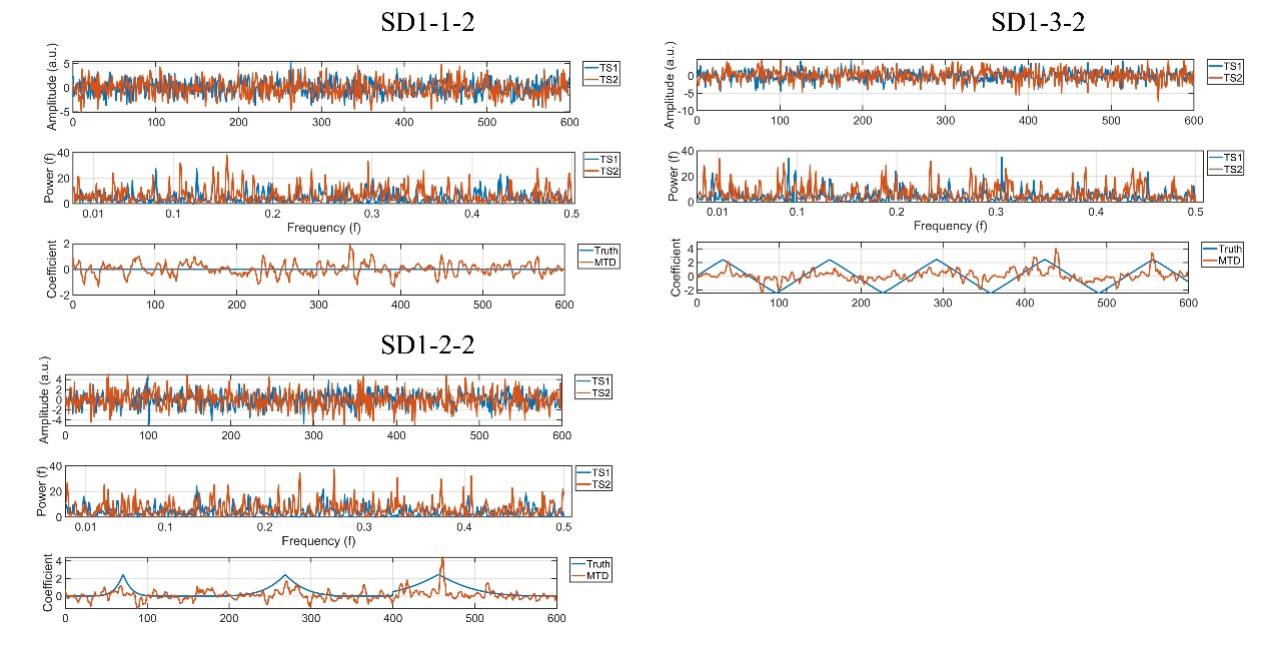

Supplementary Figure 35. Three cases of temporal and frequency characteristics, and the estimated coefficients by MTD of two time series in SD1, with covariance structures of zeros throughout the time series (SD1-1-2), Gaussian distribution (SD1-2-2), and periodic distribution (SD1-3-2), respectively. For each data, the standard deviations of noise signals were 0.3. The power for each time series were calculated using fast Fourier transform.

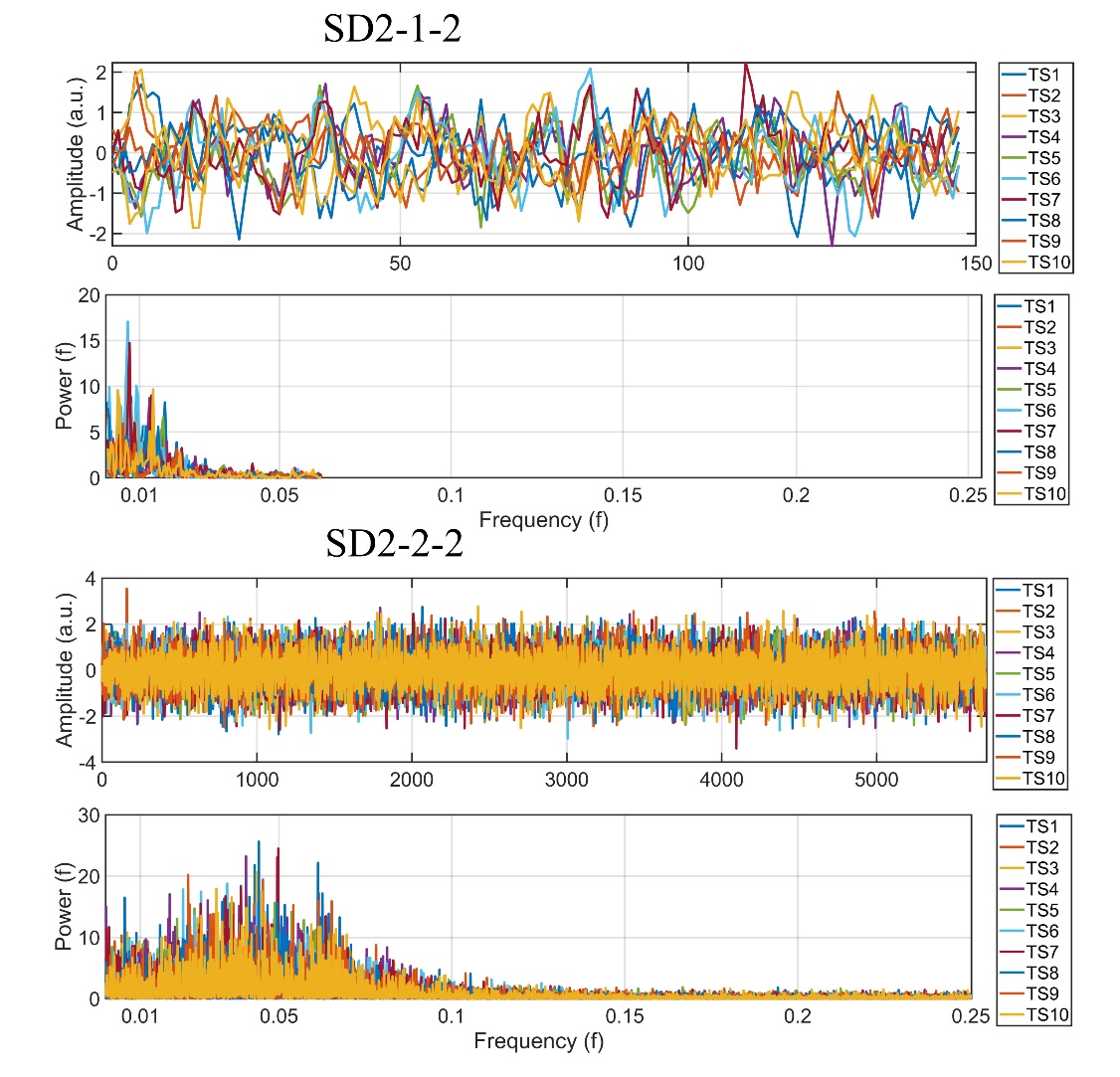

Supplementary Figure 36. Two cases of temporal and frequency characteristics of 10 time series in SD2, with covariance structures of fixed sojourn times (SD2-1-2), and Poisson distribution (SD2-2-2), respectively. For each data, the standard deviations of noise signals were 0.3. The power for each time series were calculated using fast Fourier transform.
